# ConvergeCELL: An end-to-end platform from patient transcriptomics to therapeutic hypotheses

**DOI:** 10.64898/2026.05.07.723555

**Authors:** Noam Shahar, Danielle Miller, Meishar Shahoha, Guy Lurie, Iddo Weiner

**Affiliations:** Converge Bio Ltd

## Abstract

Translating transcriptomic data into therapeutic hypotheses remains fragmented and labor-intensive. Here we present ConvergeCELL, a platform combining a patient representation model trained on over 20 million cells across 4,479 patients, an interpretability framework for gene discovery, and a large language model-driven workflow that classifies candidates along an evidence hierarchy and constructs mechanism-of-action hypotheses. Validated on held-out cohorts spanning lupus, multiple myeloma, and sepsis across single-cell and bulk modalities, ConvergeCELL recovers known disease-associated genes at or above differential expression, machine-learning, and patient-level foundation model (PaSCient) baselines. The advantage is most pronounced for clinically validated, disease-specific drug targets: ConvergeCELL ranks TNFSF13B (Belimumab; lupus), TNFRSF17/BCMA (Belantamab; myeloma), and CXCR4 (Plerixafor; myeloma) within the top 0.3% of its gene rankings - significantly outcompeting alternative approaches. ConvergeCELL delivers an end-to-end translational workflow with state-of-the-art performance on both disease-associated gene recovery and patient-level disease classification. The pretrained ConvergeCELL patient representation model and bulk distillation module are publicly available on Hugging Face (huggingface.co/ConvergeBio/virtual-cell-patient) under the Apache 2.0 license.

## Main

The promise of translational research rests on our ability to convert molecular measurements into actionable biological insight. Single-cell RNA sequencing (scRNA-seq) has transformed our understanding of cellular heterogeneity in health and disease, enabling the identification of rare cell populations, disease-specific cellular states, and developmental trajectories^1,2^. Yet a fundamental gap persists between the resolution of modern transcriptomic technologies and the actionability of their outputs. While scRNA-seq datasets continue to accumulate - now exceeding hundreds of millions of profiled cells across thousands of patient cohorts - the process of translating these data into therapeutic hypotheses remains largely manual, requiring substantial expert curation and complex multi-step analyses specific to each disease context^3^.

Common use cases for such analyses include identifying biomarkers and discovering targets for therapeutic intervention. Biomarkers are factors associated with patient outcomes, used for clinical trial design and treatment response monitoring. Therapeutic targets require evidence of causality beyond association. Despite their distinct roles, identifying both biomarkers and therapeutic targets aim to link gene expression to disease-relevant phenotypes. To maintain precision, we define an evidence hierarchy for such computationally identified genes: (i) biomarker candidates – genes statistically associated with a disease phenotype; (ii) mechanistic candidates – genes with pathway or functional evidence linking them to disease biology; (iii) target-supported candidates – genes with disease-specific mechanistic or genetic evidence supporting therapeutic intervention; and (iv) clinically validated targets – genes with approved or late-stage therapeutics. Throughout this work, we use “disease-associated genes” as the general term for computationally prioritized genes across all tiers of this hierarchy.

The conventional workflow for discovery of disease-associated genes from transcriptomic data involves differential expression analysis between conditions, followed by pathway enrichment, literature review, and manual prioritization of candidates^4^. This approach suffers from well-documented limitations: differential expression is limited in its ability to identify non-linear patterns and gene interactions, it is sensitive to batch effects, and the synthesis of findings into mechanistic hypotheses requires domain expertise, which creates bottlenecks in the discovery pipeline^5,6^. The result is a widening gap between data generation and therapeutic insight, a critical barrier to realizing the clinical potential of transcriptomics.

Recent advances in foundation models for single-cell biology have scaled rapidly, from early efforts in cell-type annotation and transfer learning^7–9^ to large-scale models such as scGPT, Cell2Sentence (C2S), and STATE trained on tens to hundreds of millions of cells^10–16^. Yet despite this expansion, no consensus state-of-the-art model has emerged across cell-level tasks, and recent benchmarks have shown that simpler methods frequently match or exceed foundation models on standard evaluations such as cell-type annotation^17^. More fundamentally, the field’s focus on modeling individual cells overlooks a key asymmetry in clinical transcriptomics: while single-cell atlases provide cellular resolution, the bulk of clinically annotated patient data - with disease labels, treatment outcomes, and severity scores - exists at the patient level. Bridging from cell-level representations to patient-level predictions that identify disease states and disease-associated genes remains an active and only partially addressed challenge.

Emerging methods are beginning to enable patient-level analysis from single-cell data, representing patients as unordered collections of cells and learning aggregate representations for phenotype prediction through diverse strategies including mixture models, prototype networks, attention mechanisms, and tensor decomposition^18–22^. Of particular relevance is PaSCient, a single-cell patient foundation model that demonstrates how softmax-attention aggregation of cell embeddings, trained on over 2,790 patients, enables disease classification and provides interpretability through integrated gradients (IG)^23^. A parallel line of work has extended patient-level scRNA-seq analysis toward therapeutic prioritization, including approaches for personalized drug response prediction, patient-specific treatment selection, and network-based drug prioritization^24–26^, alongside early efforts to leverage large language models (LLMs) for automated gene prioritization from curated knowledge bases^27^. However, existing approaches typically address individual components of the translational workflow - classification, drug repurposing, or evidence curation - rather than providing an integrated pipeline that unifies patient representation learning, disease-associated gene discovery, evidence classification, and hypothesis generation within a single framework. A further limitation is that recently introduced patient-level methods that aggregate single-cell measurements into a unified patient representation are designed exclusively for scRNA-seq input and cannot be applied to bulk transcriptomic cohorts, leaving the majority of clinical and translational cohorts - which still rely on bulk RNA-seq - unable to benefit from the cellular resolution that single-cell training provides; true translational utility therefore requires a platform that can operate seamlessly across both modalities. Finally, current tools are largely optimized for classification tasks (e.g., disease labeling), whereas meaningful performance should instead be evaluated based on the accurate identification of disease-associated genes.

Here we introduce ConvergeCELL, an integrated platform that addresses this translational gap by providing an end-to-end pipeline from transcriptomic data to validated therapeutic hypotheses. Rather than optimizing for classification performance alone, ConvergeCELL is designed to automate the complete disease associated gene discovery workflow across both single-cell and bulk transcriptomic modalities. ConvergeCELL learns patient representations directly from single-cell data, or uses a distillation approach to project bulk RNA-seq data into single-cell resolution representations, enabling unified analysis; it then identifies candidate genes through IG and synthesizes predictions with biomedical knowledge to generate actionable hypotheses. Finally, our hypothesis generation workflow connects an LLM to biomedical knowledge bases to contextualize each candidate gene, classifying the strength of existing evidence and generating structured mechanistic hypotheses that can inform downstream experimental prioritization. ConvergeCELL bridges the gap between computational gene prioritization and the structured reasoning that informs experimental design and therapeutic development. We validated ConvergeCELL on studies held out from training, demonstrating that the platform recovers known disease-associated genes with higher or comparable recall to existing approaches.

### Platform architecture and design principles

ConvergeCELL is composed of four main components designed to create a robust platform for translational scientists. The first component is the dataset used for model training. Here, we curated a corpus of 4,479 patient-level single-cell transcriptomic samples derived from scBaseCount^28^, spanning over 20 million cells across more than 350 disease annotations (Fig. 1a). These annotations were further harmonized into nine disease families plus a healthy control category (Supplementary Fig. 1a; Methods).

**Figure 1.**
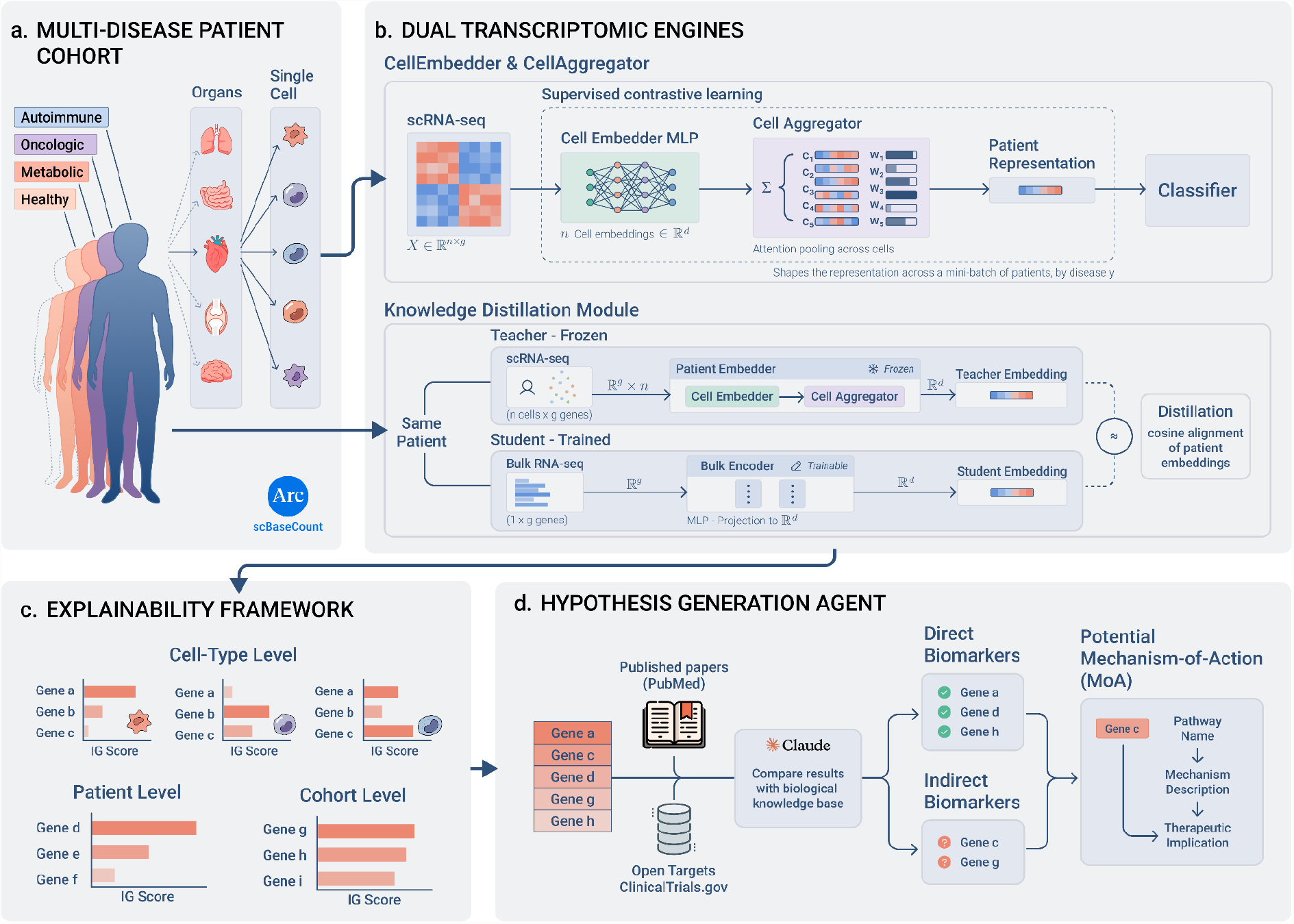
ConvergeCELL platform overview: from patient cohorts to therapeutic hypotheses. ConvergeCELL integrates four components, applied to a large-scale multi-disease patient cohort, to automate the translational workflow from transcriptomic data to actionable therapeutic hypotheses. **(a)** The input - a multi-disease patient cohort of 4,479 samples across more than 350 diseases curated from scBaseCount, organized into nine biologically meaningful disease families (e.g., immune-inflammatory, oncological, metabolic-vascular) plus a healthy control category, with patient samples spanning multiple organs and tissues. **(b)** Dual transcriptomic engines that support both single-cell and bulk modalities. The single-cell engine (top) follows a “bag of cells” architecture: a CellEmbedder multi-layer perceptron (MLP) maps individual cell expression profiles X ∈ ℝ^(n×g) to dense d-dimensional embeddings, a CellAggregator computes attention-weighted combinations across the n cells of each patient to produce a patient-level representation, and a Classifier predicts disease family membership. Training employs supervised contrastive learning, shaping patient embeddings z across mini-batches by disease label y. The Knowledge Distillation Module (bottom) extends the platform to bulk RNA-seq: paired bulk and single-cell profiles from the same patient generate a “Teacher” embedding (frozen single-cell encoder) and a “Student” embedding (trainable bulk encoder MLP), with cosine alignment of patient embeddings as the distillation objective. Once trained, the bulk encoder maps bulk profiles directly into the same patient embedding space, enabling the full downstream pipeline to operate on either modality. **(c)** The explainability framework applies IG with multi-level aggregation across three scales: patient-level gene attributions, cell-type-level rankings (e.g., for individual cell populations within a patient), and cohort-level IG scores that identify genes consistently driving disease predictions across patients. **(d)** The hypothesis generation agent connects an LLM to biomedical knowledge bases - including PubMed, Open Targets (OT) and ClinicalTrials.gov - via the MCP to classify each candidate disease-associated gene as direct (direct published evidence linking it to disease) or indirect (model-predicted but lacking direct validation), and generates structured mechanism-of-action hypotheses comprising pathway context, mechanistic description, and therapeutic implications for prioritized targets.

The second component is a patient representation model that learns unified embeddings from transcriptomic data, and is trained to classify disease states across multiple indications. The patient representation model follows the “bag of cells” paradigm, adopting the CellEmbedder-CellAggregator-Classifier architecture introduced by PaSCient while modifying the training objective, network depth, and training corpus to better capture cross-disease biological structure. The CellEmbedder maps expression profiles from individual cells to dense vectors, the CellAggregator computes learnable-weighted combinations of cell embeddings, and the PatientClassifier predicts disease family membership (Fig. 1b, top). Training employs supervised contrastive learning at the disease family level - grouping diseases into categories such as immune-inflammatory, oncologic, metabolic, and neurodegenerative - which encourages the model to learn shared pathophysiological signatures while maintaining disease-specific features. This hierarchical organization may potentially facilitate transfer to novel conditions not seen during training (see Methods for disease family assignment details).

While single-cell resolution provides informative input for the patient representation model, the majority of clinical transcriptomic data - including retrospective cohorts and routine clinical samples - exists as bulk RNA sequencing. To maximize clinical applicability, ConvergeCELL incorporates a knowledge distillation module that extends learned representations to bulk RNA sequencing data (Fig. 1b, bottom). The distillation module trains a separate BulkEmbedder network (“Student”) to map bulk expression profiles into the same embedding space as single-cell-derived patient representations (“Teacher”). The single-cell model is trained first and then frozen, and the BulkEmbedder is subsequently trained to align its output embeddings with the frozen single-cell patient embeddings by minimizing cosine distance, using paired single-cell and pseudobulk profiles from the same patients in the training corpus as input (Methods). This design enables the complete downstream pipeline (explainability and hypothesis generation) to operate seamlessly on either modality, allowing ConvergeCELL to leverage both the biological resolution of emerging single-cell datasets and the scale of existing bulk repositories.

The third component is the explainability framework, where IG are applied with multi-level aggregation to identify genes driving disease predictions at the level of patients, cell types, and cohorts (Fig. 1c). Briefly, IG quantifies each gene’s contribution to a model’s prediction by accumulating the model’s gradient with respect to that gene along a path from a reference baseline (representing the “absence” of signal) to the actual input, providing a principled measure of which genes drive disease-specific predictions. The fourth component of ConvergeCELL is a hypothesis generation agent that contextualizes each disease-associated gene by connecting an LLM to biomedical knowledge bases via the Model Context Protocol (MCP) (Fig. 1d). Each gene is classified as supported by direct evidence - published experimental or clinical data linking it to the disease, including known drug targets, validated biomarkers, or curated disease associations - or indirect evidence, where the gene lacks disease-specific validation but is implicated through pathway membership, functional analogy, or association with related conditions. This distinction supports downstream interpretation, separating genes with established disease relevance from those that may represent novel candidates requiring further validation.

### Patient classification generalizes across diseases and modalities

Using the aforementioned training dataset (Fig. 1a), patient embeddings were learned using a disease-category-aware contrastive training strategy. This approach encouraged patients from the same disease family, rather than the same specific disease, to cluster in representation space, yielding a clear family-level structure (Supplementary Fig. 1b). On a study-aware held-out validation set, ConvergeCELL achieved a weighted F1 of 0.76, performing comparably to a principal component analysis + XGBoost (PCA + XGB) baseline on raw expression and three single-cell foundation models with mean-pool aggregation (scGPT, STATE, C2S); pairwise differences in weighted F1 were not statistically significant by 1,000-resample bootstrap (Supplementary Fig. 1c). The fact that a simple PCA+XGB baseline on raw pseudobulk expression was statistically indistinguishable from ConvergeCELL suggests that patient-level disease family classification is largely attainable with standard methods. Importantly, this classification task is not itself the primary objective or intended downstream use, but rather serves as a structured proxy during training to shape informative embeddings. As such, classification performance alone is insufficient to distinguish between representation strategies. Instead, the value of the learned representations lies in the biological structure they encode. This structure becomes evident when interrogated using explainability methods, which constitute the more relevant and actionable use case, as demonstrated in the disease-specific validations below.

To evaluate generalization, we applied ConvergeCELL to three clinically distinct disease cohorts, each entirely excluded from model training: systemic lupus erythematosus (SLE), using a harmonized peripheral blood mononuclear cell (PBMC) atlas^29^; multiple myeloma (MM), using a harmonized bone marrow (BM) atlas^30^; and sepsis severity, using an independent bulk RNA-seq cohort of whole-blood samples^31^. These three contexts span oncological and immune-inflammatory biology, two tissue compartments (blood and BM), and both single-cell and bulk modalities. For each disease, we compared ConvergeCELL against three baselines: donor-aware pseudobulk differential expression analysis (DE), a standard ML baseline (PCA+XGB trained on the same multi-disease scBaseCount corpus), and for the single-cell diseases, PaSCient (30.8M parameters)^23^. All methods were compared on both classification performance and disease-associated gene recovery against Open Targets^32^. Because SLE is absent from PaSCient’s nine-class label space, multiple sclerosis (MS) - the closest immune-inflammatory condition available - was used as a proxy for both classification and IG attribution (for MM, PaSCient has a native disease class, enabling direct comparison). The Classification task was designed to reflect clinical relevance by using zero-shot model prediction to classify disease states (see disease state breakdown in Supplementary Table 1). In the sepsis setting, the pretrained BulkEmbedder was used to map each bulk RNA-seq sample into the patient embedding space without any further training or fine-tuning. An XGB classifier trained on these embeddings over the multi-disease scBaseCount training corpus (10-class disease-family labels) was then applied directly to the sepsis cohort to produce disease-family predictions. The standard ML baseline used an analogous PCA(512)+XGB classifier trained identically on the same scBaseCount corpus and applied zero-shot in the same way; the two methods therefore differ only in the upstream feature space (distillation embeddings versus PCA-reduced raw expression), isolating the contribution of the representations (Methods).

Across the single-cell settings, ConvergeCELL achieved the highest AUROC, outperforming PaSCient and the standard ML baseline in both SLE (AUROC 0.87 vs 0.67 and 0.76; Bonferroni-corrected DeLong p < 0.001 and p = 0.004, respectively; Fig. 2a) and MM (AUROC 0.72 vs 0.50 and 0.41; Bonferroni-corrected DeLong p = 0.005 and p < 0.001, respectively; Fig. 2b). In SLE, the standard ML baseline performed competitively (AUROC 0.76), reflecting that case-control classification can be achieved by expression-based methods alone. In MM, the standard ML baseline failed to discriminate between MM and its precursor states (AUROC 0.41), consistent with the greater difficulty of within-disease staging compared with case-control classification - while ConvergeCELL maintained meaningful discrimination at AUROC 0.72. In sepsis, ConvergeCELL achieved AUROC 0.56 versus 0.38 for the standard ML baseline (DeLong p < 0.001; Fig. 2c). Since all methods were applied entirely zero-shot across the three cohorts, classification performance was evaluated as disease versus healthy by treating any non-healthy predicted class as a positive prediction (full classification metrics are provided in Supplementary Fig. 2; precision-recall curves and bootstrap confidence intervals in Supplementary Fig. 3). Confusion matrices revealed clinically interpretable prediction patterns across all three diseases: in SLE, healthy controls were correctly classified in 88% of cases while managed and flare patients were assigned to the immune-inflammatory class in 64% and 81%, respectively (Fig. 2d); in MM, a graded pattern emerged with monoclonal gammopathy of undetermined significance (MGUS) predominantly classified as healthy (88%), smoldering multiple myeloma (SMM) evenly split between oncological and healthy (50% each), and MM showing a roughly comparable oncological rate (46%) but the lowest healthy classification rate (32%) and the greatest diversity of non-healthy assignments, including immune-inflammatory (16%) absent in earlier stages (Fig. 2e). In sepsis, healthy controls were correctly classified in 82% of cases, while most of Low Sequential Organ Failure Assessment (SOFA) patients (54%) and High SOFA patients (52%) were assigned to non-healthy classes - predominantly metabolic-vascular and immune-inflammatory, disease families that are aligned with sepsis pathophysiology^33^ (Fig. 2f). Low SOFA samples served as part of the negative class because they lacked acute sepsis, but they were not healthy individuals (all had presented to the emergency room with some clinical concern). ConvergeCELL’s classification of many such samples as non-healthy is therefore consistent with detection of potentially underlying non-sepsis pathology rather than a model error. The confusion matrices for the control baselines were generally less interpretable (Supplementary Fig. 4).

**Figure 2.**
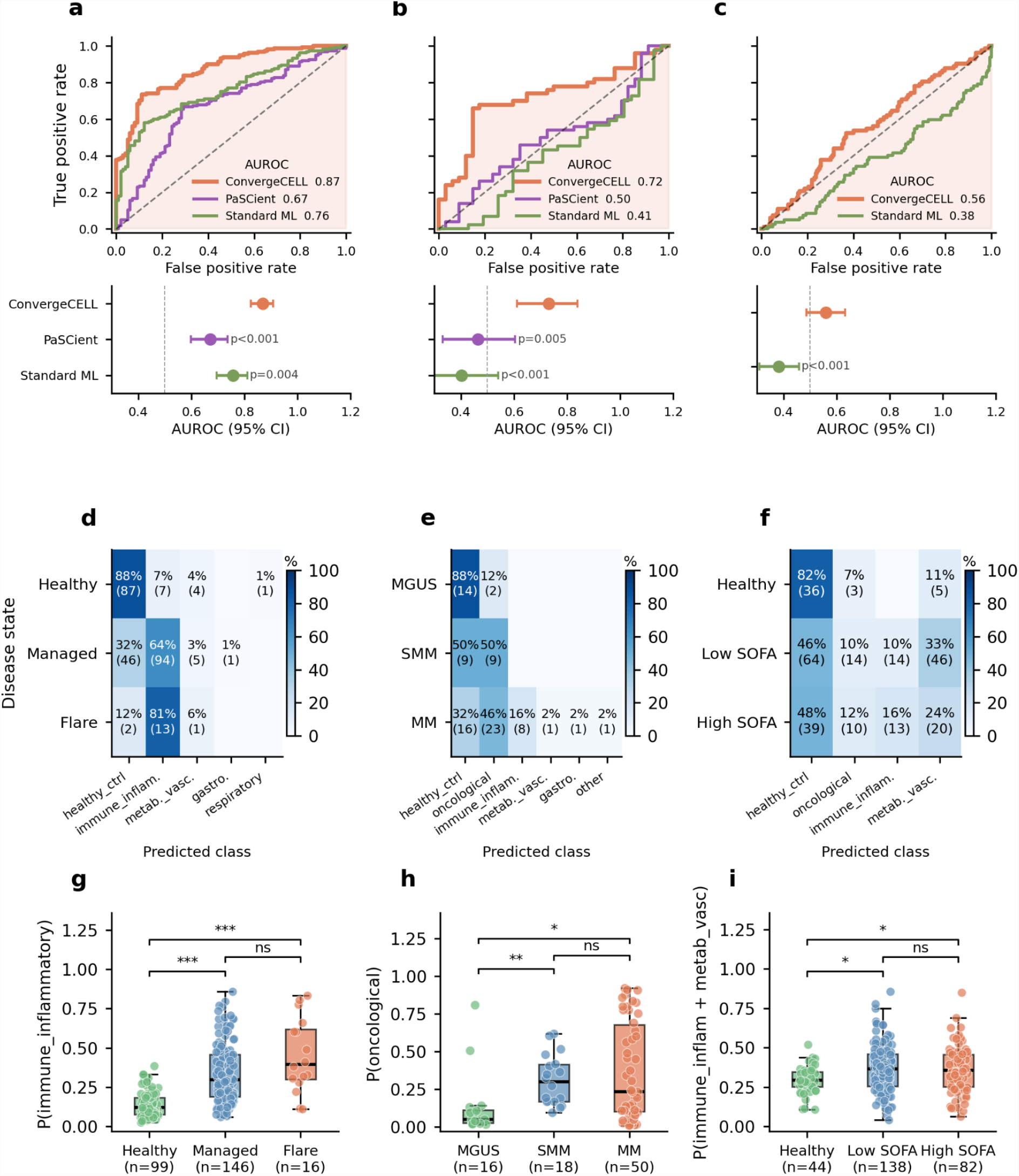
ConvergeCELL’s patient classification generalizes across diseases and modalities. onvergeCELL was evaluated on three held-out cohorts spanning oncological and immune-inflammatory biology, two tissue compartments, and both single-cell and bulk modalities. **(a-c)** Classification performance for **(a)** SLE (flare and managed) versus healthy control on a harmonized PBMC atlas of 261 donors across four cohorts, **(b)** MM versus non-MM (SMM and MGUS) on a harmonized BM atlas of 84 samples across six studies, and **(c)** sepsis disease-versus-healthy classification on an independent bulk RNA-seq cohort of 264 samples (positive class: High SOFA ≥ 5; negative class: Low SOFA < 2 combined with healthy controls). Upper: ROC curves with AUROC values for ConvergeCELL (orange), PaSCient (purple, single-cell diseases only), and the standard ML baseline (green); dashed lines indicate random classifier performance. Lower: AUROC point estimates with 95% bootstrap confidence intervals (1,000 resamples) and Bonferroni-corrected DeLong test p-values for pairwise comparisons between ConvergeCELL and each baseline. **(d-f)** Confusion matrices showing predicted class (columns) stratified by clinical stage or severity group (rows) for **(d)** SLE (healthy, managed, flare), **(e)** MM (MGUS, SMM, MM), and **(f)** sepsis (healthy, Low SOFA, High SOFA), with predicted classes shown across disease families that received assignments. Values indicate the percentage and absolute number of samples assigned to each predicted class. Cell values are rounded to integer percentages and may not sum to 100. **(g-i)** Distribution of ConvergeCELL’s predicted class probability across clinical stages for **(g)** SLE (predicted immune-inflammatory probability across healthy, managed, and flare), **(h)** MM (predicted oncological probability across MGUS, SMM, and MM), and **(i)** sepsis (predicted probability of immune-inflammatory and metabolic-vascular disease family assignment), shown across healthy, Low SOFA, and High SOFA groups. Post-hoc pairwise comparisons by Mann-Whitney U test with Bonferroni correction. *p < 0.05, **p < 0.01, ***p < 0.001, ns: not significant.

Beyond discrete classification, predicted class probabilities tracked clinical severity across diseases (Fig. 2g-i). In SLE, predicted immune-inflammatory probability increased from healthy controls through managed SLE to flare (Kruskal–Wallis p < 0.001; pairwise Bonferroni-corrected Mann–Whitney: HC vs managed p < 0.001, HC vs flare p < 0.001), though managed versus flare did not reach significance after correction (p = 0.16; Fig. 2g). In MM, predicted oncological probability increased from MGUS through SMM to MM (Kruskal–Wallis p = 0.003), with significant separation between MGUS and later stages (MGUS vs SMM p = 0.001, MGUS vs MM p = 0.011) but not between SMM and MM (p = 1.0; Fig. 2h). In sepsis, the predicted probability of belonging to a sepsis-aligned disease family (immune-inflammatory or metabolic-vascular) was significantly higher in Low and High SOFA samples than in healthy controls (Kruskal–Wallis p = 0.010; Healthy vs Low SOFA p = 0.013, Healthy vs High SOFA p = 0.017; Low vs High SOFA p = 1.0; Fig. 2i).

### Disease-associated gene recovery reveals the value of learned representations

ConvergeCELL’s primary objective is identifying disease-associated genes. To evaluate this capability, we used Open Targets’ overall association score as reference, with significant associations identified at z-score ≥ 1.5 (383 SLE-, 568 MM-, and 236 sepsis-associated genes). For each of the three aforementioned cohorts, disease-associated genes were recovered in the same zero-shot setup described above. ConvergeCELL and PaSCient rank genes by IG attribution. The standard ML baseline ranks genes by TreeSHAP (Shapley values for tree models) values back-projected through PCA loadings, and DE ranks genes by adjusted p-value from pseudobulk DESeq2, in which single-cell counts are aggregated per donor within the disease-relevant cell type prior to testing (Methods; see comparison to cell-aware Wilcoxon DE in Supplementary Fig. 5). Explainability was computed over disease-relevant cell types for the single-cell cohorts: CD4+ T (T4) cells for SLE, given their central role in driving autoantibody production through aberrant B cell help^34^, and plasma cells (PC) for MM, as the defining neoplastic compartment of the disease^35^ (cell type distributions across diseases are shown in Supplementary Fig. 6). To ensure fair comparison across methods with different gene vocabularies, all single-cell analyses were restricted to the intersection of the ConvergeCELL (18,301 genes) and PaSCient (28,231 genes) vocabularies, yielding 16,040 shared genes; pseudobulk DE was further restricted to the subset of these genes passing its expression filter (see Methods).

We evaluated disease-associated gene recovery using two complementary metrics across top-K cutoffs of K = 25, 50, 100, and 200. Gene Recall@K measures the fraction of Open Targets-significant genes recovered within each method’s top-K ranked genes, providing a binary-thresholded view of how efficiently each method concentrates known disease genes. Mean OT score@K measures the average Open Targets overall association score across each method’s top-K genes, capturing whether top-ranked genes carry stronger disease associations on average using the full continuous score rather than a significance threshold.

Across the single-cell settings, ConvergeCELL led on Gene Recall@K at almost every K tested (Fig. 3a-b). In SLE, ConvergeCELL reached 0.034 at K=200, compared with 0.029 for pseudobulk DE, 0.021 for PaSCient, and 0.018 for the standard ML baseline (Fig. 3a). In MM, PaSCient and ConvergeCELL were the top two methods at K=200 (0.035 and 0.034, respectively), with the standard ML baseline close behind (0.030); however, ConvergeCELL was the only method significantly above the permutation null at every cutoff (Fig. 3b). In bulk sepsis, ConvergeCELL and the standard ML baseline performed comparably across most K values, with the standard ML baseline surpassing ConvergeCELL at K=200 (Fig. 3c).

**Figure 3.**
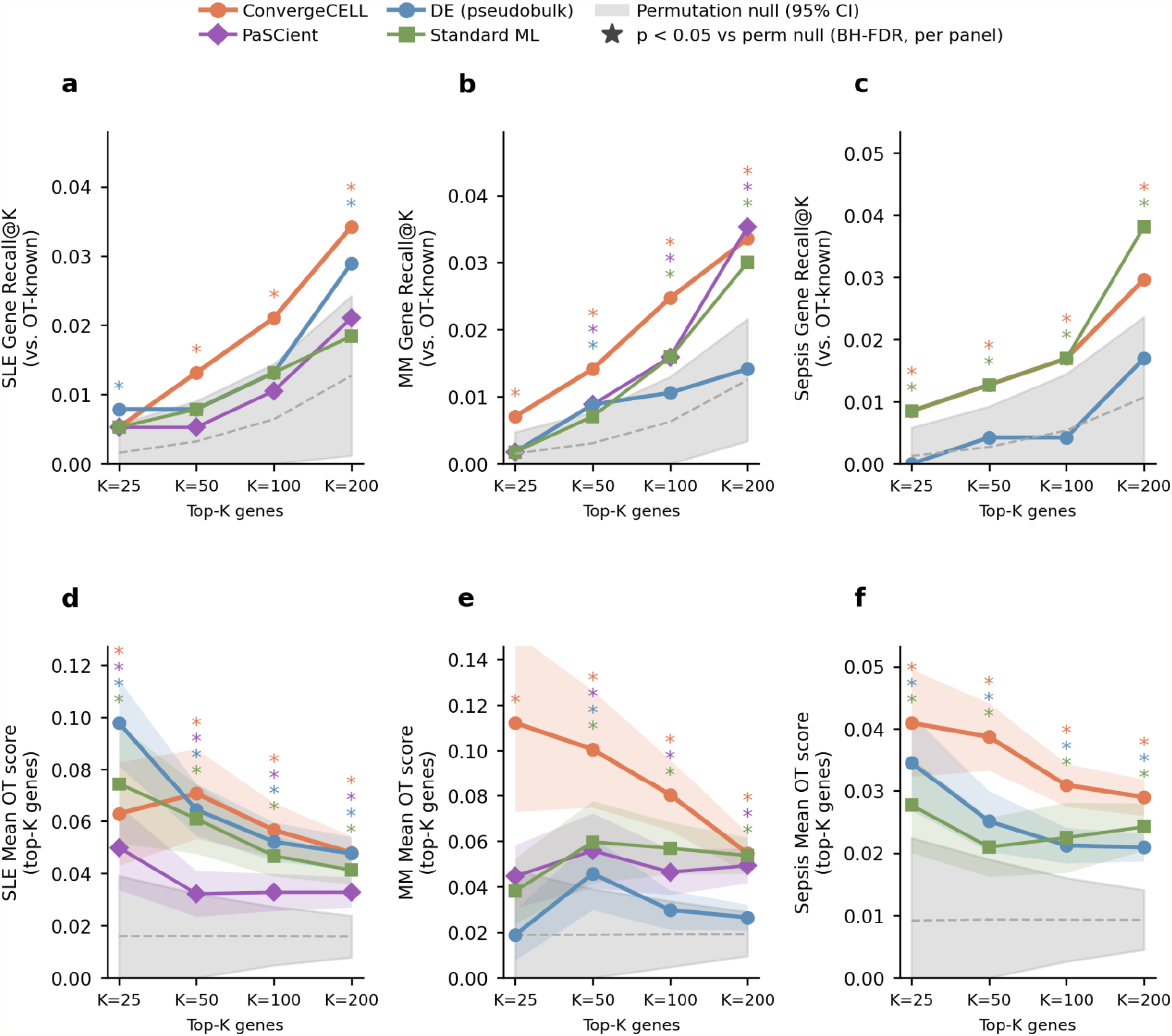
Disease-associated gene recovery reveals the value of learned representations. **(a-c)** Gene Recall@K, the fraction of Open Targets-significant disease-associated genes (z-score ≥ 1.5; 383 SLE-, 568 MM-, 236 sepsis-associated genes) appearing within the top-K ranked genes per method, evaluated at K = 25, 50, 100, 200, for **(a)** SLE, **(b)** MM, and **(c)** sepsis. **(d-f)** Mean OT score@K, the arithmetic mean of Open Targets overall association scores across the top-K ranked genes per method, for **(d)** SLE, **(e)** MM, **(f)** sepsis. ConvergeCELL (orange), PaSCient (purple, single-cell diseases only), pseudobulk DE (blue), and Standard ML (green) are shown. Colored shaded bands around each method’s curve in (d-f) represent the standard error of the mean across the top-K genes. Grey shading indicates the 95% confidence interval of a permutation null (1,000 permutations). Asterisks denote p < 0.05 versus the permutation null by z-test (Benjamini-Hochberg false discovery rate (FDR)-corrected within each panel). All methods evaluated on a shared gene universe of 16,040 genes.

Mean OT score@K provides a continuous-relevance complement to Gene Recall@K (Fig. 3d-f). In SLE at K=25, pseudobulk DE achieved the highest mean score (0.10), whereas ConvergeCELL took the lead from K=50 onwards (Fig. 3d). In MM, ConvergeCELL led across all K, with the largest margins at low K (∼2.5-fold above PaSCient and the standard ML baseline at K=25) and a narrowing gap at K=200, where the standard ML baseline approached parity (Fig. 3e). In sepsis, ConvergeCELL led at all K, with the standard ML baseline narrowing the gap at higher K but never matching ConvergeCELL’s top-K disease relevance (Fig. 3f). PaSCient’s mean OT score plateaued well below ConvergeCELL across both single-cell diseases. These results are broadly consistent across alternative Open Targets significance thresholds (Supplementary Fig. 7) and hold when evaluated against orthogonal Open Targets sub-scores that exclude potentially circular expression and literature evidence (Supplementary Fig. 8).

Beyond aggregate metrics, we asked whether each method surfaces the genes targeted by known drugs used to treat these diseases. Using Open Targets, we compiled approved drugs for each disease and identified the gene encoding each drug’s primary target. For drugs with multiple annotated targets, we manually curated a single primary target based on FDA labeling and clinical literature, excluding cases where no clear primary mechanism could be assigned (Fig. 4; Methods; Supplementary Data 1 lists all approved drugs and curated targets). Across all approved drugs evaluated in the shared gene space, ConvergeCELL achieved the lowest mean rank in SLE (1,858; Fig. 4a) and was comparable to the standard ML baseline in MM (2,017 vs. 1959; Fig. 4b), with both ranking lower than PaSCient (3,685) and DE (6,894). Given the limited number of approved drugs per disease (n = 4 SLE; n = 10 MM), we interpret these aggregate ranks descriptively rather than as a statistical comparison between methods. Looking at individual drug-target pairs, a qualitative pattern emerges: ConvergeCELL ranks the most disease-specific monoclonal-antibody targets markedly higher than any baseline. Belimumab/TNFSF13B (one of only two FDA-approved drugs developed specifically for SLE) is ranked 26 by ConvergeCELL versus 179 (DE), 1,745 (PaSCient), and 7,095 (standard ML). Belantamab mafodotin/TNFRSF17 in MM is ranked 3 by ConvergeCELL, and plerixafor/CXCR4 is ranked 28. Daratumumab/CD38, a backbone of modern MM therapy, also receives its best rank from ConvergeCELL (802 vs. 2,127-4,088). The standard ML baseline ranks higher on tubulin (vincristine/TUBB) and the proteasome (bortezomib/PSMB5) - generic cytotoxic and proteasome biology that does not require disease-specific representation learning to surface^36^. Sepsis was excluded from this analysis as it lacks strong disease-mechanism-targeted therapies, with most approved drugs directed at the underlying bacterial infection rather than the host response^37^.

**Figure 4.**
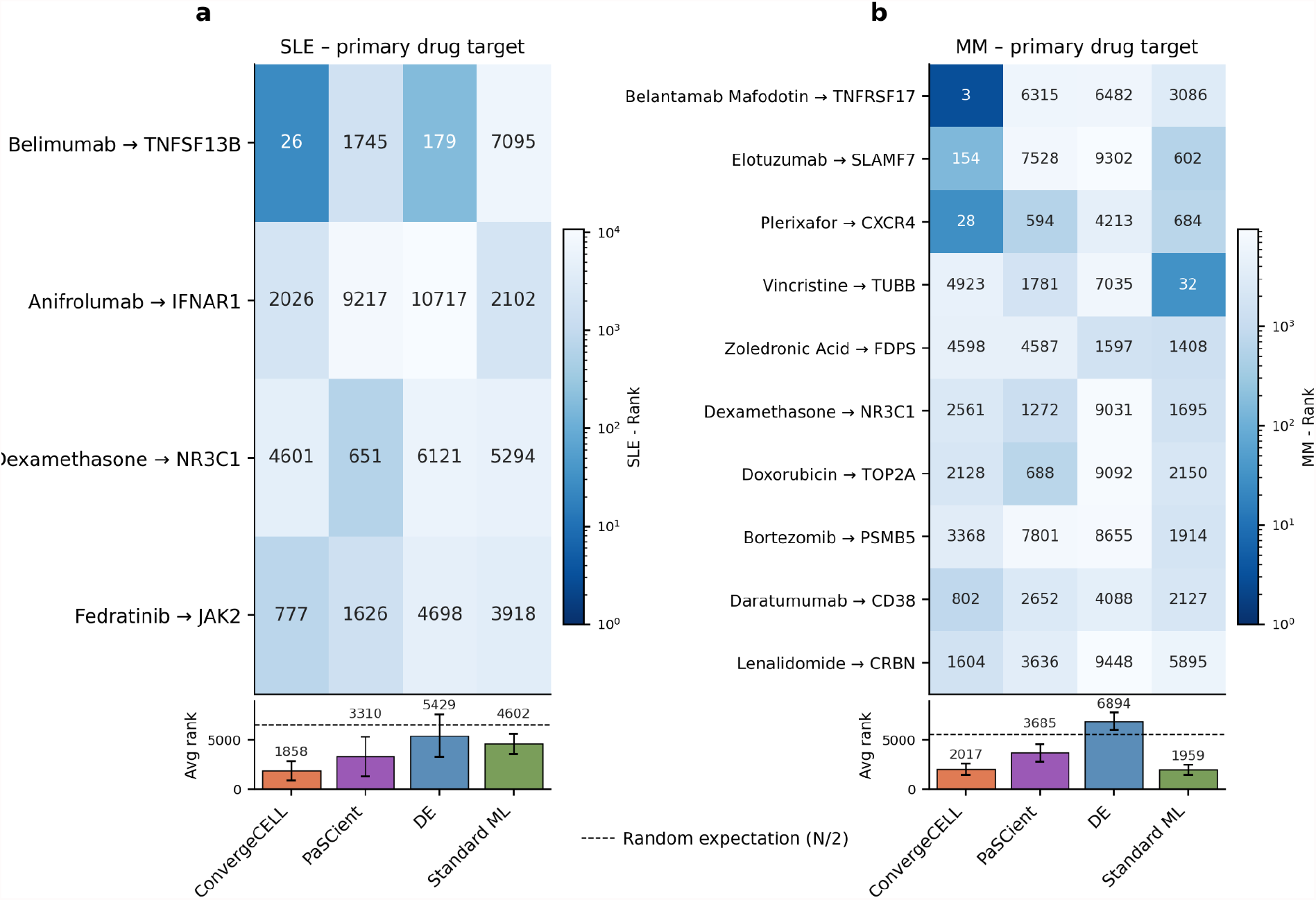
ConvergeCELL prioritizes drug targets across approved and investigational therapies. **(a-b)** ConvergeCELL prioritizes clinically validated drug targets in SLE and MM. Heatmaps show the rank assigned by each model to the curated primary mechanism-of-action target gene for each approved drug (one drug per row, one model per column; lower rank = better; log color scale). Drugs are restricted to those at maximum clinical stage ‘Approval’ in Open Targets, and for each drug a single primary target gene was curated based on FDA labeling and clinical literature (Methods). Bottom bar plots show the mean rank across all drugs in each panel for each model. (a) SLE, 4 approved drug-primary target pairs. (b) MM, 10 approved drug-primary target pairs. Error bars represent the standard error of the mean.

The biological coherence of ConvergeCELL’s top-ranked genes further supports the quality of its representations (Fig. 5). In SLE, the highest-attributed genes in T4 cells included the alarmins S100A8/A9 and the chemokine receptor CXCR4, which marks T-cell trafficking to the lupus kidney^38^, alongside CTSS, an actively pursued SLE drug target whose inhibition suppresses anti-dsDNA and nephritis in lupus models^39^ (Fig. 5a). Reactome enrichment recovered the canonical T-cell receptor (TCR) and costimulatory pathways dysregulated in lupus T cells (PD-1 signaling, CD28-VAV1, AP-1) (Fig. 5d), with AP-1/JUN recently linked to pathogenic CXCL13^+^ T-cell programs in SLE^40^. In MM, top-ranked PC genes were dominated by clonotypic immunoglobulin chains and the FDA-approved drug target TNFRSF17 (BCMA, targeted by belantamab mafodotin)^41^, alongside LAMP5, recently characterized as a malignant-plasma-cell-selective antigen and emerging antibody-drug conjugate (ADC/chimeric antigen receptor T-cell (CAR-T) target^42^ (Fig. 5b). Pathway enrichment surfaced ATF6α-driven chaperone induction at the top - the unfolded protein response axis whose activation is causal for proteasome-inhibitor sensitivity in MM^43^ (Fig. 5e). In sepsis, the top-ranked genes recapitulate two established arms of the host response: emergency granulopoiesis (CD177, VNN1, ANXA3) and platelet/megakaryocyte activation (PPBP)^44,45^. IFI27, also among the top genes, is an interferon-driven blood biomarker with established translational traction across independent sepsis cohorts^46^. Reactome enrichment recovered complement, FcγR, and scavenger-receptor programs central to sepsis pathophysiology (Fig. 5f), pathways that operationally couple these three arms through neutrophil effector function, antibody opsonisation, and IFN-primed clearance. Beyond direct-evidence genes, ConvergeCELL also surfaces indirect-evidence candidates that may warrant further investigation. Top-10 gene rankings and pathway enrichments for the baseline methods are provided in Supplementary Figs. 9-11 (full top-200 gene rankings for all methods across all three diseases in Supplementary Data 2; curated Reactome pathway reference sets in Supplementary Data 3; hypothesis-generation outputs for SLE, MM, and sepsis in Supplementary Data 4-6).

**Figure 5.**
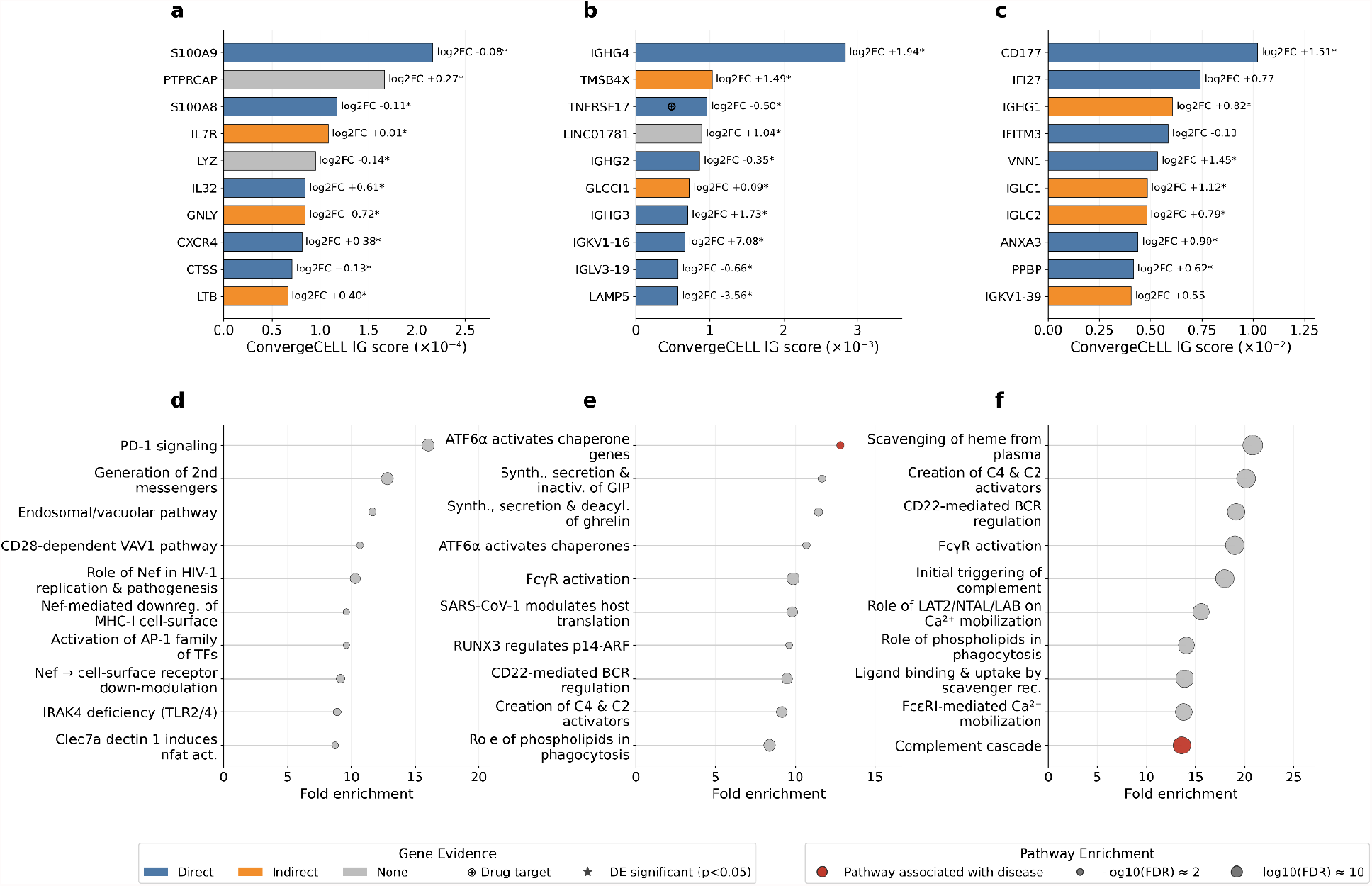
ConvergeCELL top-attributed genes and pathway enrichments across diseases. **(a-c)** Top 10 ConvergeCELL-attributed genes ranked by IG score for **(a)** SLE in T4 cells, **(b)** MM in PC, and **(c)** sepsis in whole blood. Bar color indicates evidence category as classified by the hypothesis generation workflow: direct evidence (blue), indirect evidence (orange), and no evidence (grey). The ⊕ symbol marks genes that are approved drug targets for the disease shown. Asterisks next to log2 fold-change values denote genes also reaching significance by DE (p < 0.05); log2 fold-change values are shown to the right of each bar. **(d-f)** Top enriched Reactome pathways from over-representation analysis on each disease’s top-K ConvergeCELL gene list (K = 500) for **(d)** SLE, **(e)** MM, and **(f)** sepsis. Dot size encodes -log10(FDR); pathways also represented in the curated Reactome reference set for each disease are shown in red.

Together, these results show that ConvergeCELL’s learned representations encode disease-associated biology that surfaces consistently across diseases and modalities, and that the platform prioritizes clinically validated, disease-specific drug targets - including monoclonal-antibody mechanisms that define modern therapy in SLE and MM - more sharply than expression-based or alternative patient-level baselines.

## Discussion

ConvergeCELL addresses a critical gap in translational transcriptomics: the absence of integrated tools that bridge raw molecular data and actionable therapeutic hypotheses. By unifying patient representation learning, gene-level interpretability, and LLM-driven hypothesis generation within a single framework, extensible to both single-cell and bulk transcriptomic data, ConvergeCELL automates a workflow that currently requires weeks of manual expert curation per disease context. Its validation across SLE, MM, and sepsis yields three findings that we believe matter beyond the platform itself.

First, our results highlight an important dissociation between aggregate metrics - whether classification performance, mean gene-recovery scores, or average drug ranks - and the prioritization of clinically meaningful, disease-specific targets. The clearest example emerges in MM, where the standard ML baseline does not meaningfully distinguish MM from precursor states (AUROC 0.41), yet achieves aggregate Open Targets scores and an average approved-drug rank comparable to ConvergeCELL’s. This apparent parity is, however, driven by high ranks on generic cytotoxic targets such as tubulin^36^ (vincristine/TUBB, rank 32), while the disease-defining targeted therapies that anchor modern treatment - including belantamab/TNFRSF17 (rank 3,086) - remain deeply buried. ConvergeCELL, by contrast, ranks TNFRSF17 third out of 11,198 genes. This pattern indicates that aggregate scores can obscure substantial differences in the biological specificity of what each method surfaces. The same dissociation is visible in SLE for belimumab/TNFSF13B (7,095 versus ConvergeCELL’s 26), and again in MM for plerixafor/CXCR4 (684 versus 28) and daratumumab/CD38 (2,127 versus 802). ConvergeCELL’s consistent advantage across these clinically defining targets points to a concrete implication: patient-level transcriptomic models intended for translational use should be evaluated, in part, on the recovery of specific clinically actionable targets, not only on aggregate metrics or predictive accuracy.

Second, organizing diseases into biologically meaningful families before training reflects a design choice that may have implications beyond this platform. By grouping conditions with shared pathophysiology rather than treating each disease as an isolated class, models are encouraged to learn axes that capture immune-inflammatory, oncological, metabolic-vascular biology and more in their general forms - axes that then transfer to held-out diseases never seen during training. The graded prediction probabilities tracking SLE flare severity and MM stage progression, despite training labels being family-level rather than stage-level, are consistent with this transfer. Supporting this interpretation, the standard ML baseline - which was trained on the same family-organized corpus - also recovered disease-relevant biology in aggregate, suggesting that the family-level scaffold itself contributes to gene recovery independently of the representation architecture. This argues against the prevailing pattern in disease prediction models of treating every disease as its own class, which fragments shared biological signals across narrow boundaries.

Third, ConvergeCELL extends to bulk RNA-seq through knowledge distillation, broadening the platform’s translational reach to the modality where most clinical transcriptomic data actually lives. Patient-level foundation models in single-cell biology have demonstrated impressive performance on cellular atlases, but bulk RNA-seq remains the dominant modality across clinical biobanks and retrospective cohorts. The sepsis validation demonstrates that ConvergeCELL recovers known disease biology - the neutrophil, interferon, and humoral arms of the bacterial sepsis host response - directly from bulk profiles, converting methodological progress into deployable utility on existing clinical datasets.

The platform’s contribution should be understood within several boundaries. PaSCient’s fixed nine-class label space limits the comparison, with SLE absent and MS used as the nearest immune-inflammatory proxy. We deliberately evaluated all methods zero-shot to reveal what the pretrained representations have learned, though fine-tuning will often be necessary in applied settings. The training corpus remains biased toward immune-inflammatory and oncological conditions (>37% of all training samples), and the hypothesis generation workflow inherits the limitations of current LLMs, including potential for hallucination, with systematic reproducibility evaluation an important direction for future work (an illustrative output for MMP25 in sepsis is shown in Supplementary Fig. 12).

Future directions follow naturally. Prospective validation partnerships will test whether ConvergeCELL’s prioritized targets translate to clinical settings beyond retrospective cohorts, and integration with orthogonal modalities - genetic variation, proteomics - could strengthen the evidence base.

In summary, ConvergeCELL provides an end-to-end platform for translating transcriptomic data into therapeutic hypotheses. The consistent advantage in disease-associated gene recovery, the systematic prioritization of clinically used drug targets, and the extension to bulk RNA-seq together demonstrate that the value of learned representations extends beyond prediction performance - a principle that may guide the development and evaluation of future patient-level transcriptomic models.

## Methods

### Data curation and processing

Raw scRNA-seq count matrices were obtained from scBaseCount^28^, a curated repository aggregating publicly available datasets. Study-level metadata was enriched via the NCBI Sequence Read Archive (SRA) Entrez API, extracting study titles, abstracts, organism information, and library preparation protocols. For datasets distributed in pre-normalized form (e.g., CZ CELLxGENE), raw counts were recovered from the adata.raw layer prior to processing. Studies were selected based on availability of patient-level metadata including disease diagnosis and tissue of origin.

Quality control (QC) was performed using scanpy. Cells with fewer than 200 detected genes were removed, as were cells expressing more than 5,000 genes (putative doublets). Cells with 10% or greater mitochondrial read fraction were excluded. Genes detected in fewer than three cells were filtered. To reduce sparsity-driven noise, genes were ranked by their detection rate across the training corpus and the top 50% least sparse genes were retained (95% sparsity), yielding a fixed vocabulary of 18,301 genes. Following QC, library sizes were normalized to a target sum of 10,000 counts per cell, and the data were log1p-transformed. Gene symbols were standardized and sorted alphabetically, with duplicate gene names resolved by appending fallback identifiers.

### Disease family organization

Diseases were assigned to nine biologically meaningful disease families through a combination of manual expert curation and LLM-assisted annotation: oncological, immune-inflammatory, neurological, metabolic-vascular, gastrointestinal, respiratory, epithelial barrier, sensory specialized, and an additional ‘other’ category for diseases that did not fit the primary families. Healthy control samples constituted a separate tenth class for classification.

### Patient representation model

#### Architecture

The model adopts the CellEmbedder-CellAggregator-Classifier architecture introduced by PaSCient^23^, with modifications to network depth, training objective, and training corpus. The model comprises three modules. The full single-cell model comprises 80M (79,963,661) parameters; the BulkEmbedder distillation module adds an additional 10M (9,897,986) parameters.

#### CellEmbedder

A MLP that maps each cell’s log-normalized expression profile *x*_*i*_ ∈ ℝ^*G*^(*G* = 18, 301 *genes*) to a dense embedding *h*_*i*_ ∈ ℝ^*d*^(*d* = 512) through two hidden layers, where hidden dimensions are 4,096 and 1,024, with parametric rectified linear unit (PReLU) activation preceded by batch normalization and followed by dropout (*p = 0*.*1*) at each layer. For each patient, *C =* 500 cells are sampled and independently embedded as demonstrated by the PaSCient model^23^.

#### CellAggregator

Per-cell embeddings are combined into a single patient-level representation *z* ∈ ℝ^*d*^ using a DeepSets-style attention mechanism^47^. A two-layer attention network *g*_ϕ_ : ℝ^*d*^ → ℝ computes scalar scores over cells, defined as *g*_ϕ_ (*h*).

Scores are softmax-normalized to produce attention weights:

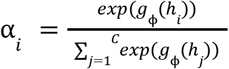

The patient embedding is the attention-weighted sum:

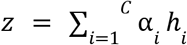

#### Classifier

A dropout layer (*p* = 0.1) followed by a single linear layer maps the patient embedding to class logits

#### Training objective

Pre-training employed the supervised contrastive loss at the disease family level^48^. For a batch of *N* patient embeddings 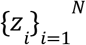 with disease family labels 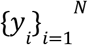. Let *P*(*i*) = {*j* : *y*_*j*_ = *y*_*i*_, *j* ≠ *i*} denote the set of positive samples sharing the same disease family label as anchor *i*, and *A*(*i*) = {1, …, *N*} \ {*i*} the set of all samples excluding the anchor. We define the SupCon loss as defined by Khosla et al.^48^ with temperature *τ* = 0.1. The loss is normalized by the base temperature *τ_base* = 0.07 following Khosla et al. All embeddings are L2-normalized prior to similarity computation. The loss encourages patients belonging to the same disease family to cluster in embedding space while separating patients from different families.

#### Training procedure

Model architecture (number of hidden dimensions and activations) was optimized via Bayesian search using Weights & Biases sweeps. The model was trained with supervised contrastive learning for 16 epochs using AdamW (*lr* = *1 ×*10 ^−5^, weight decay = 0.05) with a cosine learning rate schedule and 10% linear warmup. Batch size was 16; evaluation was performed every 100 steps. Hyperparameters were selected via Bayesian grid search over learning rate [10^−5^ - 10^−4^], temperature [0.01-0.5], and dropout [0.05-0.3], optimizing silhouette score and *k*-nearest-neighbour accuracy (*k =* 5) on the validation set. The classifier head was fine-tuned on the same training set using cross-entropy loss with entity-level averaging, which averages the loss across multiple stochastic views of each patient to prevent overrepresentation of patients with more augmented samples. The classifier was trained for 20 epochs using AdamW (*lr* = 10^−4^, weight decay = 0.05) with a cosine learning rate schedule, 10% linear warmup, and batch size of 32. Early stopping with patience of 10 epochs was applied based on validation loss. The same hyperparameter approach was taken.

Training was performed on the L40S GPU with 32 CPUs using PyTorch version 2.5.1 on CUDA 12.1 and took ∼48h for representation learning using supervised contrastive learning. Classification took ∼12h.

#### Evaluation protocol

Study-aware data splitting: To simulate real-world generalization, train/validation splits were constructed such that all samples from a given clinical study were assigned to the same split, preventing information leakage from shared batch effects or patient populations within a study.

At inference, five stochastic cell samples were drawn per patient. Predictions were aggregated across views by averaging softmax probabilities per patient before computing metrics. Performance was evaluated using accuracy, weighted precision, weighted recall, and weighted F1 score, each weighted by class support to account for class imbalance.

### Bulk RNA-seq knowledge distillation

To extend ConvergeCELL to bulk RNA-seq data, we trained a distillation module that maps bulk expression profiles into the same embedding space as single-cell-derived patient representations, enabling the full downstream pipeline to operate on either modality.

#### Pseudobulk profile generation

For each patient in the single-cell training corpus, a pseudobulk expression profile was generated by averaging raw counts across all cells per gene. The resulting profiles were library-size normalized to a target sum of 10,000 and log-transformed, mirroring the normalization applied to single-cell data during curation to ensure comparable expression scales for the distillation objective.

The frozen single-cell ConvergeCELL model was applied to each patient’s single-cell data to extract patient-level embeddings. When multiple stochastic views existed per patient, embeddings were averaged across views to produce a single teacher embedding.

#### BulkEmbedder architecture

An MLP maps each pseudobulk profile 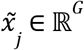 to a student embedding 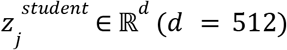 through two hidden layers of dimensions 512 and 512, with PReLU activations, batch normalization, and dropout (*p =* 0.1).

#### Training objective

The BulkEmbedder was trained to align student embeddings with frozen teacher embeddings by minimizing a cosine distillation loss: 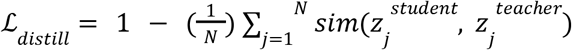, where 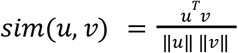 is cosine similarity, and both embeddings are L2-normalized prior to computation. Only the BulkEmbedder parameters are updated; all teacher model parameters remain frozen.

#### Training procedure

Model architecture (number of hidden dimensions and activations) was optimized via Bayesian search using Weights & Biases sweeps. The model was trained for up to 50 epochs using AdamW (*lr* = 9.8 × 10^−4^, weight decay = 0.12 and dropout = 0.2) with a cosine learning rate schedule, 10% linear warmup, and batch size of 64. Early stopping with patience of 10 epochs was applied based on validation cosine similarity. Hyperparameters were selected via Bayesian search using Weights & Biases sweeps, optimizing cosine similarity on the validation set over hidden dimensions {[512, 512], [1024, 512],[1024, 1024], [2048, 1024], [4096, 1024]}, dropout [0.05, 0.4], learning rate [1 × 10^−5^, 1 × 10^−3^], weight decay [1 × 10^−3^, 0.5], and batch size {32, 64, 128, 256}.

The distillation module learns only the embedding projection and does not include a classification head. Applying ConvergeCELL to a new bulk RNA-seq disease context requires fitting a lightweight downstream classifier on the resulting embeddings, as demonstrated in the sepsis severity evaluation.

### Filtration of single-cell benchmark datasets

#### Systemic lupus erythematosus

We obtained scRNA-seq data of PBMCs from an SLE cohort available through CZ CELLxGENE (collection 436154da-bcf1-4130-9c8b-120ff9a888f2) based on Perez et al.^29^. Raw count matrices were extracted from the adata.raw object. Gene identifiers, originally stored as Ensembl IDs, were converted to gene symbols using the feature_name annotation provided in the dataset. Cells from treated patients (disease_state == ‘treated’) were excluded to restrict the analysis to treatment-naive samples; both managed and flare SLE disease states were retained alongside healthy controls. No tissue filter was applied as all samples were derived from peripheral blood. Per-donor QC was performed independently on each of the 261 donors. Cells expressing fewer than 200 or more than 10,000 genes, and cells with mitochondrial gene content exceeding 20%, were removed. Genes detected in fewer than three cells per donor were excluded. Remaining counts were log-normalized per cell. Gene expression profiles were aligned to a fixed vocabulary of 18,301 genes used by ConvergeCELL; genes absent from a given donor were zero-filled, and genes outside the vocabulary were dropped. For disease classification and explainability, SLE donors (flare and managed) were assigned a positive label, while healthy donors were assigned a control label.

#### Multiple Myeloma

The PanImmune data was downloaded from the single-cell atlas of MM^49^. Raw count matrices were extracted from the counts layer of the original AnnData object. Donor-level clinical metadata (cohort assignment, clinical variables) were merged with per-cell annotations. We restricted the analysis to BM aspirate samples from donors flagged as analysed by the original study authors, retaining only those belonging to the MM, SMM, or MGUS disease cohorts. Predicted doublets (identified via scrublet, scr_doublet == True) and cells annotated as Doublet, Platelet, or Nonhaematopoietic lineages were excluded. Donors with fewer than 500 cells after filtering were removed. Per-donor QC was performed independently on each sample. Cells expressing fewer than 200 or more than 10,000 genes, and cells with mitochondrial gene content exceeding 20%, were removed. Genes detected in fewer than three cells per sample were excluded. Remaining counts were log-normalized per cell. Gene expression profiles were aligned to a fixed vocabulary of 18,301 genes used by ConvergeCELL; genes absent from a given sample were zero-filled, and genes outside the vocabulary were dropped. For disease classification and explainability, MM was assigned a positive label (active myeloma) while SMM and MGUS were grouped together as a precursor/control class.

#### Sepsis

Bulk RNA-seq data were obtained from GSE185263, comprising 392 whole-blood samples (348 sepsis, 44 healthy controls) collected across five sites. Pre-processed gene-level counts and transcripts per million (TPM)-normalized matrices were downloaded from NCBI GEO, aligned to GRCh38.p13 by the depositing authors. Gene expression profiles were aligned to the same 18,301-gene vocabulary used by ConvergeCELL, and bulk vectors were treated as pseudo-single-cell profiles, enabling direct application of the model without architectural modification. The classification task was framed as disease-versus-healthy: positive class = High SOFA (≥ 5, n = 82), negative class = Low SOFA (< 2, n = 138) combined with healthy controls (n = 44); samples with intermediate SOFA (2 ≤ SOFA < 5) and missing SOFA were excluded, yielding 264 classifiable samples.

Classification and biomarker attribution were performed using a fully zero-shot transfer protocol. The pretrained, frozen BulkEmbedder was applied to pseudobulk profiles of all 3,202 training and 348 validation patients in the scBaseCount corpus, yielding 512-dimensional embeddings. An XGB classifier was then trained on these embeddings against the 10-class disease-family labels, with hyperparameters tuned by grid search on validation weighted-F1 and balanced sample weights to account for class imbalance. The resulting tuned model was applied directly to the sepsis cohort without further training, fine-tuning, or cross-validation. A sample was classified as positive if its predicted argmax was non-healthy. For classification performance evaluation, AUROC was computed using 1 - P(healthy_control) as the continuous score.

For IG attribution, a parallel linear head (nn.Linear(512, 10)) was trained on the same scBaseCount embeddings as a Captum-compatible surrogate, and IG was run end-to-end on the (frozen encoder + linear head) for 25 correctly-classified High-SOFA target samples (predicted argmax ∈ {immune_inflammatory, metabolic_vascular}), using 10 random correctly-predicted-healthy baselines per target, 200 integration steps, and mu_macro cohort aggregation. The attribution target was the scalar P(immune_inflammatory) + P(metabolic_vascular), computed via a softmax-summing wrapper.

### Baseline methods

#### Differential expression analysis

For single-cell data, differential expression was performed using a pseudobulk approach to ensure valid donor-level statistical inference. Raw counts were aggregated across all cells of the disease-relevant cell type (T4 cells for SLE, PC for MM) per donor, producing one expression profile per donor. Pseudobulk profiles were analyzed using DESeq2^50^ with default parameters. Genes were ranked by Benjamini-Hochberg adjusted p-value. Our donor-aware approach avoids the pseudoreplication inherent in cell-level tests, which treat millions of cells as independent observations despite their nested structure within donors. For comparison, cell-level Wilcoxon rank-sum results computed using scanpy’s rank_genes_groups function are provided in Supplementary Fig. 5. For bulk RNA-seq data (sepsis cohort), DESeq2 was applied directly to the raw count matrix using the same default parameters, with target = High SOFA (≥ 5) and reference = (Low SOFA < 2) ∪ healthy controls. Samples with 2 ≤ SOFA < 5 and missing-SOFA samples were excluded.

#### Standard ML baseline

XGB classifiers were trained on pseudobulk expression profiles generated by mean-aggregating single-cell expression vectors over the disease-relevant cell type per patient (T4 cells for SLE, PC for MM), to align with the cell-type-specific scope used for explainability in ConvergeCELL and PaSCient. For sepsis, which is a bulk RNA-seq dataset, this distinction does not apply. Balanced class weights were computed via compute_sample_weight(“balanced”) to account for class imbalance. The model used log-loss as the evaluation metric. PCA dimensionality reduction was applied to the pseudobulk profiles prior to training, with the transform fit on training data and applied to both train and validation splits. Hyperparameters were optimized using Bayesian search. For disease-associated gene ranking, TreeSHAP values were computed per sample over PCA components using the trained XGB model, then back-projected to gene space by multiplying the per-sample SHAP matrix by the PCA loading matrix. Mean absolute SHAP value per gene across samples was used as the gene-level importance score. The sample subset used for TreeSHAP was matched per disease to balance attribution quality with sample availability: SLE used true-label disease samples (disease_state ∈ {managed, flare}, n = 162) without a prediction filter, since the SLE Standard ML model predicted “healthy” for nearly all samples and applying the correctly-classified filter left too few targets; MM used correctly-predicted samples (predicted argmax = oncological, n = 42); sepsis used correctly-predicted High-SOFA samples (SOFA ≥ 5 ∩ predicted argmax ∈ {immune_inflammatory, metabolic_vascular}, n = 62). For sepsis specifically, the TreeSHAP target was P(immune_inflammatory) + P(metabolic_vascular), computed via SHAP additivity (shap[1] + shap[3] per sample). Genes were ranked by descending |importance| - the same convention as IG.

#### PaSCient baseline

PaSCient is a patient-level single-cell foundation model that classifies disease state from unordered bags of cells. The pretrained multilabel checkpoint (30.8M parameters) was obtained upon request from the authors. To ensure independence from ConvergeCELL preprocessing, PaSCient was evaluated using per-sample h5ad files containing raw counts and the full gene set (34,849 genes for MM; 30,172 for SLE), prior to any ConvergeCELL-specific gene filtering. Genes were mapped to PaSCient’s 28,231-gene vocabulary; 24,325 genes overlapped for MM and 19,919 for SLE, with unmapped positions zero-filled. Raw counts were normalized to a target sum of 10,000 counts per cell and log1p-transformed, following PaSCient’s own preprocessing protocol. Cells were padded or subsampled to the model’s fixed input size of 1,500 cells per sample. For each sample, all cells (irrespective of cell type) were provided as input, and a forward pass produced a nine-class probability distribution via softmax. Binary disease-versus-healthy predictions were derived by classifying any sample whose argmax prediction was not the healthy class (index 4) as diseased. The probability of disease was defined as 1 − P(healthy); for SLE, P(MS) was not used as the classification score because no samples received MS as the argmax class, so 1 − P(healthy) - which captures predictions spread across non-healthy classes - provides a fairer signal. IG attributions were computed following the methodology described in the PaSCient paper. A differential forward wrapper was constructed that returns the scalar logit(disease) − logit(healthy), matching the authors’ ForwardDiffModel implementation. For MM, the target class was “multiple myeloma” (index 7); for SLE, which is absent from PaSCient’s label space, MS (index 8) was used as a proxy for immune-inflammatory disease. IG was computed using the Captum library with PaSCient’s default parameters. All samples were used for IG rather than the correctly-classified subset used for ConvergeCELL, because PaSCient correctly predicted only a small fraction of samples in both cohorts - too few for stable cohort-level attribution. We note that the PaSCient repository provides limited documentation, and our implementation reflects our best effort to apply the model in the most consistent manner; reproducibility was a recurring challenge.

### Integrated Gradients

Gene-level attribution scores were computed using IG via the Captum library. For each target sample (correctly classified disease patients, up to 25 per disease label to balance computational cost with cohort representation), ten reference baselines were randomly selected from the control class. The attribution target was the predicted probability of the corresponding ConvergeCELL disease family class: immune_inflammatory for SLE; oncological for MM and immune_inflammatory + metabolic_vascular for sepsis. Restricting to correctly classified patients ensures that IG attributions reflect genes driving accurate disease predictions rather than noise from misclassified samples, representing the biological reference state against which attributions are computed. IG was run with 200 interpolation steps using Gauss–Legendre quadrature for numerical integration. For each target–reference pair, IG produced a per-cell, per-gene attribution matrix. Attributions were first averaged across baselines, then across cells within each sample to yield a signed per-gene attribution vector per patient. Cohort-level gene importance was computed as the macro-average of signed patient attributions (mu_macro), and genes were ranked by the absolute value of this cohort-level score (|mu_macro|) - preserving directional consistency such that genes must consistently push the prediction in one direction across patients to rank highly. Cell-type-specific attributions were computed by restricting to the disease-relevant cell type (PC for MM, T4 cells for SLE). Convergence deltas were tracked as a quality metric. Statistical significance of gene-level attributions was assessed via two-tailed hypothesis testing (FDR < 0.05), identifying genes with consistently non-zero attribution across patients.

### Hypothesis generation workflow

The hypothesis generation workflow uses Claude Opus 4.6 (Anthropic) connected to biomedical knowledge bases via the MCP, a standardized interface for LLM tool access. For each candidate gene, the LLM queries PubMed, ClinicalTrials.gov via dedicated MCP servers, and receives Open Targets disease-gene association scores, drug development phase, and novelty indicators as structured context. Multiple prompt templates (standard, elaborate, and confidence-calibrated) guide the LLM to classify each gene’s evidence level and generate mechanistic hypotheses linking gene function to disease pathology. The workflow leverages the LLM’s existing capabilities for biomedical reasoning and literature synthesis rather than implementing a custom agent architecture; the contribution is the structured integration of ConvergeCELL’s gene rankings with knowledge base access and prompt design optimized for therapeutic hypothesis generation.

### Open Targets validation

Disease-gene associations were obtained from the Open Targets Platform using the GraphQL API, which provides per-target association scores aggregated across all evidence sources (genetic associations, expression, literature, clinical precedence, and others). For each disease, the official overall association score - a harmonic-sum weighted aggregate computed by Open Targets across all data sources - was used as the gene-level relevance measure. Statistically significant disease-gene associations were identified using a z-score threshold of 1.5 on the overall association score distribution. To ensure a fair comparison across methods with different gene vocabularies, all gene rankings were restricted to the intersection of the ConvergeCELL and PaSCient gene vocabularies (16,040 shared genes). Two gene-level metrics were computed at K = 25, 50, 100, and 200: (1) Gene Recall@K, defined as the fraction of Open Targets-significant genes appearing within each method’s top-K ranked genes; and (2) Mean OT score@K, defined as the arithmetic mean of Open Targets overall association scores across each method’s top-K genes. Statistical significance for both metrics was assessed against a permutation null distribution (1,000 random gene orderings). Pathway enrichment was performed on the top 500 ranked genes per method using over-representation analysis (Fisher’s exact test, FDR < 0.05) against the full Reactome pathway collection (MSigDB c2 Reactome, v2024.1.Hs). To highlight enrichment for canonical disease biology, we additionally assembled a curated, disease-specific Reactome reference set per disease; pathways from each method’s top hits that were also members of the corresponding curated reference set are highlighted in red in Fig. 5d-f and Supplementary Figs. 9-11, using over-representation analysis with Fisher’s exact test (FDR < 0.05); curated reference pathway sets were assembled per disease from Reactome based on canonical disease pathophysiology.

### Approved-drug-target rank analysis

To assess whether each model’s gene ranking surfaces clinically validated drug targets, we compiled the set of approved drugs for each disease from Open Targets (GraphQL API; disease.drugAndClinicalCandidates endpoint), retaining only drugs at maximum clinical stage APPROVAL. For each approved drug, a single primary mechanism-of-action target gene was curated through a deep-research workflow that filtered for drugs with a well-established, dominant target gene supported by clinical literature and FDA labeling, and excluded drugs whose mechanism is mediated through multiple targets without a clear primary. Disease IDs used were MONDO_0007915/EFO_0002430 (SLE) and EFO_0001378 (MM), yielding 4 and 10 approved drug-primary target pairs after curation. For each (disease, drug, model) triple, we recorded the rank of the curated primary target gene within the shared gene space together with DE (13,077 genes for SLE; 11,198 for MM). Aggregate per-model performance was reported as the mean rank across drugs. Sepsis was omitted as it lacks strong disease-mechanism-targeted approved therapies.

### Statistical analysis

#### Classification metrics on the validation set

Pairwise differences in weighted F1 between ConvergeCELL and each baseline on the study-aware held-out validation set (Supplementary Fig. 1c) were assessed by 1,000-resample paired bootstrap; reported p-values are two-sided.

#### AUROC and PR-AUC

Bootstrap 95% confidence intervals for AUROC were computed by resampling patients with replacement (1,000 iterations) and computing AUROC per resample. Pairwise AUROC comparisons between ConvergeCELL and each baseline were assessed using the DeLong test, which provides an asymptotic test for the difference between two correlated AUROCs evaluated on the same dataset. PR-AUC was computed using sklearn.metrics.average_precision_score, with bootstrap confidence intervals and pairwise significance assessed by paired bootstrap. Multiple comparisons were Bonferroni-corrected.

All statistical tests were performed in Python using scipy and statsmodels.

## Acknowledgements

The authors thank Prof. Eran Segal, Prof. David Burstein, and Glen Weiss, MD, for their critical reading of the manuscript and insightful comments. The authors also thank Shira Levy, MD, for assistance with curating scBaseCount data for disease family annotation, the Genentech team for sharing the PaSCient model weights, and Iris Burstein for designing the platform schematic.

## Author contributions

N.S. conceptualized the study, developed the methodology, built the models, conducted experiments, supervised the project, and contributed to writing the manuscript. D.M., M.S., and G.L. contributed to conceptualization and methodology, built the models, conducted experiments, and contributed to writing the manuscript. I.W. contributed to conceptualization and supervised the project.

## Model availability

The pretrained ConvergeCELL patient representation model and bulk distillation module are publicly available on Hugging Face (huggingface.co/ConvergeBio) under the Apache 2.0 license, deposited as **virtual-cell-patient** (huggingface.co/ConvergeBio/virtual-cell-patient) and **virtual-cell-distil-bulk** (huggingface.co/ConvergeBio/virtual-cell-distil-bulk). Example datasets are provided as **virtual-cell-patient-example** and **virtual-cell-distil-bulk-example** under the same organization.

## Data availability

The datasets generated and analyzed during the current study will be available upon publication of the peer-reviewed version of this article. Six Supplementary Data files accompany this manuscript: The approved drug-target curation table (Supplementary Data 1), per-method top-200 gene rankings (Supplementary Data 2), curated Reactome pathway reference sets (Supplementary Data 3), and hypothesis-generation agent outputs for SLE, MM, and sepsis (Supplementary Data 4-6). Descriptions of all Supplementary Data files (Supplementary Data 1-6) are provided in the Supplementary Data Information.

## Code availability

The code used in this study will be made publicly available upon publication of the peer-reviewed version of this manuscript.

## Abbreviation table

Abbreviation Full term

AUROC: Area under the receiver operating characteristic curve
BM: Bone marrow
C2S: Cell2Sentence
DE: Differential expression
FDR: False discovery rate
IG: Integrated gradients
SOFA: Sequential Organ Failure Assessment
LLM: Large language model
MCP: Model Context Protocol
MGUS: Monoclonal gammopathy of undetermined significance
MLP: Multi-layer perceptron
MM: Multiple myeloma
MS: Multiple sclerosis
PBMC: Peripheral blood mononuclear cell
PCA: Principal component analysis
PR-AUC: Precision-recall area under the curve
T4: CD4+T
PC: Plasma cells
PReLU: Parametric rectified linear unit
QC: Quality control
ROC: Receiver operating characteristic

