## Supplementary Data Information for "ConvergeCELL: An end-to-end platform from patient transcriptomics to therapeutic hypotheses"

| | |
|---|---|
| **Supplementary Data 1** | Approved drugs for SLE and MM compiled from Open Targets, with all annotated target genes per drug and the curated primary mechanism-of-action target gene used in Fig. 4 (4 SLE and 10 MM drug–target pairs). |
| **Supplementary Data 2** | Per-method top-200 gene rankings for SLE, MM, and sepsis (ConvergeCELL, PaSCient, pseudobulk DE, standard ML), with method-specific scores and corresponding Open Targets overall association scores. |
| **Supplementary Data 3** | Curated, disease-specific Reactome pathway reference sets used for the over-representation analysis in Fig. 5d-f and Supplementary Figs. 9-11, listed per disease (SLE, MM, sepsis) with Reactome pathway identifiers. |
| **Supplementary Data 4** | Hypothesis-generation outputs for the top-ranked genes in SLE, including direct/indirect/no-evidence classification and structured mechanistic hypotheses (Fig. 5a). |
| **Supplementary Data 5** | Hypothesis-generation outputs for the top-ranked genes in MM, including evidence classification and mechanistic hypotheses (Fig. 5b). |
| **Supplementary Data 6** | Hypothesis-generation outputs for the top-ranked genes in sepsis, including evidence classification, mechanistic hypotheses, and the illustrative MMP25 case shown in Supplementary Fig. 13. |
