## Supplementary Materials for "ConvergeCELL: An end-to-end platform from patient transcriptomics to therapeutic hypotheses"

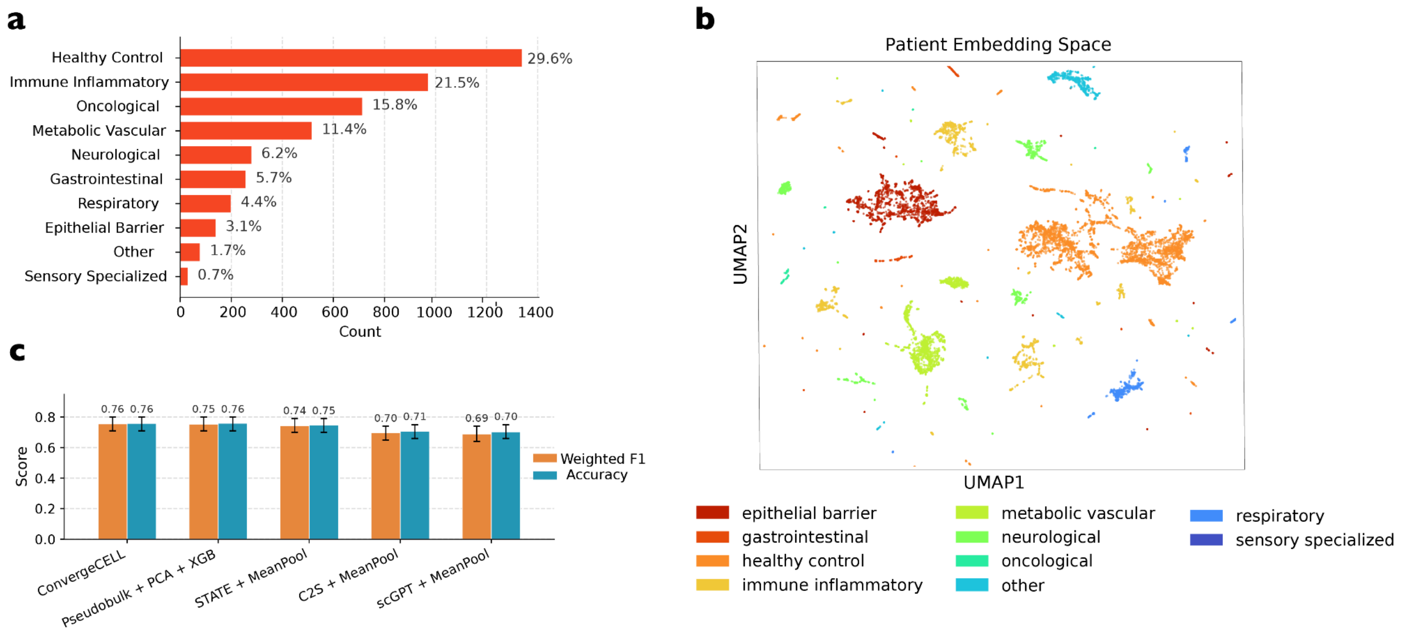


**Supplementary Figure 1. Training data composition and patient-level classification performance. (a)** Distribution of 4,479 patient samples across nine disease families and a healthy control category in the ConvergeCELL training corpus, derived from scBaseCount and curated to include primary human single-cell transcriptomic samples spanning hundreds of diseases. Healthy controls comprise 29.6% of the corpus and disease samples 70.4%. **(b)** UMAP projection of ConvergeCELL patient embeddings colored by disease family. **(c)** Patient-level classification performance (weighted F1 and accuracy) on a study-aware held-out validation set, comparing ConvergeCELL against four baselines: Pseudobulk + PCA + XGB, STATE + MeanPool, C2S + MeanPool, and scGPT + MeanPool. Error bars represent 95% confidence intervals from 1,000 bootstrap resamples of the validation set. Pairwise differences in weighted F1 between ConvergeCELL and each baseline were not statistically significant (paired bootstrap, all $p > 0.05$).

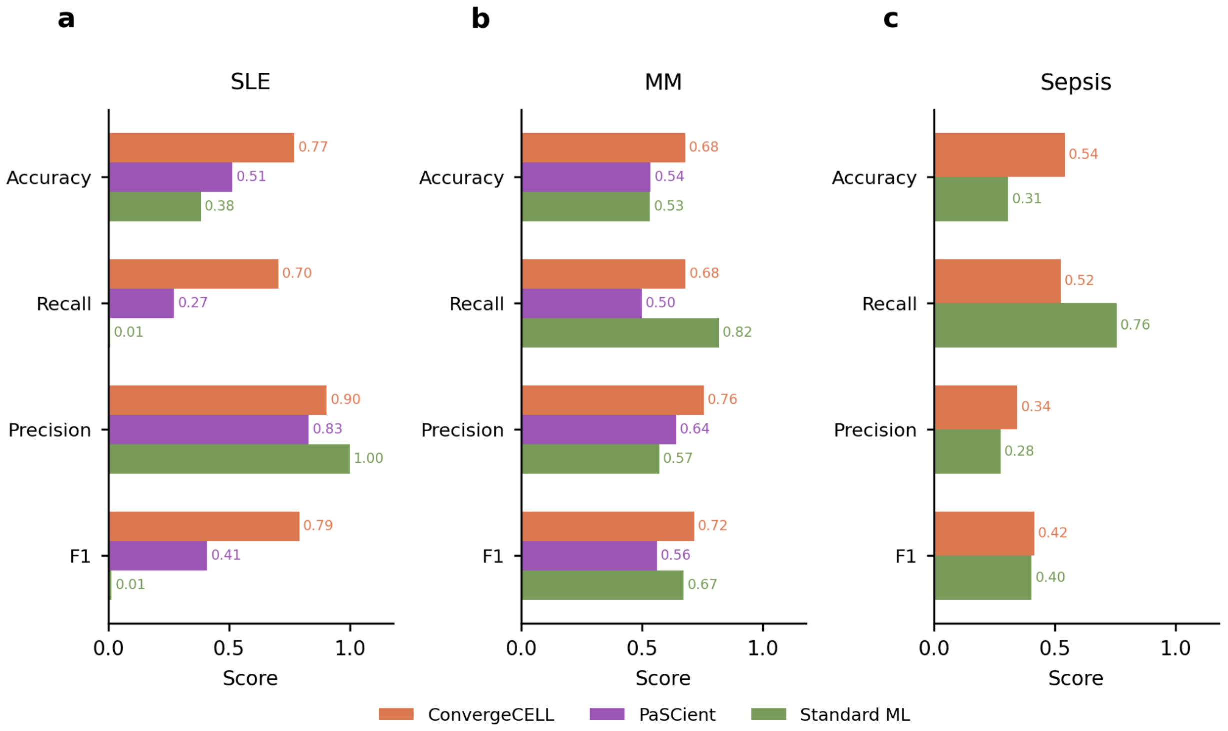


**Supplementary Figure 2. Classification performance metrics across methods and diseases on held-out cohorts.** Accuracy, recall, precision, and F1 score for ConvergeCELL (orange), PaSCient (purple), and the standard ML baseline (green) on **(a)** SLE (flare and managed versus healthy controls), **(b)** MM versus its precursor states (SMM and MGUS), and **(c)** sepsis (High SOFA ≥ 5 versus Low SOFA < 2 combined with healthy controls). PaSCient is shown for SLE and MM only (single-cell diseases).

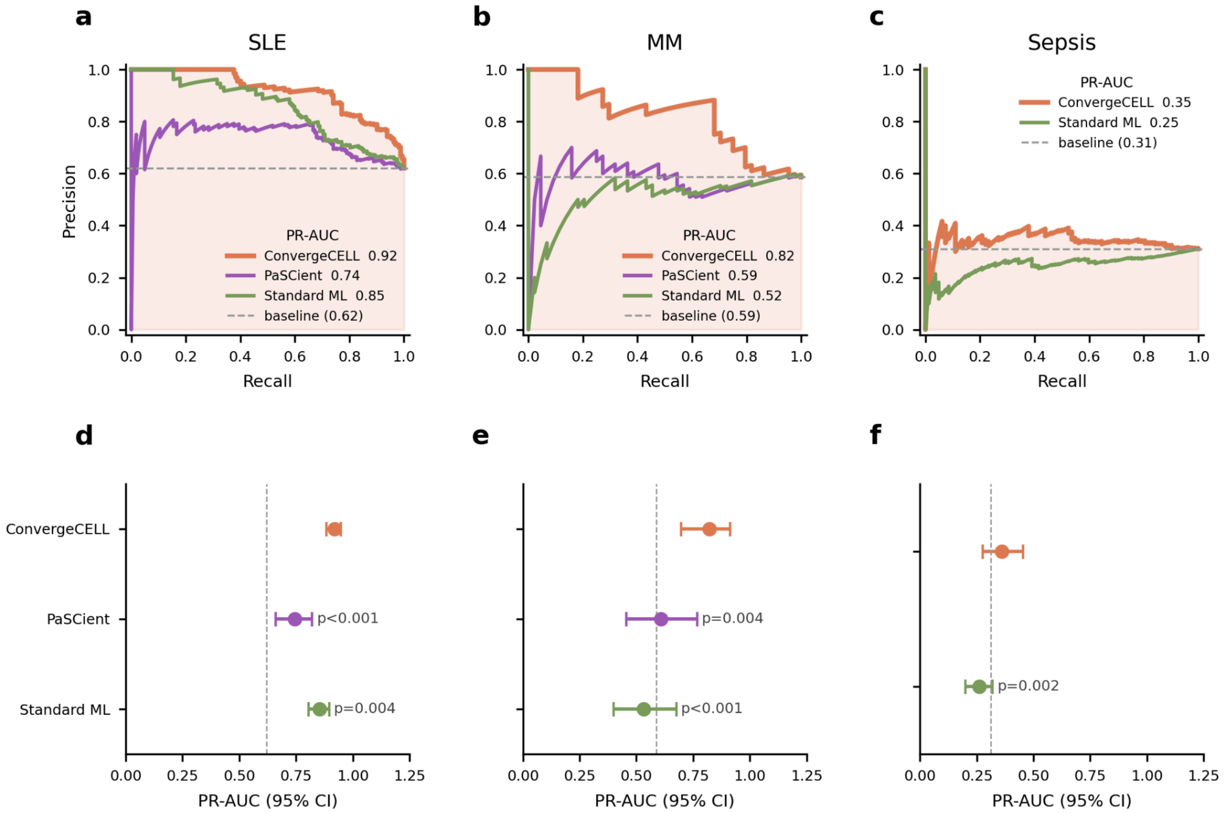


**Supplementary Figure 3. Precision-recall analysis across methods and diseases on held-out cohorts. (a-c)** Precision-recall curves for ConvergeCELL (orange), PaSCient (purple, single-cell diseases only), and the standard ML baseline (green) on **(a)** SLE, **(b)** MM, and **(c)** sepsis. PR-AUC values shown in legends; dashed lines indicate positive class prevalence (random baseline). **(d-f)** Bootstrap 95% confidence intervals for PR-AUC (1,000 resamples) with Bonferroni-corrected pairwise significance testing between ConvergeCELL and each baseline.

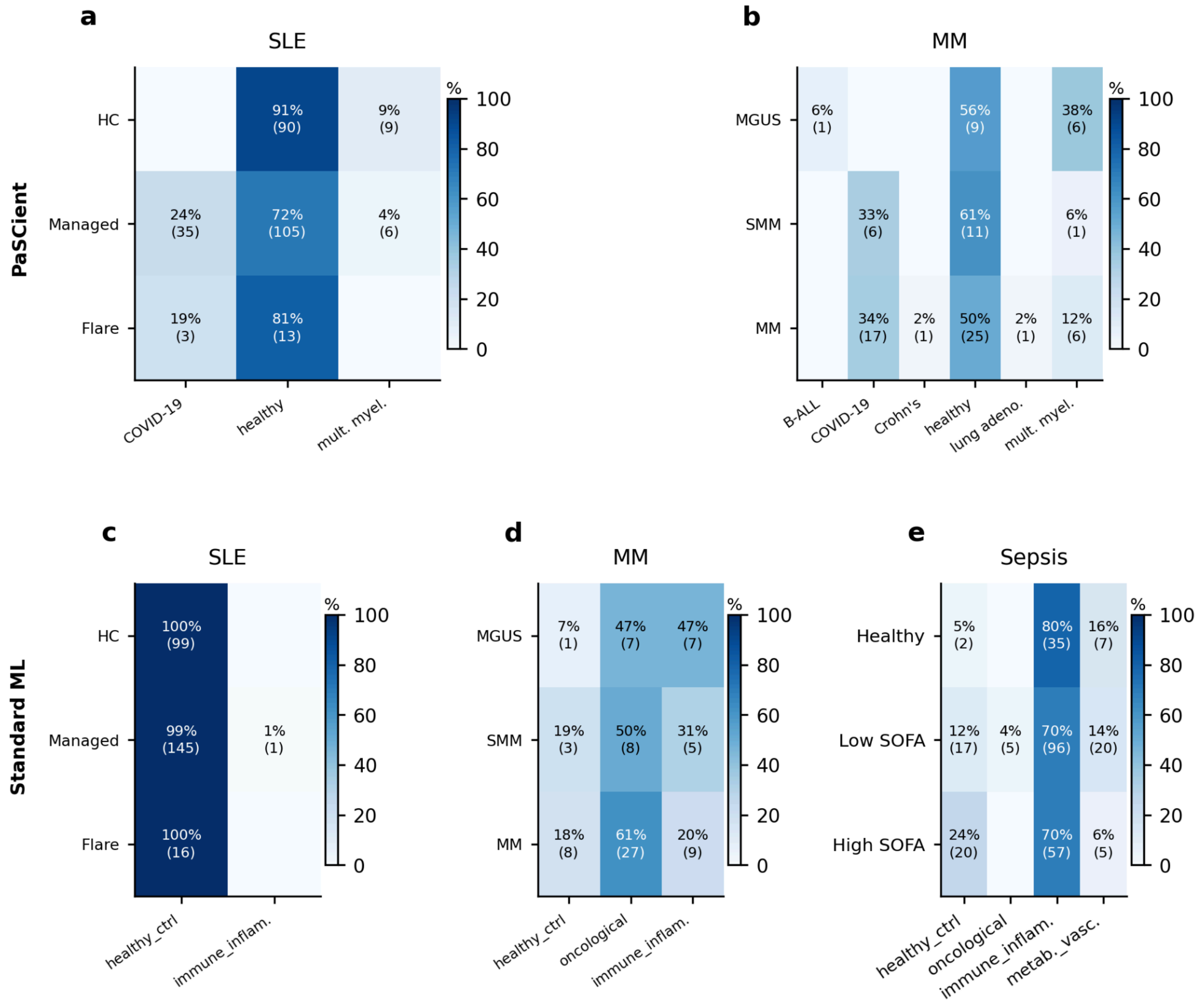


**Supplementary Figure 4. Confusion matrices for baseline methods.** Predicted class (columns) stratified by clinical stage or severity group (rows) for PaSCient on **(a)** SLE and **(b)** MM, and the standard ML baseline on **(c)** SLE, **(d)** MM, and **(e)** sepsis. Values indicate the percentage and absolute number of samples assigned to each predicted class. PaSCient is shown for single-cell diseases only.

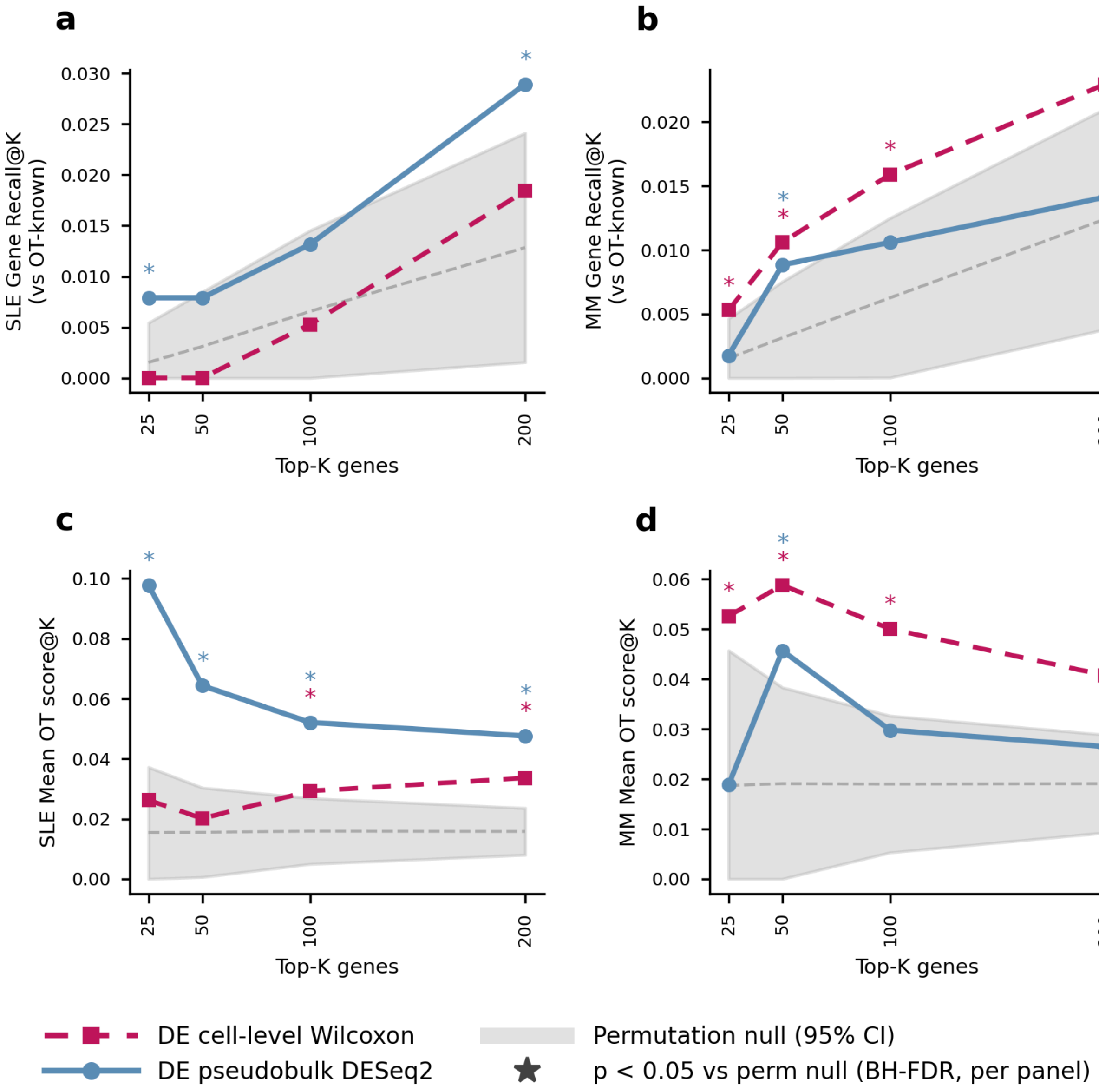


**Supplementary Figure 5. Comparison of cell-level and pseudobulk differential expression for disease-associated gene recovery. (a-b)** Gene Recall@K and (c-d) Mean OT score@K for cell-level Wilcoxon DE (red, dashed) and donor-aware pseudobulk DESeq2 (blue, solid) across **(a,c)** SLE in T4 cells and **(b,d)** MM in PC, evaluated at K = 25, 50, 100, 200. Pseudobulk profiles were generated by aggregating raw counts across all cells of the disease-relevant cell type per donor and analyzed with DESeq2; cell-level Wilcoxon DE was computed across individual cells of the same cell type using scanpy's rank_genes_groups (Wilcoxon rank-sum test). Grey shading indicates the 95% confidence interval of a permutation null (1,000 random gene orderings); asterisks denote $p < 0.05$ versus the permutation null by z-test (Benjamini-Hochberg FDR-corrected within each panel).

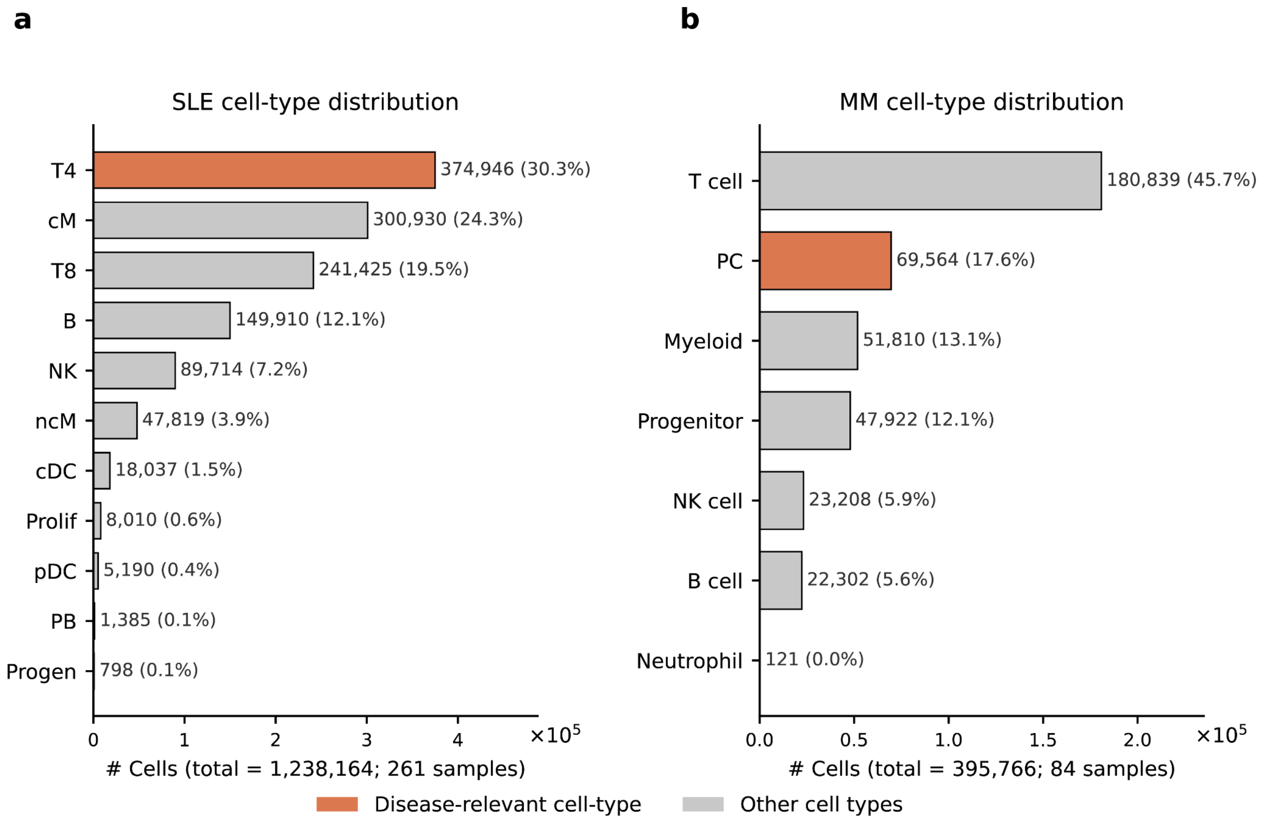


**Supplementary Figure 6. Cell-type composition of the SLE and MM held-out cohorts.** Distribution of cells across major cell populations in the **(a)** SLE PBMC cohort (1,238,164 cells from 261 samples) and **(b)** MM bone marrow cohort (395,766 cells from 84 samples). Cell counts and percentages of total cells are shown next to each bar. The disease-relevant cell type used for explainability analyses in the main text is highlighted in orange (T4 cells for SLE; PC for MM); all other cell populations are shown in grey. Cell-type annotations are taken from the original publications.

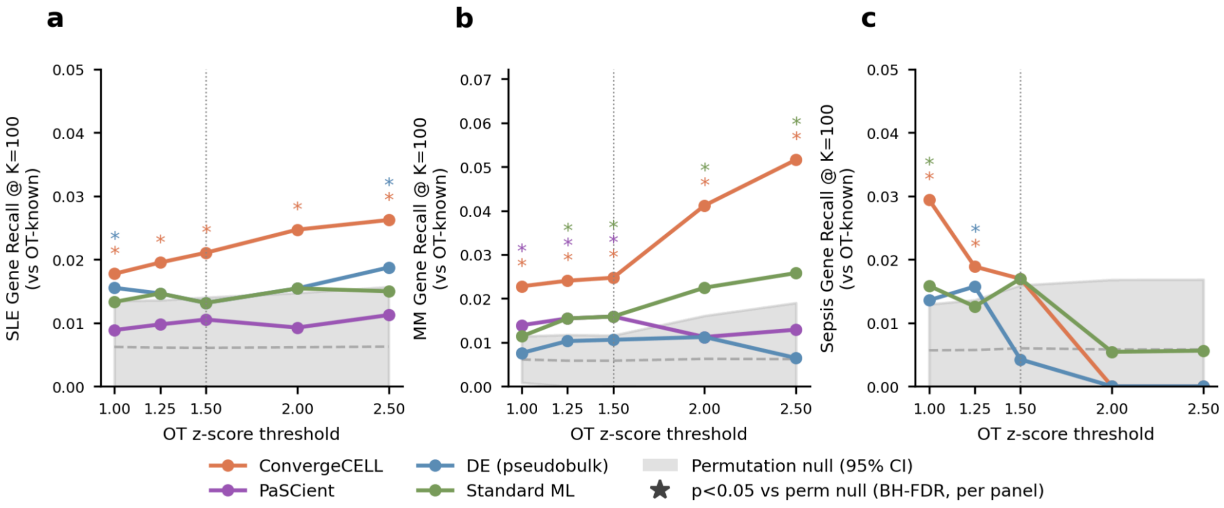


**Supplementary Figure 7. Sensitivity of Gene Recall to Open Targets significance threshold.** Gene Recall@K=100 as a function of the z-score threshold used to define significant disease-gene associations from Open Targets overall association scores, for **(a)** SLE, **(b)** MM, and **(c)** sepsis. Thresholds tested: 1.0, 1.25, 1.5 (used in main manuscript; shown as dotted vertical line), 2.0, and 2.5. Results shown for ConvergeCELL (orange), PaSCient (purple, single-cell diseases only), pseudobulk DE (blue), and the standard ML baseline (green). Grey shading indicates the 95% confidence interval of a permutation null. Asterisks denote $p < 0.05$ versus the permutation null by z-test (Benjamini-Hochberg FDR-corrected within each panel). Mean OT score is omitted from this sensitivity analysis as it uses the continuous Open Targets association score across all genes and is therefore independent of the significance threshold.

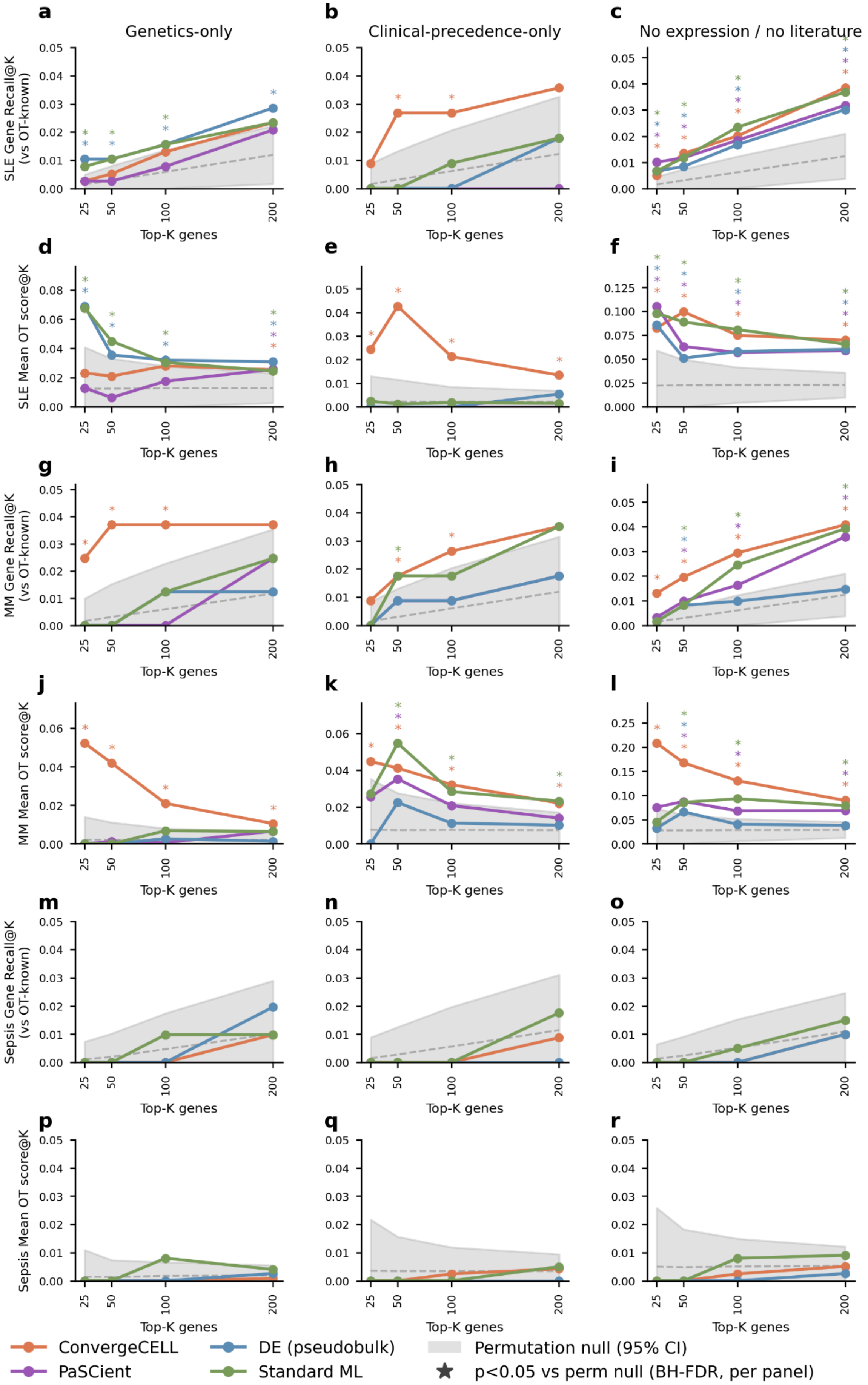

a
Genetics-only
b
Clinical-precedence-only
c
No expression / no literature
SLE Gene Recall@K (vs OT-known)
SLE Mean OT score@K
MM Gene Recall@K (vs OT-known)
MM Mean OT score@K
Sepsis Gene Recall@K (vs OT-known)
Sepsis Mean OT score@K
Top-K genes
ConvergeCELL
PaSCient
DE (pseudobulk)
Standard ML
Permutation null (95% CI)
p<0.05 vs perm null (BH-FDR, per panel)

**Supplementary Figure 8. Gene recovery under Open Targets evidence stratifications that exclude expression and literature sources.** Top-K gene recovery against three Open Targets sub-scores that exclude potentially circular evidence types: genetics-only **(a, d, g, j, m, p)**, clinical-precedence-only **(b, e, h, k, n, q)**, and a composite excluding both expression and literature evidence **(c, f, i, l, o, r)**. Rows show Gene Recall@K and Mean OT score@K for SLE (a-f), multiple myeloma (g-l), and sepsis (m-r). Each panel evaluates the top-K genes (K = 25, 50, 100, 200) against disease-associated genes defined under the corresponding OT sub-score (z-score >= 1.5). Lines compare ConvergeCELL (orange), PaSCient (purple), pseudobulk differential expression (blue), and the Standard ML baseline (green). The grey band shows the 95% confidence interval of a permutation null obtained by sampling random gene sets of equal size from the shared gene universe; the dashed line marks the null mean. Asterisks indicate Top-K values at which a method exceeds the permutation null at $p < 0.05$ (Benjamini-Hochberg FDR-corrected within each panel), colored by method.

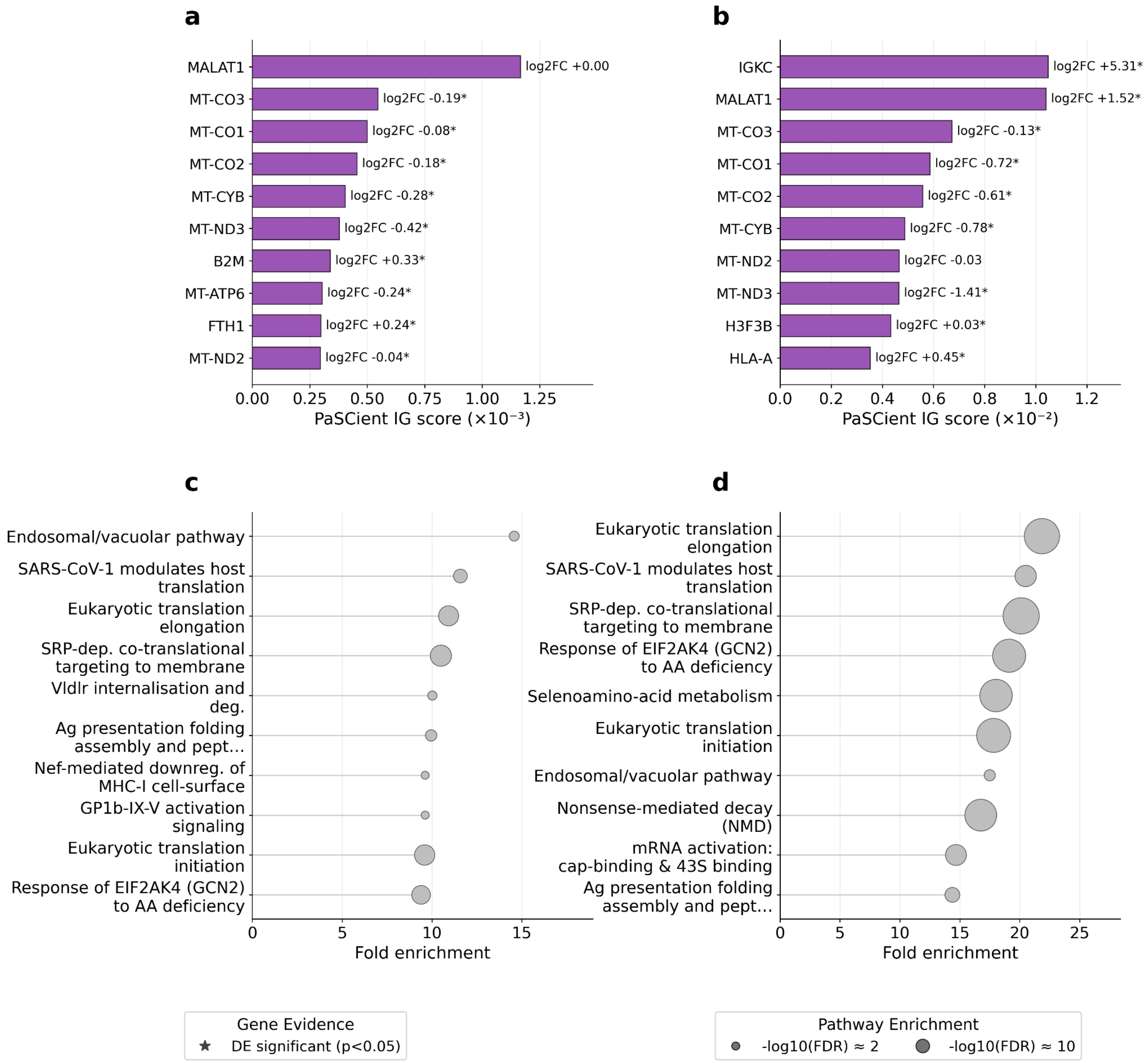


**Supplementary Figure 9. PaSCient top-attributed genes and pathway enrichments. (a-b)** Top 10 PaSCient-attributed genes ranked by IG score for **(a)** SLE in T4 cells and **(b)** MM in PC. Asterisks next to log2 fold-change values denote genes also reaching significance by DE ($p < 0.05$); log2 fold-change values are shown to the right of each bar. **(d-e)** Top enriched Reactome pathways from over-representation analysis on each disease's top-K PaSCient gene list (K = 500) for **(d)** SLE and **(e)** MM. Dot size encodes -log10(FDR).

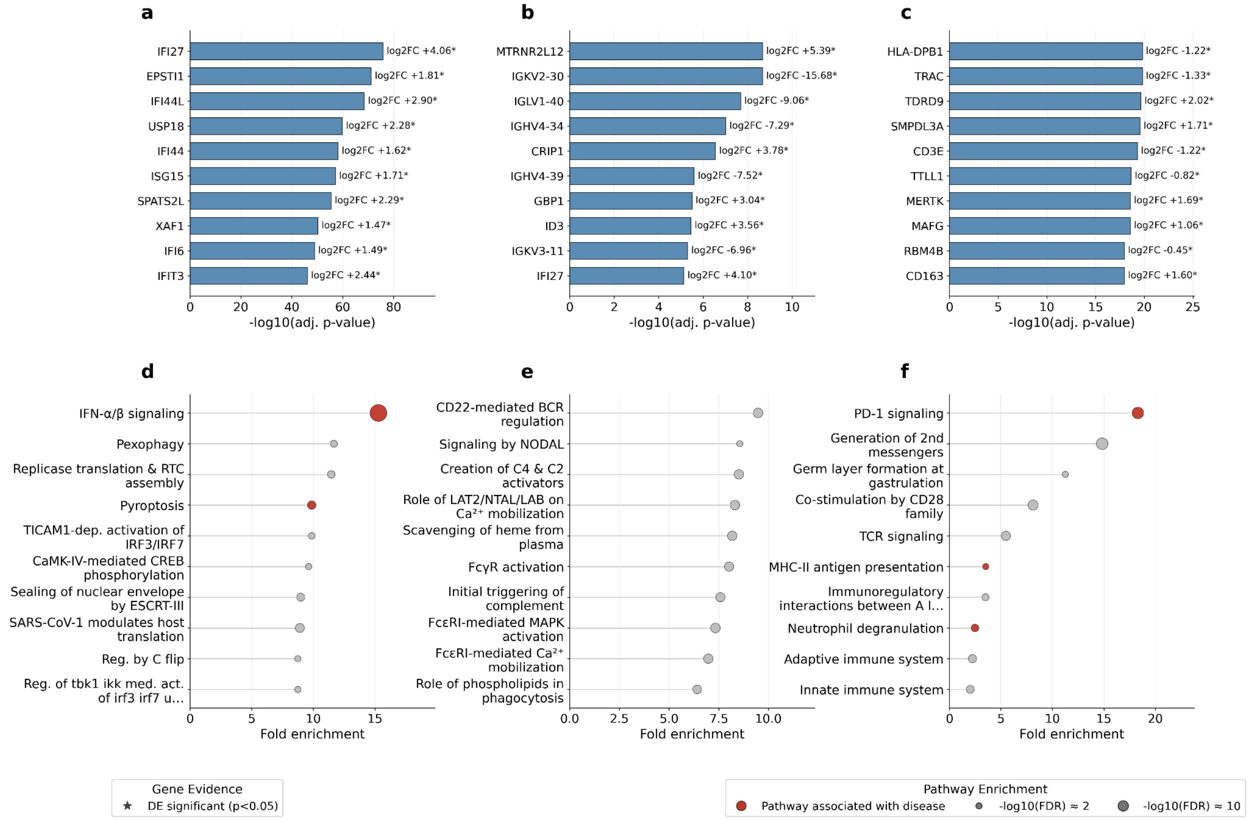


**Supplementary Figure 10. Pseudobulk DE top-ranked genes and pathway enrichments. (a-c)** Top 10 DE-attributed genes ranked by -log10(adjusted p-value) score for **(a)** SLE in T4 cells, **(b)** MM in PC, and **(c)** sepsis in whole blood. Asterisks next to log2 fold-change values denote genes also reaching significance by DE ($p < 0.05$); log2 fold-change values are shown to the right of each bar. **(d-f)** Top enriched Reactome pathways from over-representation analysis on each disease's top-K DE gene list (K = 500) for **(d)** SLE, **(e)** MM, and **(f)** sepsis. Dot size encodes -log10(FDR); pathways also represented in the curated Reactome reference set for each disease are shown in red.

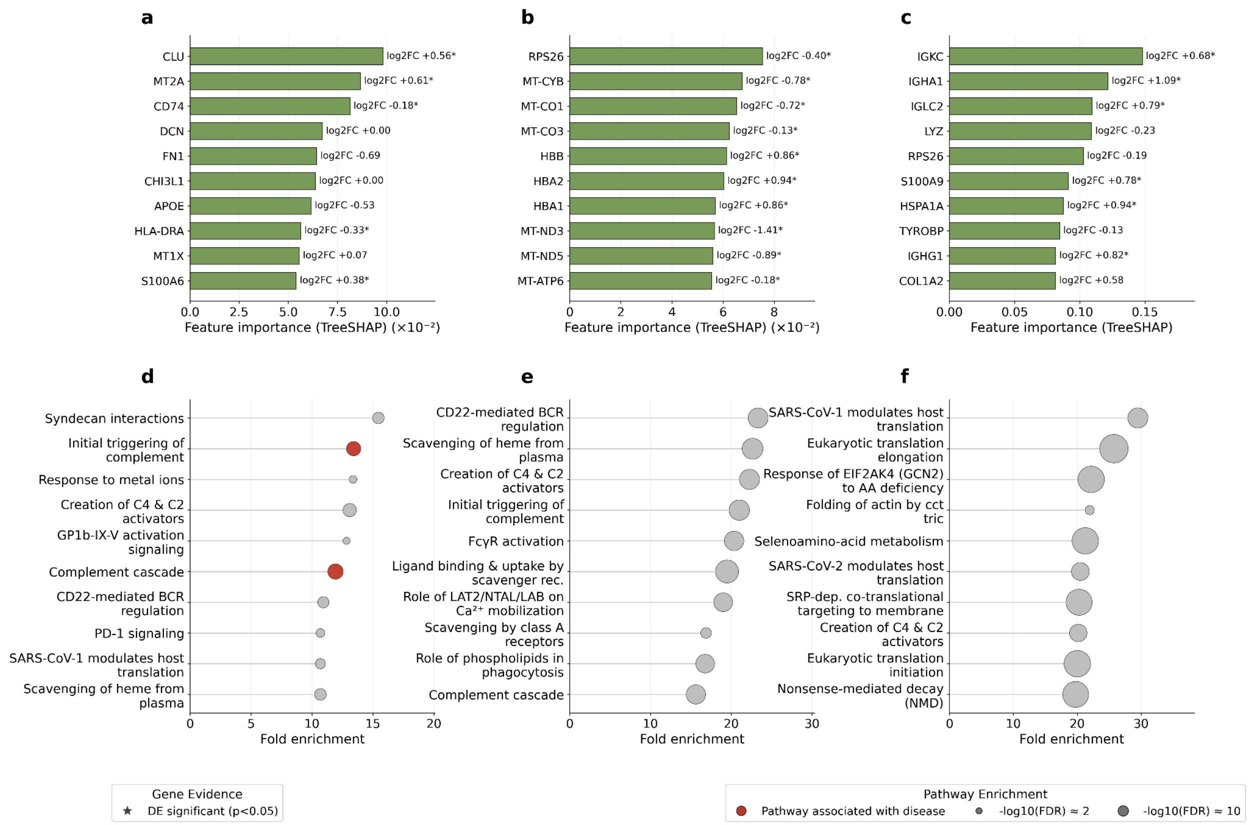


**Supplementary Figure 11. Standard ML baseline top-ranked genes and pathway enrichments. (a-c)** Top 10 standard ML-attributed genes ranked by TreeSHAP attribution score for **(a)** SLE in T4 cells, **(b)** MM in PC, and **(c)** sepsis in whole blood. Asterisks next to log2 fold-change values denote genes also reaching significance by DE ($p < 0.05$); log2 fold-change values are shown to the right of each bar. **(d-f)** Top enriched Reactome pathways from over-representation analysis on each disease's top-K standard ML gene list (K = 500) for **(d)** SLE, **(e)** MM, and **(f)** sepsis. Dot size encodes -log10(FDR); pathways also represented in the curated Reactome reference set for each disease are shown in red.

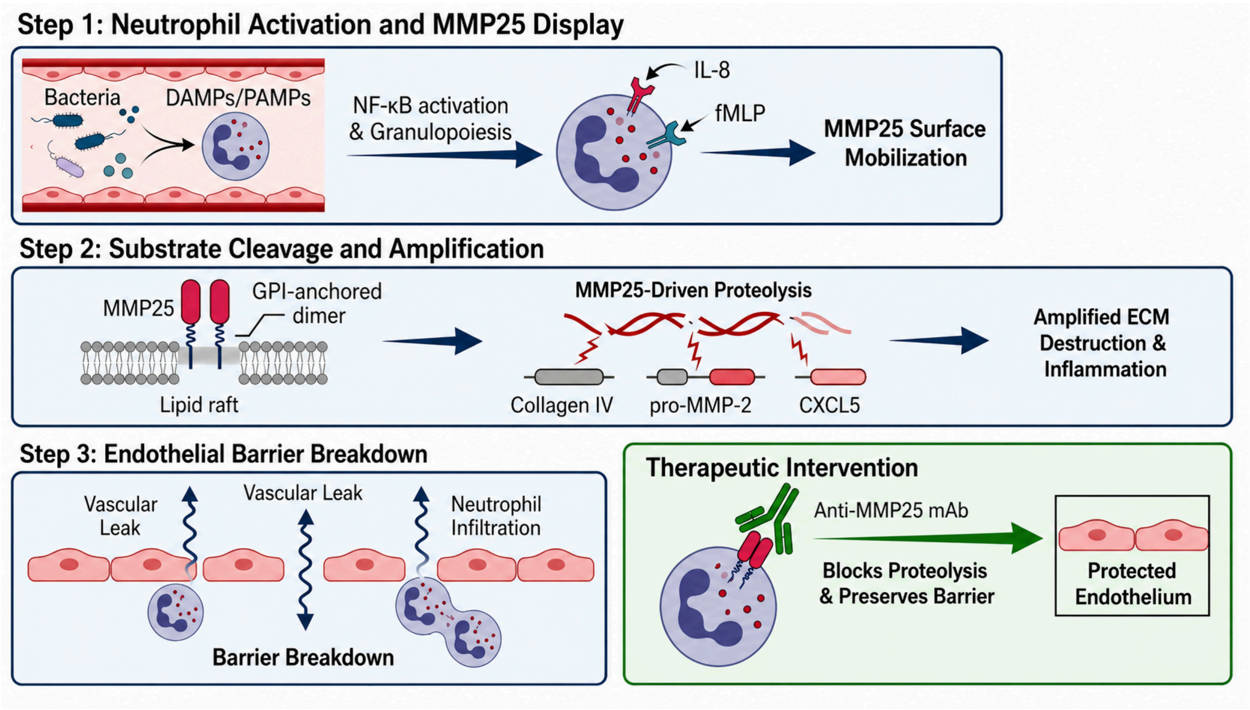


**Supplementary Figure 12. Mechanism-of-action hypothesis for MMP25 in sepsis, created by the ConvergeCELL hypothesis generation workflow.** MMP25 was identified as a mechanistic candidate through IG attribution on the bulk RNA-seq severity stratification task (ranked 49th). The workflow constructed a three-step mechanistic model linking MMP25 to sepsis pathophysiology. **(Step 1)** Bacterial infection and release of DAMPs/PAMPs activate NF-kB signalling and emergency granulopoiesis, producing activated neutrophils that mobilize MMP25 to the cell surface in response to chemotactic signals (IL-8, fMLP). **(Step 2)** MMP25, a GPI-anchored homodimer localized to lipid rafts, cleaves extracellular substrates including collagen IV, pro-MMP-2, and the chemokine CXCL5, generating a proteolytic cascade that amplifies extracellular matrix destruction and inflammation. **(Step 3)** Sustained MMP25-driven proteolysis disrupts endothelial barrier integrity, leading to vascular leak and further neutrophil infiltration. **(Therapeutic intervention, green)** The workflow proposes anti-MMP25 monoclonal antibody therapy to block proteolysis and preserve endothelial barrier function, noting that intervention timing is critical and should target the hyperinflammatory window (first 24-72 hours) before transition to the immunosuppressive phase. This hypothesis remains unvalidated and is presented as an illustrative example of the platform's output.

| **Indication** | **systemic lupus erythematosus (SLE)** | **multiple myeloma (MM)** | **Sepsis** |
|---|---|---|---|
| **Reference** | Perez, R. K. et al. Single-cell RNA-seq reveals cell type–specific molecular and genetic associations to lupus. *Science* **376**, eabf1970 (2022) | Foster, K. A. et al. Tumour-intrinsic features shape T cell differentiation through precursor to symptomatic multiple myeloma. *Nat. Commun.* **17**, 2400 (2026) | Baghela, A. et al. Predicting sepsis severity at first clinical presentation: the role of endotypes and mechanistic signatures. *EBioMedicine* **75**, 103776 (2022) |
| **Data type** | Single-cell transcriptomics | Single-cell transcriptomics | Bulk transcriptomics |
| **Sample type** | Peripheral blood mononuclear cell (PBMC) | Bone marrow | Whole-blood |
| **Disease family** | Immune-Inflammatory | Oncological | Immune-Inflammatory<br>Metabolic-Vascular |
| **Total cells** | 1,238,164 | 395,766 | NA |
| **Total donors** | 261 | 84 | 264 |
| **Donors by sub-classes** | HC: 99<br>Managed: 146<br>Flare: 16 | MM: 50<br>SMM: 18<br>MGUS: 16 | High SOFA: 82<br>Low SOFA: 138<br>Healthy: 44 |

**Supplementary Table 1.** Held out cohorts used for evaluation of model generalization.
