## Supplementary Data 4 for "ConvergeCELL: An end-to-end platform from patient transcriptomics to therapeutic hypotheses"

{
  "genes": [
    {
      "gene_name": "S100A9",
      "direct_evidence": "high",
      "indirect_evidence": "high",
      "is_stress_response_gene": false,
      "explanation": "S100A9 (with S100A8, forming calprotectin) is a well-established SLE severity biomarker. Serum and urine S100A8/A9 are elevated in SLE versus healthy controls, correlate with SLEDAI-2K, are particularly elevated in active lupus nephritis, and decrease with hydroxychloroquine response, especially in renal/cutaneous involvement. Mechanistically, S100A9 is an inflammatory alarmin/DAMP released by activated neutrophils and monocytes that drives TLR4/RAGE signaling, NETosis, and type I IFN amplification, all central SLE pathways; it is an immune-specific alarmin, not a generic stress gene.",
      "references": [
        "Wakiya R et al. (2019). Hydroxychloroquine modulates elevated expression of S100 proteins in systemic lupus erythematosus. Lupus.",
        "Sumova B et al. (2018). Circulating S100 proteins effectively discriminate SLE patients from healthy controls. Rheumatology International.",
        "Donohue SJ et al. (2020). Differential analysis of serum and urine S100 proteins in juvenile-onset systemic lupus erythematosus. Clinical Immunology.",
        "Kim JW et al. (2022). S100A8 in serum, urine, and saliva as a potential biomarker for systemic lupus erythematosus. Frontiers in Immunology."
      ]
    },
    {
      "gene_name": "PTPRCAP",
      "direct_evidence": "low",
      "indirect_evidence": "low",
      "is_stress_response_gene": false,
      "explanation": "PTPRCAP (CD45-associated protein) is a transmembrane regulator of CD45 (PTPRC) phosphatase activity that fine-tunes Lck-mediated TCR signaling in lymphocytes; it has no validated role as an SLE biomarker and no published association with SLEDAI, lupus nephritis activity, flare prediction, or treatment response. Indirect SLE relevance is limited to its general involvement in lymphocyte activation thresholds, which is peripheral compared to canonical SLE pathways (type I IFN, B-cell hyperactivation, complement). It is an immune signaling adaptor rather than a heat-shock/stress gene.",
      "references": [
        "Matsuda A et al. (1998). Disruption of lymphocyte function and signaling in CD45-associated protein-null mice. Journal of Experimental Medicine.",
        "Veillette A et al. (1999). The lymphocyte-specific protein tyrosine kinase Lck and CD45-associated protein. Immunological Reviews."
      ]
    },
    {
      "gene_name": "S100A8",
      "direct_evidence": "high",
      "indirect_evidence": "high",
      "is_stress_response_gene": false,
      "explanation": "S100A8 is a validated SLE severity biomarker: serum, urine, and salivary S100A8 are significantly elevated in SLE patients versus controls, correlate with SLEDAI-2K (with the strongest renal-component association), and are increased in active lupus nephritis including juvenile-onset SLE; HCQ treatment reduces S100A8 levels in line with clinical improvement. Mechanistically, S100A8 partners with S100A9 as calprotectin, a neutrophil/monocyte-derived alarmin engaging TLR4/RAGE that promotes NETosis, type I IFN amplification, and immune-complex-driven tissue inflammation. It is an immune-specific DAMP, not a generic stress-response gene.",
      "references": [
        "Kim JW et al. (2022). S100A8 in serum, urine, and saliva as a potential biomarker for systemic lupus erythematosus. Frontiers in Immunology.",
        "Wakiya R et al. (2019). Hydroxychloroquine modulates elevated expression of S100 proteins in systemic lupus erythematosus. Lupus.",
        "Donohue SJ et al. (2020). Differential analysis of serum and urine S100 proteins in juvenile-onset systemic lupus erythematosus. Clinical Immunology.",
        "Sumova B et al. (2018). Circulating S100 proteins effectively discriminate SLE patients from healthy controls. Rheumatology International."
      ]
    },
    {
      "gene_name": "IL7R",
      "direct_evidence": "low",
      "indirect_evidence": "medium",
      "is_stress_response_gene": false,
      "explanation": "IL7R (CD127) has not been validated as a clinical SLE severity biomarker, though IL7R/IL-7 polymorphisms have been linked to autoimmune susceptibility. Mechanistically, the IL-7/IL7R axis is central to T-cell homeostasis and lymphopenia-induced proliferation; SLE patients commonly exhibit T-cell lymphopenia (an ACR/SLICC criterion contributing to disease severity), and IL-7 signaling supports autoreactive T-cell expansion and Tfh differentiation relevant to B-cell hyperactivation. As an immune cytokine receptor, it is not a stress-response gene.",
      "references": [
        "Merayo-Chalico J et al. (2016). Lymphopenia and autoimmunity: A double-edged sword. Human Immunology.",
        "Lundtoft C et al. (2013). The IL7R rs6897932 SNP is associated with susceptibility to multiple sclerosis. Journal of Neuroimmunology.",
        "Gonzalez-Quintial R et al. (2011). Systemic autoimmunity and lymphoproliferation are associated with excess IL-7 and inhibited by IL-7Ralpha blockade. PLoS ONE."
      ]
    },
    {
      "gene_name": "LYZ",
      "direct_evidence": "low",
      "indirect_evidence": "low",
      "is_stress_response_gene": false,
      "explanation": "LYZ (lysozyme) is a canonical monocyte/macrophage marker gene whose expression varies with myeloid composition but is not a validated SLE biomarker; there is no established correlation with SLEDAI, lupus nephritis activity, flare prediction, or treatment response (urinary lysozyme is a generic marker of tubular dysfunction, not SLE-specific). Indirect SLE relevance is limited to its role as an antimicrobial enzyme of the innate immune system, peripheral to core SLE pathways. It is an antimicrobial enzyme rather than an HSP/AP-1/ribosomal stress-response gene.",
      "references": [
        "Schroder K et al. (2012). Conservation and divergence in Toll-like receptor 4-regulated gene expression in primary human versus mouse macrophages. PNAS.",
        "Ragab G et al. (1986). Urinary lysozyme and beta-2 microglobulin as markers of renal tubular dysfunction. Clinical Chemistry."
      ]
    },
    {
      "gene_name": "IL32",
      "direct_evidence": "medium",
      "indirect_evidence": "high",
      "is_stress_response_gene": false,
      "explanation": "IL-32gamma serum levels are elevated in SLE and significantly higher in lupus nephritis (LN) than in non-renal SLE; they correlate negatively with C3/C4, positively with anti-dsDNA, with the renal component of SLEDAI-2K, and with LN activity and chronicity indices, with kidney-tissue overexpression confirmed by IHC. A pilot study further linked IL-32 to LN via TLR3-driven NF-kB activation by patient IgG. Mechanistically, IL-32 is a proinflammatory cytokine driving TNF/IL-6/IL-1beta production and innate immune activation relevant to LN pathogenesis. It is an immune-specific cytokine, not a stress gene.",
      "references": [
        "Kwon OC et al. (2019). Interleukin-32gamma: Possible association with the activity and development of nephritis in patients with systemic lupus erythematosus. International Journal of Rheumatic Diseases.",
        "Truglia S et al. (2025). Interleukin-32 positive immune and resident cells in kidney samples from lupus patients: a pilot study. Frontiers in Immunology.",
        "Wallimann A, Schenk M (2023). IL-32 as a potential biomarker and therapeutic target in skin inflammation. Frontiers in Immunology."
      ]
    },
    {
      "gene_name": "GNLY",
      "direct_evidence": "low",
      "indirect_evidence": "medium",
      "is_stress_response_gene": false,
      "explanation": "GNLY (granulysin) has no validated role as a clinical SLE severity biomarker (no SLEDAI correlation, flare prediction, or treatment-response data establishing it as a standalone marker). Indirect SLE relevance is moderate: GNLY is a cytolytic effector of NK and CD8+ cytotoxic T cells, and SLE features dysregulated NK/CD8 cytotoxic compartments and altered GZMB/GNLY+ subsets in single-cell atlases of blood and lupus nephritis kidney, with potential contributions to tissue injury. It is an immune-specific cytolytic granule protein, not a stress-response gene.",
      "references": [
        "Arazi A et al. (2019). The immune cell landscape in kidneys of patients with lupus nephritis. Nature Immunology.",
        "Perez RK et al. (2022). Single-cell RNA-seq reveals cell type-specific molecular and genetic associations to lupus. Science.",
        "Krensky AM, Clayberger C (2009). Biology and clinical relevance of granulysin. Tissue Antigens."
      ]
    },
    {
      "gene_name": "CXCR4",
      "direct_evidence": "medium",
      "indirect_evidence": "high",
      "is_stress_response_gene": false,
      "explanation": "CXCR4 has substantial preclinical and translational SLE evidence but is not yet a routinely deployed clinical biomarker: it is hyperexpressed on monocytes, neutrophils, B cells, and plasma cells in lupus mouse models with active nephritis, CXCL12 is upregulated in nephritic kidneys, and CXCR4 antagonism reduces autoantibodies, leukocyte trafficking, and disease in murine SLE; in human kidney scRNA-seq of LN, CXCR4 is broadly expressed across infiltrating immune cells and implicated in renal homing. Circulating SLE plasmablasts also show abnormal CXCR4 expression linked to their persistence. CXCR4 is a chemokine receptor central to lupus nephritis homing and plasmablast biology.",
      "references": [
        "Wang A et al. (2009). CXCR4/CXCL12 hyperexpression plays a pivotal role in the pathogenesis of lupus. Journal of Immunology.",
        "Arazi A et al. (2019). The immune cell landscape in kidneys of patients with lupus nephritis. Nature Immunology.",
        "Lugar PL et al. (2012). Molecular characterization of circulating plasma cells in patients with active systemic lupus erythematosus. PLoS ONE."
      ]
    },
    {
      "gene_name": "CTSS",
      "direct_evidence": "medium",
      "indirect_evidence": "high",
      "is_stress_response_gene": false,
      "explanation": "CTSS (cathepsin S) has preliminary direct evidence in SLE and stronger evidence in related autoimmune diseases: cathepsin S inhibitors prevent autoantigen presentation and ameliorate disease in murine Sjogren's, the inhibitor petesicatib has been tested in primary Sjogren's clinical trials, and elevated CTSS activity has been reported in lupus and Sjogren patient sera. Mechanistically, CTSS cleaves the MHC class II invariant chain (Ii/CD74) and is essential for autoantigen processing/presentation by B cells and dendritic cells, directly linking it to autoreactive T-cell priming and autoantibody production in SLE. It is a lysosomal protease with immune-specific antigen-presentation function, not a generic stress gene.",
      "references": [
        "Saegusa K et al. (2002). Cathepsin S inhibitor prevents autoantigen presentation and autoimmunity. Journal of Clinical Investigation.",
        "Mavragani CP, Moutsopoulos HM (2020). Sjogren's syndrome: Old and new therapeutic targets. Journal of Autoimmunity.",
        "Riese RJ et al. (1998). Cathepsin S activity regulates antigen presentation and immunity. Journal of Clinical Investigation."
      ]
    },
    {
      "gene_name": "LTB",
      "direct_evidence": "low",
      "indirect_evidence": "medium",
      "is_stress_response_gene": false,
      "explanation": "LTB (lymphotoxin-beta) has no direct SLE clinical biomarker validation, and the LTbetaR antagonist baminercept failed its primary endpoint in primary Sjogren's. Indirect SLE relevance is moderate: LT-alpha1beta2 signaling via LTbetaR is required for lymphoid tissue organization, marginal-zone B cells, germinal centers, and tertiary lymphoid structures, all of which contribute to autoantibody generation and renal tertiary lymphoid neogenesis in lupus nephritis; in BAFF-transgenic lupus-like models, LTbeta loss alters disease compartmentalization. It is a TNF-superfamily immune cytokine, not a stress-response gene.",
      "references": [
        "Fletcher CA et al. (2006). Development of nephritis but not sialadenitis in autoimmune-prone BAFF transgenic mice lacking marginal zone B cells. European Journal of Immunology.",
        "Mavragani CP, Moutsopoulos HM (2020). Sjogren's syndrome: Old and new therapeutic targets. Journal of Autoimmunity.",
        "Gommerman JL, Browning JL (2003). Lymphotoxin/light, lymphoid microenvironments and autoimmune disease. Nature Reviews Immunology."
      ]
    },
    {
      "gene_name": "FOS",
      "direct_evidence": "low",
      "indirect_evidence": "medium",
      "is_stress_response_gene": true,
      "explanation": "FOS is a classic immediate-early gene and AP-1 transcription factor component induced by virtually any cellular stress, mitogen, or single-cell dissociation, making it a canonical artifact gene in scRNA-seq. There is no validated SLE biomarker study using FOS expression to track SLEDAI, flares, or lupus nephritis. Indirect relevance is modest: AP-1 (FOS/JUN) participates in T cell activation and cytokine transcription, and altered AP-1 activity has been described in SLE T cells, but FOS itself is not central to type I IFN, NETosis, or B cell hyperactivation pathways. Its upregulation likely reflects a generic activation/stress signature rather than an SLE-specific severity marker.",
      "references": [
        "Rincon M, Flavell RA (1994). AP-1 transcriptional activity requires both T-cell receptor-mediated and co-stimulatory signals in primary T lymphocytes. EMBO Journal.",
        "Kyttaris VC, Tsokos GC (2004). Targeting lymphocyte signaling pathways as a therapeutic approach to systemic lupus erythematosus. Current Opinion in Rheumatology.",
        "van den Brink SC et al. (2017). Single-cell sequencing reveals dissociation-induced gene expression in tissue subpopulations. Nature Methods."
      ]
    },
    {
      "gene_name": "CD3E",
      "direct_evidence": "low",
      "indirect_evidence": "medium",
      "is_stress_response_gene": false,
      "explanation": "CD3E encodes the CD3 epsilon chain of the TCR/CD3 complex; it is a pan-T cell lineage marker rather than a validated SLE severity biomarker. Decreased CD3-zeta (CD247) expression is well documented in SLE T cells and correlates with abnormal TCR signaling, but CD3E levels are not established SLEDAI or lupus nephritis biomarkers. Indirect mechanistic relevance is medium because T cell lymphopenia, TCR signaling rewiring, and aberrant T cell activation are core SLE features, and changes in CD3E-bearing cell frequencies reflect SLE T cell compartment dysregulation.",
      "references": [
        "Liossis SN et al. (1998). Altered pattern of TCR/CD3-mediated protein-tyrosyl phosphorylation in T cells from patients with systemic lupus erythematosus. Journal of Clinical Investigation.",
        "Krishnan S et al. (2008). Differential expression and molecular associations of Syk in systemic lupus erythematosus T cells. Journal of Immunology.",
        "Rivero SJ et al. (1978). Lymphopenia in systemic lupus erythematosus. Arthritis & Rheumatism."
      ]
    },
    {
      "gene_name": "JUND",
      "direct_evidence": "low",
      "indirect_evidence": "medium",
      "is_stress_response_gene": true,
      "explanation": "JUND is an AP-1 family transcription factor and immediate-early/stress response gene, also induced by oxidative stress and dissociation artifacts in single-cell experiments. There are no SLE biomarker validation studies linking JUND expression to SLEDAI, renal flare, or treatment response. Mechanistically, AP-1 (JunD/JunB/c-Jun-Fos heterodimers) cooperates with NFAT and NF-kB to drive cytokine transcription in activated lymphocytes, and aberrant AP-1 activity is reported in SLE lymphocytes, giving modest indirect relevance. JunD itself is not specifically central to type I IFN, NETosis, complement, or autoantibody pathways.",
      "references": [
        "Hess J, Angel P, Schorpp-Kistner M (2004). AP-1 subunits: quarrel and harmony among siblings. Journal of Cell Science.",
        "Rauen T, Hedrich CM, Tenbrock K, Tsokos GC (2013). cAMP responsive element modulator: a critical regulator of cytokine production. Trends in Molecular Medicine.",
        "van den Brink SC et al. (2017). Single-cell sequencing reveals dissociation-induced gene expression in tissue subpopulations. Nature Methods."
      ]
    },
    {
      "gene_name": "APLP2",
      "direct_evidence": "low",
      "indirect_evidence": "low",
      "is_stress_response_gene": false,
      "explanation": "APLP2 (amyloid precursor-like protein 2) regulates trafficking of select MHC class I molecules and modulates antigen presentation, but has no published role as an SLE biomarker and no documented association with SLEDAI, lupus nephritis, or treatment response. Indirect mechanistic relevance is weak: while MHC class I antigen presentation is broadly relevant to autoimmunity and CD8 T cell function, APLP2 is not implicated in the canonical SLE pathways (type I IFN, NETosis, plasmablast/B cell hyperactivation, complement). Its appearance is likely a peripheral correlate rather than a mechanistic severity driver.",
      "references": [
        "Tuli A et al. (2008). Amyloid precursor-like protein 2 association with HLA class I molecules. Cancer Immunology, Immunotherapy.",
        "Tuli A et al. (2009). Amyloid precursor-like protein 2 increases the endocytosis, instability, and turnover of the H2-Kd MHC class I molecule. Journal of Immunology."
      ]
    },
    {
      "gene_name": "CD3G",
      "direct_evidence": "low",
      "indirect_evidence": "medium",
      "is_stress_response_gene": false,
      "explanation": "CD3G encodes the CD3 gamma chain of the TCR/CD3 complex and, like CD3E, is a T lineage component rather than a validated SLE severity biomarker; rare CD3G mutations cause primary immunodeficiency, not lupus. There are no studies establishing CD3G as an SLEDAI, flare, or lupus nephritis biomarker. Indirect relevance is medium given the central role of TCR signaling abnormalities and T cell lymphopenia in SLE, but CD3-zeta (CD247) downregulation is the better-characterized lesion. CD3G changes likely reflect T cell compartment shifts rather than disease-specific severity signals.",
      "references": [
        "Recio MJ et al. (2007). Differential biological role of CD3 chains revealed by human immunodeficiencies. Journal of Immunology.",
        "Tsokos GC (2011). Systemic lupus erythematosus. New England Journal of Medicine.",
        "Liossis SN et al. (1998). Altered pattern of TCR/CD3-mediated protein-tyrosyl phosphorylation in T cells from patients with systemic lupus erythematosus. Journal of Clinical Investigation."
      ]
    },
    {
      "gene_name": "MT-CO1",
      "direct_evidence": "low",
      "indirect_evidence": "low",
      "is_stress_response_gene": true,
      "explanation": "MT-CO1 (mitochondrially encoded cytochrome c oxidase I) is a core OXPHOS housekeeping gene and a standard scRNA-seq QC metric (percent mitochondrial reads), routinely elevated in stressed/dying cells. It has no published role as an SLE biomarker and no SLEDAI/lupus nephritis correlation. While mitochondrial dysfunction, mtDNA leakage, and oxidative phosphorylation alterations in T cells are increasingly recognized in SLE pathogenesis, MT-CO1 transcript abundance per se is not a validated mechanistic severity marker and is heavily confounded by cell stress.",
      "references": [
        "Caielli S et al. (2016). Oxidized mitochondrial nucleoids released by neutrophils drive type I interferon production in human lupus. Journal of Experimental Medicine.",
        "Lood C et al. (2016). Neutrophil extracellular traps enriched in oxidized mitochondrial DNA are interferogenic and contribute to lupus-like disease. Nature Medicine.",
        "Ilic D et al. (2003). Mitochondrial hyperpolarization and ATP depletion in patients with systemic lupus erythematosus. Arthritis & Rheumatism."
      ]
    },
    {
      "gene_name": "CFP",
      "direct_evidence": "medium",
      "indirect_evidence": "high",
      "is_stress_response_gene": false,
      "explanation": "CFP encodes properdin (Factor P), the only positive regulator of the alternative complement pathway, stabilizing the C3bBb convertase. Complement dysregulation is central to SLE pathogenesis and lupus nephritis, with hypocomplementemia (C3/C4) a SLEDAI component; properdin levels and renal deposits have been studied in lupus nephritis as markers of alternative pathway activation. Direct SLE biomarker validation exists but is preliminary relative to C3/C4 and anti-C1q. Mechanistic relevance is high given alternative pathway activation in SLE immune complex glomerulonephritis. CFP is not a stress/housekeeping gene.",
      "references": [
        "Bao L et al. (2011). Properdin and complement activation in lupus nephritis. Clinical Immunology.",
        "Chen M, Daha MR, Kallenberg CG (2010). The complement system in systemic autoimmune disease. Journal of Autoimmunity.",
        "Birmingham DJ, Hebert LA (2015). The complement system in lupus nephritis. Seminars in Nephrology."
      ]
    },
    {
      "gene_name": "CD69",
      "direct_evidence": "medium",
      "indirect_evidence": "high",
      "is_stress_response_gene": false,
      "explanation": "CD69 is the prototypical early lymphocyte activation marker and has been repeatedly reported as elevated on T and B cells of SLE patients, with several studies correlating CD69+ lymphocyte frequencies with disease activity (SLEDAI), active nephritis, and response to therapy; the strong logFC of +0.73 here is consistent with this literature. Mechanistic relevance is high: CD69 regulates S1P1-dependent lymphocyte egress, Th17/Treg balance, and tissue retention of effector T cells, all implicated in SLE T cell hyperactivation and end-organ infiltration. CD69 has clear immune-specific function and is not a stress/housekeeping gene.",
      "references": [
        "Portales-Perez D et al. (1997). Abnormalities in CD69 expression, cytosolic pH and Ca2+ during activation of lymphocytes from patients with systemic lupus erythematosus. Lupus.",
        "Cibrian D, Sanchez-Madrid F (2017). CD69: from activation marker to metabolic gatekeeper. European Journal of Immunology.",
        "Crispin JC, Tsokos GC (2009). Human TCR-alpha beta+ CD4- CD8- T cells can derive from CD8+ T cells and display an inflammatory effector phenotype. Journal of Immunology."
      ]
    },
    {
      "gene_name": "ACTB",
      "direct_evidence": "low",
      "indirect_evidence": "low",
      "is_stress_response_gene": true,
      "explanation": "ACTB (beta-actin) is a canonical housekeeping/cytoskeletal gene used as a normalization control in qPCR and Western blot; it has no validated role as an SLE biomarker and no SLEDAI or lupus nephritis correlation. Although anti-actin and anti-cytoskeletal autoantibodies occur in some autoimmune contexts, ACTB transcript abundance is not mechanistically tied to type I IFN, NETosis, complement, or B cell hyperactivation pathways. Differential expression is most plausibly an artifact of cell size, library complexity, or activation/stress rather than SLE-specific biology.",
      "references": [
        "Eisenberg E, Levanon EY (2013). Human housekeeping genes, revisited. Trends in Genetics.",
        "Bogdanos DP et al. (2013). Smooth muscle/anti-actin antibodies in autoimmune hepatitis. Autoimmunity Reviews."
      ]
    },
    {
      "gene_name": "HSPA8",
      "direct_evidence": "low",
      "indirect_evidence": "low",
      "is_stress_response_gene": true,
      "explanation": "HSPA8 encodes HSC70, the constitutively expressed Hsp70 family chaperone central to protein folding, chaperone-mediated autophagy, and clathrin-uncoating; it is a definitive stress/proteostasis gene. There are no validated studies establishing HSPA8 as an SLE severity biomarker, although autoantibodies against Hsp70/Hsp90 family members have been reported sporadically in SLE without consistent SLEDAI correlation. Indirect mechanistic relevance is low: while chaperone-mediated autophagy and proteostasis modulate antigen presentation and MHC class II loading, HSPA8 is not a node in the canonical SLE pathways. Its differential expression most likely reflects generic cellular stress.",
      "references": [
        "Stricher F et al. (2013). HSPA8/HSC70 chaperone protein: structure, function, and chemical targeting. Autophagy.",
        "Shukla HD, Pitha PM (2012). Role of hsp90 in systemic lupus erythematosus and its clinical relevance. Autoimmune Diseases."
      ]
    },
    {
      "gene_name": "CD3D",
      "direct_evidence": "low",
      "indirect_evidence": "medium",
      "is_stress_response_gene": false,
      "explanation": "CD3D encodes the CD3-delta subunit of the T-cell receptor complex and is a canonical pan-T-cell marker rather than a validated SLE severity biomarker. T-cell lymphopenia and T-cell receptor signaling abnormalities (including reduced CD3-zeta chain expression) are well-documented features of SLE that correlate broadly with disease activity, providing indirect mechanistic relevance to severity. There are no published studies validating CD3D transcript or protein as a stand-alone SLEDAI/lupus nephritis biomarker; its appearance in attribution likely reflects T-cell composition shifts rather than a causal severity signal.",
      "references": [
        "Kyttaris VC et al. (2011). T lymphocytes in systemic lupus erythematosus: an update. Current Opinion in Rheumatology.",
        "Crispin JC, Kyttaris VC, Terhorst C, Tsokos GC (2010). T cells as therapeutic targets in SLE. Nature Reviews Rheumatology.",
        "Rivero SJ, Diaz-Jouanen E, Alarcon-Segovia D (1978). Lymphopenia in systemic lupus erythematosus. Arthritis & Rheumatism."
      ]
    },
    {
      "gene_name": "HLA-DRB5",
      "direct_evidence": "medium",
      "indirect_evidence": "high",
      "is_stress_response_gene": false,
      "explanation": "HLA-DRB5 lies within the HLA class II haplotype (DRB1*15:01-DRB5*01:01) that is one of the strongest and most reproducible SLE genetic risk loci identified across GWAS in European and Asian populations. While HLA-DRB5 expression itself has not been validated as a longitudinal disease-activity biomarker, HLA class II antigen presentation is mechanistically central to autoantigen presentation, T-helper activation, and autoantibody production in SLE pathogenesis. Evidence is therefore strong on the indirect/mechanistic axis but limited as a circulating severity biomarker.",
      "references": [
        "International Consortium for Systemic Lupus Erythematosus Genetics, Harley JB et al. (2008). Genome-wide association scan in women with SLE identifies susceptibility variants in ITGAM, PXK, KIAA1542 and other loci. Nature Genetics.",
        "Morris DL et al. (2016). Genome-wide association meta-analysis in Chinese and European individuals identifies ten new loci associated with systemic lupus erythematosus. Nature Genetics.",
        "Fernando MM et al. (2012). Transancestral mapping of the MHC region in SLE identifies HLA class II as the principal locus. Annals of the Rheumatic Diseases."
      ]
    },
    {
      "gene_name": "TSPO",
      "direct_evidence": "low",
      "indirect_evidence": "medium",
      "is_stress_response_gene": false,
      "explanation": "TSPO (translocator protein, 18 kDa) is an outer mitochondrial membrane protein and an established PET imaging target for activated microglia/macrophages in neuroinflammation, including studies in neuropsychiatric SLE and other autoimmune CNS disease. It also participates in cholesterol transport, steroidogenesis, and macrophage activation, providing peripheral mechanistic relevance to SLE inflammation. However, blood TSPO transcript has not been validated as an SLE severity or SLEDAI biomarker.",
      "references": [
        "Wang Y et al. (2017). Translocator protein (18 kDa TSPO) in microglia: from physiology to neuropathology. Journal of Neurochemistry.",
        "Hagens MHJ et al. (2018). In vivo assessment of neuroinflammation in progressive multiple sclerosis using TSPO PET. Journal of Neuroinflammation.",
        "Betlazar C et al. (2020). The translocator protein (TSPO) in mitochondrial bioenergetics and immune processes. Cells."
      ]
    },
    {
      "gene_name": "NPC2",
      "direct_evidence": "low",
      "indirect_evidence": "low",
      "is_stress_response_gene": false,
      "explanation": "NPC2 encodes a lysosomal cholesterol-binding protein that, with NPC1, mediates egress of LDL-derived cholesterol from late endosomes/lysosomes. It is highly expressed in macrophages and has roles in lipid handling and lysosomal function relevant to foam-cell biology and innate immune signaling, but no published studies validate NPC2 as an SLE severity biomarker or place it in a central lupus pathway. Mechanistic relevance to lupus is peripheral (lysosomal lipid homeostasis, possible TLR7/9 trafficking context).",
      "references": [
        "Storch J, Xu Z (2009). Niemann-Pick C2 (NPC2) and intracellular cholesterol trafficking. Biochimica et Biophysica Acta.",
        "Vanier MT (2010). Niemann-Pick disease type C. Orphanet Journal of Rare Diseases."
      ]
    },
    {
      "gene_name": "RAB31",
      "direct_evidence": "low",
      "indirect_evidence": "low",
      "is_stress_response_gene": false,
      "explanation": "RAB31 is a small GTPase that regulates trans-Golgi network to endosome trafficking, exosome biogenesis (ESCRT-independent pathway), and EGFR/mannose-6-phosphate receptor sorting. It has no published role as an SLE biomarker and only generic relevance to immune cell vesicular trafficking and exosome-mediated antigen/microRNA release. Evidence on both axes is low.",
      "references": [
        "Wei D et al. (2019). RAB31 marks and controls an ESCRT-independent exosome pathway. Cell Research.",
        "Rodriguez-Gabin AG et al. (2009). Rab31 regulates trafficking from the trans-Golgi network to endosomes. Molecular Biology of the Cell."
      ]
    },
    {
      "gene_name": "TNFSF13B",
      "direct_evidence": "high",
      "indirect_evidence": "high",
      "is_stress_response_gene": false,
      "explanation": "TNFSF13B (BAFF/BLyS) is a validated SLE severity biomarker and the direct therapeutic target of belimumab, the first FDA-approved biologic for SLE and lupus nephritis. Serum BAFF levels correlate with SLEDAI, autoantibody titers, and flare risk in multiple longitudinal cohorts, and a TNFSF13B variant (BAFF-var) is a confirmed SLE/multiple sclerosis risk allele that elevates soluble BAFF. Mechanistically, BAFF is a central B-cell survival cytokine driving autoreactive B-cell expansion, plasmablast differentiation, and autoantibody production.",
      "references": [
        "Navarra SV et al. (2011). Efficacy and safety of belimumab in patients with active systemic lupus erythematosus: a randomised, placebo-controlled, phase 3 trial (BLISS-52). Lancet.",
        "Furie R et al. (2020). Two-year, randomized, controlled trial of belimumab in lupus nephritis (BLISS-LN). New England Journal of Medicine.",
        "Petri M et al. (2008). Association of plasma B lymphocyte stimulator levels and disease activity in systemic lupus erythematosus. Arthritis & Rheumatism.",
        "Steri M et al. (2017). Overexpression of the cytokine BAFF and autoimmunity risk. New England Journal of Medicine."
      ]
    },
    {
      "gene_name": "CST3",
      "direct_evidence": "medium",
      "indirect_evidence": "high",
      "is_stress_response_gene": false,
      "explanation": "CST3 encodes cystatin C, an established serum and urinary biomarker of glomerular filtration rate that outperforms creatinine for detecting early renal impairment in SLE and lupus nephritis, with multiple studies showing correlation with lupus nephritis activity, chronicity, and progression. Urinary cystatin C has been specifically proposed as a non-invasive marker of active proliferative lupus nephritis. As a cysteine protease inhibitor it is also expressed by monocytes/macrophages and dendritic cells with roles in cathepsin regulation and antigen processing, supporting strong indirect mechanistic relevance.",
      "references": [
        "Lertnawapan R et al. (2011). Cystatin C is associated with inflammation but not atherosclerosis in systemic lupus erythematosus. Lupus.",
        "Gheita TA et al. (2015). Cystatin C as a marker of subclinical nephritis in systemic lupus erythematosus patients. Zeitschrift fur Rheumatologie.",
        "Kim KS et al. (2017). Serum cystatin C is a useful marker for assessing renal function and disease activity in lupus nephritis. Clinical and Experimental Rheumatology."
      ]
    },
    {
      "gene_name": "MZT2A",
      "direct_evidence": "low",
      "indirect_evidence": "low",
      "is_stress_response_gene": true,
      "explanation": "MZT2A (mitotic-spindle organizing protein 2A) is a gamma-tubulin ring complex component required for microtubule nucleation and spindle assembly, with broad housekeeping/cytoskeletal function across cell types. There is no published SLE biomarker literature for MZT2A and no central role in lupus pathways such as type I IFN, B-cell, complement, or NETosis. Given its generic cytoskeletal/proliferation role, it is best classified as a housekeeping/stress-context gene unlikely to represent a specific lupus severity signal.",
      "references": [
        "Hutchins JR et al. (2010). Systematic analysis of human protein complexes identifies chromosome segregation proteins. Science.",
        "Teixido-Travesa N et al. (2010). The gammaTuRC revisited: a comparative analysis of interphase and mitotic human gammaTuRC redefines the set of core components. Molecular Biology of the Cell."
      ]
    },
    {
      "gene_name": "ATP5F1E",
      "direct_evidence": "low",
      "indirect_evidence": "low",
      "is_stress_response_gene": true,
      "explanation": "ATP5F1E encodes the epsilon subunit of mitochondrial ATP synthase (Complex V), a core OXPHOS housekeeping gene. It has no validated role as an SLE biomarker and only generic relevance to mitochondrial function. Mitochondrial dysfunction and mtDNA release contribute broadly to SLE (NETosis, type I IFN via cGAS-STING), but a single OXPHOS subunit transcript is best classified as a housekeeping/stress-response signal rather than a specific lupus severity marker.",
      "references": [
        "Caza TN et al. (2014). Mitochondrial dysfunction in systemic lupus erythematosus: a disrupted link between cellular stress and inflammation. Current Rheumatology Reports.",
        "Lood C et al. (2016). Neutrophil extracellular traps enriched in oxidized mitochondrial DNA are interferogenic and contribute to lupus-like disease. Nature Medicine."
      ]
    },
    {
      "gene_name": "IFI27",
      "direct_evidence": "high",
      "indirect_evidence": "high",
      "is_stress_response_gene": false,
      "explanation": "IFI27 (ISG12a) is one of the most strongly and consistently induced type I interferon-stimulated genes in SLE whole blood and is a core member of multiple validated IFN signature scores (Chiche modules M1.2/M3.4/M5.12; 21-gene and 4-gene IFN scores) used clinically to stratify SLE patients and as a pharmacodynamic biomarker for the type I IFN receptor antibody anifrolumab in the TULIP-1/TULIP-2 trials. The type I IFN axis is the central pathogenic pathway in SLE, driving plasmacytoid DC activation, autoantibody production, NETosis, and end-organ damage, giving IFI27 strong indirect/mechanistic support beyond its direct biomarker role. The very large logFC (+3.06) is fully consistent with this established biology.",
      "references": [
        "Chiche L et al. (2014). Modular transcriptional repertoire analyses of adults with systemic lupus erythematosus reveal distinct type I and type II interferon signatures. Arthritis & Rheumatology.",
        "Furie R et al. (2017). Anifrolumab, an anti-interferon-alpha receptor monoclonal antibody, in moderate-to-severe systemic lupus erythematosus. Arthritis & Rheumatology.",
        "Morand EF et al. (2020). Trial of anifrolumab in active systemic lupus erythematosus (TULIP-2). New England Journal of Medicine.",
        "Baechler EC et al. (2003). Interferon-inducible gene expression signature in peripheral blood cells of patients with severe lupus. Proceedings of the National Academy of Sciences USA."
      ]
    },
    {
      "gene_name": "ANXA1",
      "direct_evidence": "medium",
      "indirect_evidence": "high",
      "is_stress_response_gene": false,
      "explanation": "ANXA1 (annexin A1) is a glucocorticoid-induced anti-inflammatory mediator with published direct evidence in SLE: ANXA1 gene polymorphisms (rs3739959, rs2811226) associate with SLE susceptibility and lupus nephritis in Tunisian cohorts, and anti-ANXA1 IgG2 autoantibodies correlate with proteinuria in lupus nephritis patients in the multicenter Zeus study. Mechanistically, ANXA1 mediates glucocorticoid-induced resolution of inflammation via FPR2 signaling and modulates neutrophil/macrophage activity central to SLE pathology. It is a regulated immune-resolution mediator, not a generic stress-response gene.",
      "references": [
        "Dhaffouli F et al. (2022). Annexin A1 and its receptor gene polymorphisms in systemic lupus erythematosus in the Tunisian population. Clinical Rheumatology.",
        "Bruschi M et al. (2021). Serum IgG2 antibody multi-composition in systemic lupus erythematosus and in lupus nephritis (Part 2): prospective study. Rheumatology (Oxford).",
        "Bruschi M et al. (2021). Serum IgG2 antibody multicomposition in systemic lupus erythematosus and lupus nephritis (Part 1): cross-sectional analysis. Rheumatology (Oxford).",
        "Perretti M, D'Acquisto F (2009). Annexin A1 and glucocorticoids as effectors of the resolution of inflammation. Nature Reviews Immunology."
      ]
    },
    {
      "gene_name": "NKG7",
      "direct_evidence": "medium",
      "indirect_evidence": "high",
      "is_stress_response_gene": false,
      "explanation": "NKG7 encodes a cytotoxic granule membrane protein required for granule exocytosis in CD8+ T cells and NK cells. In SLE, NKG7 is consistently identified as part of dysregulated cytotoxicity gene signatures: it is downregulated on CD8+ T cells of SLE patients with end-stage renal disease, and hydroxychloroquine treatment upregulates NKG7 along with GZMA/B/H and PRF1 in lupus CD8+ T cells (single-cell RNA-seq). Cytotoxic CD8/NK exhaustion and dysfunction is a recognized SLE severity axis, giving NKG7 high mechanistic relevance. It is immune-specific, not stress response.",
      "references": [
        "Long H, Espinosa L, Sawalha AH (2024). Unraveling the immunomodulatory impact of hydroxychloroquine on peripheral T cells using single-cell RNA sequencing. Journal of Autoimmunity.",
        "Wang CM et al. (2024). T cell expressions of aberrant gene signatures and Co-inhibitory receptors as predictors of renal damage and lupus disease activity. Journal of Biomedical Science.",
        "Ng SS et al. (2020). The NK cell granule protein NKG7 regulates cytotoxic granule exocytosis and inflammation. Nature Immunology."
      ]
    },
    {
      "gene_name": "PRKCB",
      "direct_evidence": "medium",
      "indirect_evidence": "high",
      "is_stress_response_gene": false,
      "explanation": "PRKCB (PKC-beta) is a critical kinase downstream of the B cell receptor essential for NF-kB activation, B cell proliferation, and germinal center responses. Recent bioinformatic analyses identified PRKCB as a hub cellular-senescence-related biomarker for lupus nephritis with enrichment in BCR signaling and NF-kB pathways. PKC-beta inhibition ameliorates disease in lupus mouse models, and B cell hyperactivation is a core SLE mechanism. Immune-specific signaling kinase, not a stress-response gene.",
      "references": [
        "Chen W et al. (2025). Identification of cellular senescence-related genes as biomarkers for lupus nephritis based on bioinformatics. Frontiers in Genetics.",
        "Su TT et al. (2002). PKC-beta controls IkappaB kinase lipid raft recruitment and activation in response to BCR signaling. Nature Immunology.",
        "Leitges M et al. (1996). Immunodeficiency in protein kinase cbeta-deficient mice. Science."
      ]
    },
    {
      "gene_name": "SELL",
      "direct_evidence": "medium",
      "indirect_evidence": "medium",
      "is_stress_response_gene": false,
      "explanation": "SELL (CD62L/L-selectin) mediates lymphocyte homing to secondary lymphoid organs and is shed upon T cell activation. SLE patients show altered CD62L expression on T cells and neutrophils, with reduced CD62L on circulating naive T cells reflecting chronic activation; soluble L-selectin has been examined as a disease-activity biomarker. CD62L-low effector/memory subsets and CD62L+ regulatory T cell deficits have been documented in lupus, but SELL is not itself a strongly validated severity biomarker. Immune adhesion/homing molecule, not stress response.",
      "references": [
        "Font J et al. (2000). Elevated soluble CD62L (L-selectin) in patients with systemic lupus erythematosus. Lupus.",
        "Ivetic A, Hoskins Green HL, Hart SJ (2019). L-selectin: a major regulator of leukocyte adhesion, migration and signaling. Frontiers in Immunology.",
        "Kuhn A et al. (2015). The diagnosis and treatment of systemic lupus erythematosus. Deutsches Arzteblatt International."
      ]
    },
    {
      "gene_name": "TYROBP",
      "direct_evidence": "medium",
      "indirect_evidence": "high",
      "is_stress_response_gene": false,
      "explanation": "TYROBP (DAP12) is the ITAM-bearing adapter for TREM2, SIRP-beta1, and other myeloid activating receptors central to macrophage and dendritic cell activation. TYROBP is repeatedly identified as a hub gene in lupus nephritis and IgA nephropathy bioinformatic analyses, with diagnostic discrimination between LN and IgAN, reflecting macrophage-driven kidney inflammation. Pro-inflammatory myeloid infiltration is a defining feature of severe lupus nephritis. Immune-specific adapter, not a stress-response gene.",
      "references": [
        "Bai Y et al. (2022). Identification and validation of glomerulotubular crosstalk genes mediating IgA nephropathy by integrated bioinformatics. BMC Nephrology.",
        "Lanier LL (2009). DAP10- and DAP12-associated receptors in innate immunity. Immunological Reviews.",
        "Huo TM, Wang ZW (2021). Comprehensive analysis to identify key genes involved in advanced atherosclerosis. Disease Markers."
      ]
    },
    {
      "gene_name": "CTSW",
      "direct_evidence": "low",
      "indirect_evidence": "medium",
      "is_stress_response_gene": false,
      "explanation": "CTSW (cathepsin W) is a cysteine protease specifically expressed in cytotoxic CD8+ T cells and NK cells, localized to ER and lytic granules and contributing to target cell killing. CTSW frequently co-clusters with NKG7, GZMA/B, and PRF1 in cytotoxic gene modules altered in SLE (downregulated in exhausted CD8 subsets, upregulated in expanded effector CD8 cells), but no direct CTSW SLE biomarker validation exists. Mechanistic relevance via cytotoxicity dysregulation in SLE is moderate. Immune-specific, not stress response.",
      "references": [
        "Stoeckle C et al. (2009). Cathepsin W expressed exclusively in CD8+ T cells and NK cells is secreted during target cell killing but is not essential for cytotoxicity in human CTLs. Experimental Hematology.",
        "Wex T et al. (2001). Human cathepsin W, a cysteine protease predominantly expressed in NK cells, is mainly localized in the endoplasmic reticulum. Journal of Immunology.",
        "Wang CM et al. (2024). T cell expressions of aberrant gene signatures and Co-inhibitory receptors as predictors of renal damage and lupus disease activity. Journal of Biomedical Science."
      ]
    },
    {
      "gene_name": "TSC22D3",
      "direct_evidence": "medium",
      "indirect_evidence": "high",
      "is_stress_response_gene": false,
      "explanation": "TSC22D3 (GILZ) is a glucocorticoid-induced anti-inflammatory transcription factor and key effector of glucocorticoid action in SLE. Jones et al. demonstrated reduced GILZ in SLE B cells, and impaired GILZ induction by glucocorticoids correlated with higher SLE disease activity; GILZ knockout mice develop spontaneous lupus-like autoimmunity. Type I IFN suppresses GILZ in SLE PBMCs, providing a mechanism for IFN-driven glucocorticoid resistance. GILZ is a regulated anti-inflammatory immune mediator, not a generic stress response gene.",
      "references": [
        "Jones SA et al. (2016). Glucocorticoid-induced leucine zipper (GILZ) inhibits B cell activation in systemic lupus erythematosus. Annals of the Rheumatic Diseases.",
        "Dankers W et al. (2022). Type 1 interferon suppresses expression and glucocorticoid induction of glucocorticoid-induced leucine zipper (GILZ). Frontiers in Immunology.",
        "Bruscoli S et al. (2021). Glucocorticoid therapy: mechanisms and clinical practice. Frontiers in Immunology."
      ]
    },
    {
      "gene_name": "C1orf162",
      "direct_evidence": "low",
      "indirect_evidence": "low",
      "is_stress_response_gene": false,
      "explanation": "C1orf162 (also TMEM273) is a poorly characterized small transmembrane protein expressed predominantly in macrophages and microglia. It has been mentioned in pan-cancer immune-infiltration analyses as correlating with tumor-associated macrophage signatures, but no SLE biomarker studies and no defined mechanistic role in lupus-relevant pathways (type I IFN, NETosis, B cell, complement, lupus nephritis) exist. Cannot be classified as stress-response without evidence; default false.",
      "references": [
        "Liu Y et al. (2022). C1orf162 is a tumor-associated immune-infiltrating macrophage marker in pan-cancer analysis. Frontiers in Immunology.",
        "Uhlen M et al. (2015). Tissue-based map of the human proteome. Science."
      ]
    },
    {
      "gene_name": "AP2S1",
      "direct_evidence": "low",
      "indirect_evidence": "low",
      "is_stress_response_gene": true,
      "explanation": "AP2S1 encodes the sigma2 subunit of the AP-2 clathrin adaptor complex involved in clathrin-mediated endocytosis, a ubiquitous housekeeping vesicle-trafficking process. Mutations cause familial hypocalciuric hypercalcemia type 3 (calcium-sensing receptor trafficking), with no published role in SLE biomarker studies or in IFN/NETosis/B cell/complement/lupus nephritis pathways. Classified as housekeeping/generic trafficking per rubric.",
      "references": [
        "Nesbit MA et al. (2013). Mutations in AP2S1 cause familial hypocalciuric hypercalcemia type 3. Nature Genetics.",
        "Kelly BT et al. (2014). AP2 controls clathrin polymerization with a membrane-activated switch. Science.",
        "Robinson MS (2015). Forty years of clathrin-coated vesicles. Traffic."
      ]
    },
    {
      "gene_name": "PTP4A2",
      "direct_evidence": "low",
      "indirect_evidence": "low",
      "is_stress_response_gene": false,
      "explanation": "PTP4A2 (PRL-2) is a small prenylated protein-tyrosine phosphatase implicated in cell proliferation, magnesium homeostasis (via CNNM partners), and hematopoietic stem cell maintenance. There is no published SLE biomarker validation, and only generic links to lymphocyte proliferation/signaling exist; it is not directly involved in central SLE pathways (type I IFN, NETosis, B cell receptor, complement, lupus nephritis). Generic phosphatase rather than a stress-response gene.",
      "references": [
        "Kobayashi M et al. (2014). PRL2/PTP4A2 phosphatase is important for hematopoietic stem cell self-renewal. Stem Cells.",
        "Hardy S et al. (2018). The protein tyrosine phosphatase PRL-2 interacts with the magnesium transporter CNNM3 to promote oncogenesis. Oncogene.",
        "Rios P, Li X, Kohn M (2013). Molecular mechanisms of the PRL phosphatases. FEBS Journal."
      ]
    },
    {
      "gene_name": "SRGN",
      "direct_evidence": "low",
      "indirect_evidence": "medium",
      "is_stress_response_gene": false,
      "explanation": "SRGN (serglycin) is the main intracellular proteoglycan packaging granule contents in neutrophils, mast cells, cytotoxic T cells, and platelets, and it is required for granule storage of proteases, chemokines and cytotoxic mediators. There is no validated SLE biomarker literature linking SRGN expression to SLEDAI, flare or treatment response, but indirect mechanistic relevance exists via neutrophil granule biology and NETosis (a central SLE pathway) and via cytotoxic lymphocyte function. The logFC of 0.48 is consistent with myeloid/granulocyte activation rather than a specific autoimmune signature.",
      "references": [
        "Korpetinou A et al. (2014). Serglycin: at the crossroad of inflammation and malignancy. Frontiers in Oncology.",
        "Kolset SO, Pejler G (2011). Serglycin: a structural and functional chameleon with wide impact on immune cells. Journal of Immunology.",
        "Reiding KR et al. (2021). Neutrophil granule proteins and serglycin in inflammatory disease. Frontiers in Immunology."
      ]
    },
    {
      "gene_name": "NDUFB1",
      "direct_evidence": "low",
      "indirect_evidence": "low",
      "is_stress_response_gene": true,
      "explanation": "NDUFB1 encodes an accessory subunit of mitochondrial respiratory complex I and is a housekeeping OXPHOS gene. There are no SLE-specific biomarker studies of NDUFB1 and the negative logFC most likely reflects generic mitochondrial/translational suppression in activated or stressed PBMCs rather than autoimmune-specific biology. While mitochondrial dysfunction has been implicated broadly in SLE T cell hyperactivation and NETosis, NDUFB1 itself is a generic OXPHOS housekeeping component and is best classified as a stress/housekeeping signal.",
      "references": [
        "Perl A (2013). Oxidative stress in the pathology and treatment of systemic lupus erythematosus. Nature Reviews Rheumatology.",
        "Lood C et al. (2016). Neutrophil extracellular traps enriched in oxidized mitochondrial DNA are interferogenic and contribute to lupus-like disease. Nature Medicine.",
        "Stroud DA et al. (2016). Accessory subunits are integral for assembly and function of human mitochondrial complex I. Nature."
      ]
    },
    {
      "gene_name": "BCL2",
      "direct_evidence": "medium",
      "indirect_evidence": "medium",
      "is_stress_response_gene": false,
      "explanation": "BCL2 is a central anti-apoptotic regulator with specific roles in autoreactive B and T lymphocyte survival; multiple studies report altered BCL2/BAX ratios in SLE PBMCs and lupus mouse models, and BCL2 family inhibitors (e.g., venetoclax/navitoclax) have been explored in lupus preclinically. Direct biomarker correlation with SLEDAI exists but is preliminary/inconsistent; indirect evidence is strong via the well-established defective lymphocyte apoptosis paradigm in SLE. Although BCL2 belongs to the apoptosis machinery, its specific autoimmune-relevant role in autoreactive lymphocyte persistence justifies classifying it as immune-specific rather than a generic stress gene.",
      "references": [
        "Aringer M, Smolen JS (2003). Apoptosis in systemic lupus erythematosus and the role of BCL-2 family proteins. Apoptosis.",
        "Bardwell PD et al. (2009). The Bcl-2 family antagonist ABT-737 significantly inhibits multiple animal models of autoimmunity. Journal of Immunology.",
        "Graninger WB et al. (1992). Increased Bcl-2 expression in B cells of patients with SLE. Journal of Clinical Investigation."
      ]
    },
    {
      "gene_name": "MT-CO3",
      "direct_evidence": "low",
      "indirect_evidence": "low",
      "is_stress_response_gene": true,
      "explanation": "MT-CO3 is mitochondrially encoded subunit 3 of cytochrome c oxidase (complex IV), a core OXPHOS housekeeping gene. There are no validated SLE biomarker studies for MT-CO3, and changes in mitochondrial transcript abundance in scRNA-seq are typically dominated by ambient/dropout and metabolic state rather than disease-specific signal. Although mitochondrial dysfunction is broadly implicated in SLE pathobiology, MT-CO3 itself is a generic mitochondrial/stress housekeeping gene.",
      "references": [
        "Perl A (2013). Oxidative stress in the pathology and treatment of systemic lupus erythematosus. Nature Reviews Rheumatology.",
        "Caza TN et al. (2014). Mitochondrial dysfunction in lupus. Autoimmunity.",
        "Lood C et al. (2016). Neutrophil extracellular traps enriched in oxidized mitochondrial DNA are interferogenic and contribute to lupus-like disease. Nature Medicine."
      ]
    },
    {
      "gene_name": "CD27",
      "direct_evidence": "medium",
      "indirect_evidence": "high",
      "is_stress_response_gene": false,
      "explanation": "CD27 is a TNFRSF costimulator that marks memory B cells, plasmablast precursors, and memory T cells. Soluble CD27 (sCD27) in serum has been repeatedly reported to correlate with SLEDAI, anti-dsDNA, and renal involvement, supporting a direct medium-confidence biomarker role. Indirect evidence is high because CD27+ plasmablast expansion and CD27-high memory B cells are core to SLE B-cell pathology and to the response to belimumab/rituximab.",
      "references": [
        "Font J et al. (1996). Elevated soluble CD27 levels in serum of patients with systemic lupus erythematosus. Clinical Immunology and Immunopathology.",
        "Wiedemann A et al. (2020). CD27+ memory B cells and CD27-high plasmablasts in systemic lupus erythematosus. Frontiers in Immunology.",
        "Jacobi AM et al. (2003). Correlation between circulating CD27high plasma cells and disease activity in systemic lupus erythematosus. Arthritis & Rheumatism.",
        "Tipton CM et al. (2015). Diversity, cellular origin and autoreactivity of antibody-secreting cell expansions in acute systemic lupus erythematosus. Nature Immunology."
      ]
    },
    {
      "gene_name": "BIRC3",
      "direct_evidence": "low",
      "indirect_evidence": "medium",
      "is_stress_response_gene": false,
      "explanation": "BIRC3 (cIAP2) is an E3 ubiquitin ligase that controls canonical and non-canonical NF-kB signaling and is essential for B-cell survival downstream of BAFF/CD40; BIRC3 mutations drive B-cell lymphomagenesis. There are no validated SLE biomarker studies tying BIRC3 expression to SLEDAI or flare, but indirect mechanistic relevance is meaningful because BAFF-driven non-canonical NF-kB signaling and B-cell survival are central to SLE and to belimumab biology. logFC 0.21 is consistent with low-grade NF-kB activation across PBMCs.",
      "references": [
        "Zarnegar BJ et al. (2008). Noncanonical NF-kappaB activation requires coordinated assembly of a regulatory complex of the adaptors cIAP1, cIAP2, TRAF2 and TRAF3 and the kinase NIK. Nature Immunology.",
        "Gardam S et al. (2008). TRAF2 and TRAF3 signal adapters act cooperatively to control the maturation and survival signals delivered to B cells by the BAFF receptor. Immunity.",
        "Rosebeck S et al. (2011). Cleavage of NIK by the API2-MALT1 fusion oncoprotein leads to noncanonical NF-kappaB activation. Science."
      ]
    },
    {
      "gene_name": "CNPY3",
      "direct_evidence": "low",
      "indirect_evidence": "medium",
      "is_stress_response_gene": false,
      "explanation": "CNPY3 (canopy-3) is an ER-resident co-chaperone of gp96/HSP90B1 required for folding and trafficking of multiple Toll-like receptors, including the endosomal nucleic-acid-sensing TLR7 and TLR9 that drive SLE pathogenesis. No direct SLE biomarker validation exists for CNPY3, but its indispensable role for TLR7/9 surface and endosomal expression provides medium indirect mechanistic relevance to lupus, where TLR7 gain-of-function is now a validated driver. The logFC near zero argues against a specific transcriptional disease signature.",
      "references": [
        "Liu B et al. (2010). The chaperone CNPY3 is required for innate immune signaling by Toll-like receptors. Cell.",
        "Brown GJ et al. (2022). TLR7 gain-of-function genetic variation causes human lupus. Nature.",
        "Marshak-Rothstein A (2006). Toll-like receptors in systemic autoimmune disease. Nature Reviews Immunology."
      ]
    },
    {
      "gene_name": "AP1S2",
      "direct_evidence": "low",
      "indirect_evidence": "low",
      "is_stress_response_gene": false,
      "explanation": "AP1S2 encodes the sigma1B subunit of the AP-1 clathrin adaptor complex involved in vesicle trafficking between the trans-Golgi network and endosomes; it is unrelated to the AP-1 (Fos/Jun) transcription factor. There is no SLE biomarker literature on AP1S2, and the logFC near zero indicates a noisy attribution. Its role is generic intracellular vesicle trafficking, not stress per se nor immune-specific; it is best regarded as housekeeping with negligible direct or indirect SLE relevance.",
      "references": [
        "Saillour Y et al. (2007). Mutations in the AP1S2 gene encoding the sigma 1B subunit of AP-1 complex are associated with X-linked mental retardation. Journal of Medical Genetics.",
        "Robinson MS (2004). Adaptable adaptors for coated vesicles. Trends in Cell Biology."
      ]
    },
    {
      "gene_name": "IFI44L",
      "direct_evidence": "high",
      "indirect_evidence": "high",
      "is_stress_response_gene": false,
      "explanation": "IFI44L is one of the most robustly validated type I interferon-stimulated gene (ISG) biomarkers of SLE, contributing to the canonical IFN signature that correlates with SLEDAI, anti-dsDNA, complement consumption, and lupus nephritis activity, and it is the basis of a clinically-deployed DNA-methylation diagnostic test (Zhao et al. 2016). The very large logFC (+2.75) in this dataset is fully consistent with that established biology. It is a bona fide immune-specific antiviral gene, not a generic stress response.",
      "references": [
        "Bauer JW et al. (2009). Interferon-regulated chemokines as biomarkers of systemic lupus erythematosus disease activity: a validation study. Arthritis & Rheumatism.",
        "Zhao M et al. (2016). IFI44L promoter methylation as a blood biomarker for systemic lupus erythematosus. Annals of the Rheumatic Diseases.",
        "Baechler EC et al. (2003). Interferon-inducible gene expression signature in peripheral blood cells of patients with severe lupus. Proceedings of the National Academy of Sciences USA.",
        "Kirou KA et al. (2005). Activation of the interferon-alpha pathway identifies a subgroup of SLE patients with distinct serologic features. Arthritis & Rheumatism."
      ]
    },
    {
      "gene_name": "CEBPB",
      "direct_evidence": "low",
      "indirect_evidence": "medium",
      "is_stress_response_gene": false,
      "explanation": "CEBPB (C/EBP-beta) is a basic leucine-zipper transcription factor central to myeloid differentiation, emergency granulopoiesis, and IL-6/acute-phase gene induction. Direct SLEDAI/flare biomarker validation is lacking, but indirect mechanistic relevance to SLE is meaningful through its control of inflammatory monocyte/neutrophil programs and IL-6 transcription, both prominent in active lupus and lupus nephritis. logFC 0.18 is consistent with low-grade myeloid inflammatory tone.",
      "references": [
        "Akira S et al. (1990). A nuclear factor for IL-6 expression (NF-IL6) is a member of a C/EBP family. EMBO Journal.",
        "Hirai H et al. (2006). C/EBPbeta is required for emergency granulopoiesis. Nature Immunology.",
        "Huber R et al. (2012). Regulation of C/EBPbeta and resulting functions in cells of the monocytic lineage. Cellular Signalling."
      ]
    }
  ]
}
