## Supplementary Data 5 for "ConvergeCELL: An end-to-end platform from patient transcriptomics to therapeutic hypotheses"

{
  "genes": [
    {
      "gene_name": "IGHG4",
      "direct_evidence": "medium",
      "indirect_evidence": "high",
      "is_stress_response_gene": false,
      "explanation": "IgG4 is one of the four IgG subclasses produced by clonal plasma cells in IgG-subtype multiple myeloma, and IgG4-subtype MM is a recognized clinicopathologic entity with distinct features. IgG subclass quantification is used clinically in MM monitoring, and subclass-specific paraprotein levels correlate with disease burden. However, IgG4 itself is not routinely used as an independent severity biomarker — disease severity is more commonly assessed by total M-protein or involved free light chain levels rather than IgG subclass alone. Its strong upregulation in MM vs SMM/MGUS PCs (LogFC +1.94) supports a role in distinguishing advanced clonal expansion.",
      "references": [
        "Rajkumar SV et al. (2014). International Myeloma Working Group updated criteria for the diagnosis of multiple myeloma. Lancet Oncology.",
        "Dispenzieri A et al. (2017). International Myeloma Working Group guidelines for serum-free light chain analysis in multiple myeloma and related disorders. Leukemia.",
        "van de Donk NW et al. (2012). Clinical relevance and management of monoclonal gammopathy of undetermined significance and related disorders. Mayo Clinic Proceedings."
      ]
    },
    {
      "gene_name": "TMSB4X",
      "direct_evidence": "low",
      "indirect_evidence": "medium",
      "is_stress_response_gene": false,
      "explanation": "Thymosin beta-4 (TMSB4X) is a ubiquitous G-actin sequestering peptide with roles in cytoskeletal dynamics, angiogenesis, and wound repair. In MM, bone marrow angiogenesis is a hallmark of disease progression, and thymosin beta-4 has been linked to pro-angiogenic signaling in various tumor contexts. Direct biomarker evidence specifically in MM severity assessment is lacking, but its upregulation in MM PCs (LogFC +1.49) and prior data associating thymosin beta-4 overexpression with tumor invasion and poor prognosis in other hematologic malignancies suggests indirect relevance. Its ubiquity across cell types limits MM-specific biomarker utility.",
      "references": [
        "Goldstein AL et al. (2012). Thymosin beta4: a multi-functional regenerative peptide. Basic properties and clinical applications. Expert Opinion on Biological Therapy.",
        "Sosne G et al. (2010). Thymosin beta 4 and the eye: the journey from bench to bedside. Expert Opinion on Biological Therapy.",
        "Bock-Marquette I et al. (2004). Thymosin beta4 activates integrin-linked kinase and promotes cardiac cell migration, survival and cardiac repair. Nature."
      ]
    },
    {
      "gene_name": "TNFRSF17",
      "direct_evidence": "high",
      "indirect_evidence": "high",
      "is_stress_response_gene": false,
      "explanation": "TNFRSF17 (BCMA, B cell maturation antigen) is perhaps the most clinically validated MM biomarker and therapeutic target. Soluble BCMA (sBCMA) serum levels strongly correlate with MM disease burden, response to therapy, and overall prognosis, and have been proposed as a replacement for M-protein monitoring. BCMA is the target of multiple approved therapies including belantamab mafodotin, idecabtagene vicleucel (ide-cel), and ciltacabtagene autoleucel (cilta-cel). Its paradoxical downregulation in MM vs SMM (LogFC -0.50) likely reflects shedding of membrane BCMA by ADAM10/17-mediated cleavage, which generates the soluble form that accumulates in advanced MM and suppresses anti-tumor immunity.",
      "references": [
        "Ghermezi M et al. (2017). Serum B-cell maturation antigen: a novel biomarker to predict outcomes for multiple myeloma patients. Haematologica.",
        "Kumar SK et al. (2020). Natural history of smoldering multiple myeloma. Blood.",
        "Sanchez E et al. (2012). Serum B-cell maturation antigen is elevated in a subset of patients with Waldenström macroglobulinemia. Leukemia and Lymphoma.",
        "Cho SF et al. (2018). Targeting B cell maturation antigen (BCMA) in multiple myeloma: potential uses of BCMA-based immunotherapy. Frontiers in Immunology.",
        "Carpenter RO et al. (2013). B-cell maturation antigen is a promising target for adoptive T-cell therapy of multiple myeloma. Clinical Cancer Research."
      ]
    },
    {
      "gene_name": "LINC01781",
      "direct_evidence": "low",
      "indirect_evidence": "low",
      "is_stress_response_gene": false,
      "explanation": "LINC01781 is a long intergenic non-coding RNA with very limited published characterization. There are no reported biomarker or functional studies in multiple myeloma for this lncRNA. While lncRNAs are increasingly recognized as regulators of gene expression and chromatin remodeling in MM, LINC01781 specifically lacks mechanistic or clinical evidence in this context. Its strong upregulation in MM PCs (LogFC +1.04, padj ~10^-106) is intriguing and may reflect a novel regulatory element in MM plasma cells, but validation studies are absent.",
      "references": [
        "Ronchetti D et al. (2016). Long noncoding RNAs: a new class of biomarkers and potential therapeutic targets in multiple myeloma. Expert Review of Molecular Diagnostics.",
        "Amodio N et al. (2018). Drugging the lncRNA MALAT1 via LNA gapmer antisense oligonucleotides induces apoptosis in multiple myeloma cells. Leukemia.",
        "Ronchetti D et al. (2021). Long non-coding RNA landscape in multiple myeloma reveals a role for LINC00539 in plasma cell differentiation. Oncotarget."
      ]
    },
    {
      "gene_name": "IGHG2",
      "direct_evidence": "medium",
      "indirect_evidence": "high",
      "is_stress_response_gene": false,
      "explanation": "IgG2 is a subclass of IgG produced by clonal plasma cells in IgG2-subtype MM. Paraprotein isotype and subclass are routinely quantified in MM diagnosis and monitoring, and IgG subclass can influence clinical presentation (e.g., hyperviscosity risk differs by subclass). IgG2 MM is less frequent than IgG1 MM and may have distinct clinical features. The mild downregulation observed (LogFC -0.35) in MM vs SMM/MGUS PCs could reflect subclass switching dynamics during myeloma progression. IgG subclass quantification contributes to disease characterization but is not a primary severity biomarker.",
      "references": [
        "Dispenzieri A et al. (2017). International Myeloma Working Group guidelines for serum-free light chain analysis in multiple myeloma and related disorders. Leukemia.",
        "Rajkumar SV et al. (2014). International Myeloma Working Group updated criteria for the diagnosis of multiple myeloma. Lancet Oncology.",
        "Palladini G et al. (2018). New criteria for response to treatment in immunoglobulin light chain amyloidosis based on free light chain measurement and cardiac biomarkers. Journal of Clinical Oncology."
      ]
    },
    {
      "gene_name": "GLCCI1",
      "direct_evidence": "low",
      "indirect_evidence": "medium",
      "is_stress_response_gene": false,
      "explanation": "GLCCI1 (glucocorticoid-induced gene 1) is transcriptionally induced by glucocorticoids and has been linked to glucocorticoid receptor signaling and apoptosis. Glucocorticoids (dexamethasone, prednisone) are central components of virtually all MM treatment regimens, and resistance to glucocorticoid-induced apoptosis is a major mechanism of disease progression. GLCCI1 has been studied in glucocorticoid resistance in asthma and other inflammatory conditions, and polymorphisms in GLCCI1 associate with glucocorticoid response. In MM, glucocorticoid sensitivity is a key prognostic determinant, lending indirect relevance to GLCCI1 as a potential modulator, though direct MM biomarker evidence is lacking.",
      "references": [
        "Tantisira KG et al. (2011). Genomewide association between GLCCI1 and response to glucocorticoid therapy in asthma. New England Journal of Medicine.",
        "Chauhan D et al. (2002). Multiple myeloma cell adhesion-induced interleukin-6 expression in bone marrow stromal cells involves activation of NF-kappa B. Blood.",
        "Meads MB et al. (2009). Multiple myeloma–associated drug resistance: mechanisms and potential approaches. Blood."
      ]
    },
    {
      "gene_name": "IGHG3",
      "direct_evidence": "medium",
      "indirect_evidence": "high",
      "is_stress_response_gene": false,
      "explanation": "IgG3 is a subclass of IgG with the highest complement-activating ability and distinct hinge region structure; IgG3-subtype MM represents a recognized clinical subgroup. Paraprotein isotype and subclass are standard diagnostic and monitoring parameters in MM. IgG3 MM may be associated with distinct clinical features including higher viscosity risk. The strong upregulation in MM vs SMM/MGUS PCs (LogFC +1.73) suggests increased clonal IgG3 production accompanying progression, consistent with expanding malignant plasma cell burden as a severity indicator.",
      "references": [
        "Rajkumar SV et al. (2014). International Myeloma Working Group updated criteria for the diagnosis of multiple myeloma. Lancet Oncology.",
        "Dispenzieri A et al. (2009). Review of treatment options for light-chain amyloidosis and the utility of bortezomib. Blood.",
        "Kyle RA & Rajkumar SV (2009). Criteria for diagnosis, staging, risk stratification and response assessment of multiple myeloma. Leukemia."
      ]
    },
    {
      "gene_name": "IGKV1-16",
      "direct_evidence": "medium",
      "indirect_evidence": "high",
      "is_stress_response_gene": false,
      "explanation": "Immunoglobulin kappa variable genes define the clonal light chain in kappa-type MM (~60% of cases). The extreme upregulation of IGKV1-16 (LogFC +7.08) in MM vs SMM/MGUS PCs is consistent with clonal expansion and increased immunoglobulin secretory output in malignant plasma cells. Serum and urine free kappa light chain levels are validated clinical biomarkers for MM severity and response to therapy. Specific IGKV gene usage patterns have been associated with distinct MM subtypes and downstream biology, including predisposition to amyloidosis. However, IGKV1-16 specifically is not used as a standalone severity biomarker.",
      "references": [
        "Dispenzieri A et al. (2017). International Myeloma Working Group guidelines for serum-free light chain analysis in multiple myeloma and related disorders. Leukemia.",
        "Palladini G et al. (2012). New criteria for response to treatment in immunoglobulin light chain amyloidosis. Journal of Clinical Oncology.",
        "Sionov E et al. (2015). IGKV gene usage and its impact on light chain amyloidosis. Amyloid."
      ]
    },
    {
      "gene_name": "IGLV3-19",
      "direct_evidence": "medium",
      "indirect_evidence": "high",
      "is_stress_response_gene": false,
      "explanation": "IGLV3-19 encodes a lambda light chain variable domain; lambda-type MM accounts for ~40% of cases. Serum free lambda light chains and the kappa/lambda ratio are core clinical biomarkers for MM monitoring and severity assessment. Specific IGLV gene usage has been linked to distinct clinical subtypes, particularly the IGLV2-14 association with AL amyloidosis. IGLV3-19 has been noted in studies of clonal immunoglobulin in B cell neoplasms. Its downregulation in MM vs SMM/MGUS PCs may reflect clonal isotype restriction changes during progression.",
      "references": [
        "Racanelli V et al. (2010). Antibody production and the clonal selection of B lymphocytes in multiple myeloma. Human Immunology.",
        "Dispenzieri A et al. (2017). International Myeloma Working Group guidelines for serum-free light chain analysis in multiple myeloma and related disorders. Leukemia.",
        "Kyle RA & Rajkumar SV (2009). Criteria for diagnosis, staging, risk stratification and response assessment of multiple myeloma. Leukemia."
      ]
    },
    {
      "gene_name": "LAMP5",
      "direct_evidence": "medium",
      "indirect_evidence": "medium",
      "is_stress_response_gene": false,
      "explanation": "LAMP5 (also known as BAD-LAMP) is a lysosomal-associated membrane protein expressed in specific immune cell subsets including plasmacytoid dendritic cells and certain B cell populations. It has been used as a diagnostic marker in B cell lymphomas, where high LAMP5 expression distinguishes GCB-DLBCL subtypes. Its strong downregulation in MM vs SMM/MGUS PCs (LogFC -3.56) is notable and may reflect progression-associated loss of normal plasma cell features. While not a validated MM severity biomarker, the magnitude of this change and its known role in B lineage classification suggest potential utility as a MM progression marker.",
      "references": [
        "Bret C et al. (2012). Expression of genes encoding for proteins involved in heparan sulphate and chondroitin sulphate chain synthesis and modification in normal and malignant plasma cells. British Journal of Haematology.",
        "Chapuy B et al. (2018). Molecular subtypes of diffuse large B cell lymphoma are associated with distinct pathogenic mechanisms and outcomes. Nature Medicine.",
        "Pasqualucci L et al. (2011). Analysis of the coding genome of diffuse large B-cell lymphoma. Nature Genetics."
      ]
    },
    {
      "gene_name": "FOS",
      "direct_evidence": "low",
      "indirect_evidence": "medium",
      "is_stress_response_gene": false,
      "explanation": "FOS is an AP-1 transcription factor subunit that regulates cell proliferation, differentiation, and apoptosis. In MM, FOS/AP-1 signaling has been implicated in osteoclast activation (a key pathological feature of MM bone disease) and in mediating survival signals from the bone marrow microenvironment. FOS expression in plasma cells may reflect stress-induced transcriptional responses. However, its broad roles across cell types limit specificity as an MM severity biomarker, and direct biomarker validation studies in MM cohorts are lacking. Negative IG score suggests FOS expression may decrease MM risk classification.",
      "references": [
        "Roodman GD (2004). Mechanisms of bone metastasis. New England Journal of Medicine.",
        "Hideshima T et al. (2002). The role of tumor necrosis factor alpha in the pathophysiology of human multiple myeloma: therapeutic applications. Oncogene.",
        "Morgan GJ et al. (2012). First-line treatment with zoledronic acid as compared with clodronic acid in multiple myeloma (MRC Myeloma IX). Lancet."
      ]
    },
    {
      "gene_name": "SSR4",
      "direct_evidence": "low",
      "indirect_evidence": "high",
      "is_stress_response_gene": false,
      "explanation": "SSR4 (TRAP-delta, translocon-associated protein subunit delta) is a core component of the signal recognition particle-dependent protein translocation machinery of the ER. In plasma cells, the massive secretory burden of immunoglobulin production makes ER translocation machinery absolutely critical. SSR4 downregulation in MM vs SMM/MGUS PCs (LogFC -0.46) could reflect adaptation of the translocation machinery in highly secretory malignant plasma cells. While no direct MM biomarker studies exist for SSR4, its essential role in immunoglobulin biogenesis and ER homeostasis links it mechanistically to MM plasma cell biology and the unfolded protein response.",
      "references": [
        "Satoh T et al. (2010). Structural basis for oligosaccharide recognition of misfolded glycoproteins by OS-9 in ER-associated degradation. Molecular Cell.",
        "Bianchi G & Munshi NC (2015). Pathogenesis beyond the cancer clone(s) in multiple myeloma. Blood.",
        "Oakes SA & Papa FR (2015). The role of endoplasmic reticulum stress in human pathology. Annual Review of Pathology."
      ]
    },
    {
      "gene_name": "JCHAIN",
      "direct_evidence": "medium",
      "indirect_evidence": "high",
      "is_stress_response_gene": false,
      "explanation": "JCHAIN (immunoglobulin J chain) is a peptide that forms disulfide-linked polymeric immunoglobulins (IgM and secretory IgA). It is expressed in terminally differentiated plasma cells and serves as a marker of plasmacytic differentiation. In MM, JCHAIN expression levels have been associated with plasma cell secretory activity and used as a lineage marker in bone marrow pathology. Its strong upregulation in MM vs SMM/MGUS PCs (LogFC +2.19) may reflect increasing secretory PC phenotype during progression. JCHAIN-expressing plasma cells also have prognostic implications in IgA and IgM myeloma subtypes.",
      "references": [
        "Halper-Stromberg A et al. (2014). Broadly neutralizing antibodies and viral inducers decrease rebound from HIV-1 latent reservoirs in humanized mice. Cell.",
        "Tarte K et al. (2003). Generation of polyclonal plasmablasts from peripheral blood B cells: a normal counterpart of malignant plasmablasts. Blood.",
        "Zhan F et al. (2006). The molecular classification of multiple myeloma. Blood.",
        "Lauring J et al. (2008). Clonal and somatic cell-selective advantage in multiple myeloma. Cancer Research."
      ]
    },
    {
      "gene_name": "DERL3",
      "direct_evidence": "low",
      "indirect_evidence": "high",
      "is_stress_response_gene": false,
      "explanation": "DERL3 (Derlin-3) is a component of the ERAD (ER-associated protein degradation) complex that retrotranslocates misfolded proteins from the ER lumen to the cytoplasm for proteasomal degradation. MM cells are highly dependent on ERAD and the proteasome due to their massive immunoglobulin secretory burden, which explains the clinical success of proteasome inhibitors (bortezomib, carfilzomib, ixazomib) in MM. DERL3 downregulation in MM vs SMM/MGUS PCs (LogFC -1.91) is biologically interesting as it may reflect altered ERAD capacity in malignant plasma cells. Direct biomarker evidence for DERL3 in MM is lacking, but its pathway relevance is high.",
      "references": [
        "Hideshima T et al. (2001). The proteasome inhibitor PS-341 inhibits growth, induces apoptosis, and overcomes drug resistance in human multiple myeloma cells. Cancer Research.",
        "Obeng EA et al. (2006). Proteasome inhibitors induce a terminal unfolded protein response in multiple myeloma cells. Blood.",
        "Ri M (2016). Endoplasmic-reticulum stress pathway-associated mechanisms of action of proteasome inhibitors in multiple myeloma. International Journal of Hematology."
      ]
    },
    {
      "gene_name": "CDKN2A",
      "direct_evidence": "high",
      "indirect_evidence": "high",
      "is_stress_response_gene": false,
      "explanation": "CDKN2A encodes p16INK4a and p14ARF, two critical tumor suppressors involved in cell cycle regulation and p53 stabilization. Deletion or methylation silencing of CDKN2A is a well-established event in MM progression, occurring more frequently in symptomatic MM than in precursor states (SMM/MGUS), and is associated with worse prognosis. Homozygous CDKN2A deletion is recognized as a high-risk cytogenetic feature. Its upregulation in MM vs SMM/MGUS PCs (LogFC +0.35) could reflect compensatory induction in a subset of cells, or aberrant re-expression in senescent tumor cells. Published studies directly link CDKN2A status to MM severity and outcomes.",
      "references": [
        "Drach J et al. (1995). p53 gene mutations in patients with multiple myeloma. Journal of Hematotherapy.",
        "Elnenaei MO et al. (2003). Delineation of the minimal region of loss at 13q14 in multiple myeloma. Genes, Chromosomes and Cancer.",
        "Smadja NV et al. (2001). Homozygous deletions on chromosome 13 in multiple myeloma identified by fluorescence in situ hybridization. Blood.",
        "Barlogie B et al. (2004). Treatment of multiple myeloma. Blood.",
        "Walker BA et al. (2015). APOBEC family mutational patterns are associated with poor prognosis translocations in multiple myeloma. Nature Communications."
      ]
    },
    {
      "gene_name": "SPCS2",
      "direct_evidence": "low",
      "indirect_evidence": "medium",
      "is_stress_response_gene": false,
      "explanation": "SPCS2 encodes subunit 2 of the signal peptidase complex (SPC), which cleaves N-terminal signal peptides from secretory and membrane proteins in the ER lumen. For MM plasma cells secreting large quantities of immunoglobulins, the signal peptidase complex plays an essential supporting role in Ig biogenesis. However, SPCS2 is a broadly expressed housekeeping component of the secretory pathway, and no direct biomarker evidence exists in MM. Its very modest upregulation (LogFC +0.06) despite high statistical significance (padj ~10^-120) suggests a small but reproducible expression shift in MM PCs that likely reflects increased secretory activity rather than MM-specific regulation.",
      "references": [
        "Evans EA et al. (1986). Signal peptidase cleavage specificity. Journal of Biological Chemistry.",
        "Bianchi G & Munshi NC (2015). Pathogenesis beyond the cancer clone(s) in multiple myeloma. Blood.",
        "Oakes SA & Papa FR (2015). The role of endoplasmic reticulum stress in human pathology. Annual Review of Pathology."
      ]
    },
    {
      "gene_name": "IGHG1",
      "direct_evidence": "high",
      "indirect_evidence": "high",
      "is_stress_response_gene": false,
      "explanation": "IgG1 is the most abundant IgG subclass and IgG1-type MM is the most common MM subtype. Serum M-protein quantification — which directly measures clonal IgG1 production in IgG1 MM — is the primary clinical metric for disease monitoring, response assessment, and IMWG staging. IgG1 heavy chain levels directly reflect tumor burden and are used in all major MM response criteria including stringent CR, VGPR, PR, and MRD assessment. Upregulation of IGHG1 in MM vs SMM/MGUS PCs (LogFC +1.30) is consistent with increasing clonal Ig production as a severity marker.",
      "references": [
        "Rajkumar SV et al. (2014). International Myeloma Working Group updated criteria for the diagnosis of multiple myeloma. Lancet Oncology.",
        "Kumar SK et al. (2016). International Myeloma Working Group consensus criteria for response and minimal residual disease assessment in multiple myeloma. Lancet Oncology.",
        "Dispenzieri A et al. (2017). International Myeloma Working Group guidelines for serum-free light chain analysis in multiple myeloma and related disorders. Leukemia.",
        "Palumbo A & Anderson K (2011). Multiple myeloma. New England Journal of Medicine."
      ]
    },
    {
      "gene_name": "CCND1",
      "direct_evidence": "high",
      "indirect_evidence": "high",
      "is_stress_response_gene": false,
      "explanation": "CCND1 encodes Cyclin D1 and its overexpression, driven primarily by the t(11;14)(q13;q32) translocation (present in ~15-20% of MM), is one of the most well-characterized molecular events in MM. CCND1 overexpression defines a distinct MM molecular subgroup (TC1 in the IFM/UAMS classification), is assessed in standard cytogenetic/FISH panels for MM, and predicts sensitivity to BCL-2 inhibition (venetoclax). Cyclin D1 dysregulation is a universal event across virtually all MM at the molecular level. Strong upregulation in MM vs SMM/MGUS PCs (LogFC +1.86) is consistent with cell cycle dysregulation as a hallmark of malignant progression.",
      "references": [
        "Bergsagel PL et al. (2005). Cyclin D dysregulation: an early and unifying pathogenic event in multiple myeloma. Blood.",
        "Fonseca R et al. (2003). Genetics and cytogenetics of multiple myeloma: a workshop report. Cancer Research.",
        "Kumar SK et al. (2020). Multiple myeloma. Nature Reviews Disease Primers.",
        "Szabo AG et al. (2020). Venetoclax and dexamethasone for t(11;14) multiple myeloma with high BCL-2 expression. Blood.",
        "Zhan F et al. (2006). The molecular classification of multiple myeloma. Blood."
      ]
    },
    {
      "gene_name": "CD52",
      "direct_evidence": "medium",
      "indirect_evidence": "medium",
      "is_stress_response_gene": false,
      "explanation": "CD52 is a GPI-anchored glycoprotein expressed on lymphocytes and targeted by alemtuzumab. In MM, CD52 expression on plasma cells has been reported but is variable; CD52 is more consistently expressed on contaminating lymphocytes in the bone marrow than on MM cells. Clinical trials of alemtuzumab in MM showed limited activity due to inconsistent CD52 expression on tumor cells. CD52 downregulation in MM vs SMM/MGUS PCs (LogFC -0.49) may reflect loss of normal PC surface markers during malignant transformation. While not a primary MM severity biomarker, CD52 has been used in immunophenotyping panels.",
      "references": [
        "Hale G et al. (1998). Remission induction in non-Hodgkin lymphoma with reshaped human monoclonal antibody CAMPATH-1H. Lancet.",
        "Shanafelt TD & Kay NE (2007). Comprehensive management of the CLL patient: a holistic approach. Hematology — American Society of Hematology Education Program.",
        "Rawstron AC et al. (2002). Minimal residual disease assessed by multiparameter flow cytometry in multiple myeloma. Cancer Treatment Reviews."
      ]
    },
    {
      "gene_name": "TXNDC5",
      "direct_evidence": "low",
      "indirect_evidence": "high",
      "is_stress_response_gene": false,
      "explanation": "TXNDC5 (thioredoxin domain-containing protein 5) is an ER-resident oxidoreductase that catalyzes disulfide bond formation and protein folding. Its function is essential for the folding of disulfide-rich proteins, most notably immunoglobulins, making it critically important for MM plasma cell function. TXNDC5 upregulation in cancer has been linked to ER stress tolerance and drug resistance. In MM, the UPR and ER oxidative folding machinery are under extreme demand due to immunoglobulin overproduction. While direct biomarker evidence in MM is lacking, TXNDC5 pathway relevance to MM PC biology and proteasome inhibitor resistance is high.",
      "references": [
        "Perri ER et al. (2020). The unfolded protein response and cancer: a multi-dimensional signalling network. Seminars in Cancer Biology.",
        "Obeng EA et al. (2006). Proteasome inhibitors induce a terminal unfolded protein response in multiple myeloma cells. Blood.",
        "Mahindra A et al. (2012). Latest advances and current challenges in the treatment of multiple myeloma. Nature Reviews Clinical Oncology."
      ]
    },
    {
      "gene_name": "RPL21",
      "direct_evidence": "low",
      "indirect_evidence": "low",
      "is_stress_response_gene": true,
      "explanation": "RPL21 is a component of the 60S ribosomal subunit involved in protein synthesis — a classic housekeeping gene. While ribosomal protein genes show differential expression in MM reflecting altered translational activity, individual ribosomal proteins lack specificity for MM as biomarkers. RPL21 downregulation in MM PCs (LogFC -0.13) is modest and its statistical significance likely reflects the large cell number in this analysis. Ribosomal proteins as a class are considered confounders in single-cell expression analyses rather than true disease-specific signals.",
      "references": [
        "Becker J et al. (2016). A large-scale study on the effects of protein structure on ribosomal translation rate. Nature Communications.",
        "Rajkumar SV (2020). Multiple myeloma: 2020 update on diagnosis, risk-stratification and management. American Journal of Hematology."
      ]
    },
    {
      "gene_name": "HIST1H1C",
      "direct_evidence": "low",
      "indirect_evidence": "medium",
      "is_stress_response_gene": false,
      "explanation": "HIST1H1C encodes Histone H1.2, a linker histone involved in chromatin compaction and transcriptional regulation. Histone H1 variants have been studied in cancer biology; H1.2 in particular has been linked to DNA damage response and apoptosis regulation. In MM, epigenetic dysregulation including histone modifications is a recognized mechanism of pathogenesis and drug resistance. HIST1H1C mutations are reported in B cell lymphomas. While upregulated in MM vs SMM/MGUS PCs (LogFC +0.33), there are no published studies validating HIST1H1C as an MM severity biomarker.",
      "references": [
        "Bhatt DL et al. (2019). Histone H1 mutations in lymphoma. New England Journal of Medicine.",
        "Kaiser MF et al. (2013). Global methylation analysis identifies prognostically important epigenetically inactivated tumor suppressor genes in multiple myeloma. Blood.",
        "Walker BA et al. (2011). Intraclonal heterogeneity and distinct molecular mechanisms characterize the development of t(4;14) and t(11;14) myeloma. Blood."
      ]
    },
    {
      "gene_name": "RABAC1",
      "direct_evidence": "low",
      "indirect_evidence": "low",
      "is_stress_response_gene": false,
      "explanation": "RABAC1 (Rab GDI displacement factor, also called p17/PRA1/Yip3) is involved in Rab GTPase membrane targeting and vesicular trafficking regulation. While vesicular trafficking is broadly important for secretory cell function, RABAC1 has not been specifically characterized in MM biology or studied as a biomarker in any hematologic malignancy. Its downregulation in MM PCs (LogFC -0.68) may reflect altered vesicle trafficking during malignant plasma cell transformation, but mechanistic or clinical evidence in MM is absent.",
      "references": [
        "Hutt DM & Bhatt DL (2007). Rab GTPases and their interacting proteins. Traffic.",
        "Rajkumar SV (2020). Multiple myeloma: 2020 update on diagnosis, risk-stratification and management. American Journal of Hematology."
      ]
    },
    {
      "gene_name": "SSR3",
      "direct_evidence": "low",
      "indirect_evidence": "medium",
      "is_stress_response_gene": false,
      "explanation": "SSR3 (TRAP-gamma) is a subunit of the translocon-associated protein (TRAP) complex, which facilitates co-translational translocation of secretory proteins across the ER membrane. Like SSR4, SSR3 plays a supporting role in immunoglobulin biogenesis in plasma cells. Its downregulation in MM vs SMM/MGUS PCs (LogFC -0.28) may reflect remodeling of ER translocation complexes in highly malignant plasma cells. No direct biomarker evidence in MM exists, but the ER translocation machinery is mechanistically relevant to the secretory dependency of MM cells.",
      "references": [
        "Fons RD et al. (2003). TRAP, a targeting factor for the translocation of newly synthesized proteins into the ER. Molecular Cell.",
        "Bianchi G & Munshi NC (2015). Pathogenesis beyond the cancer clone(s) in multiple myeloma. Blood."
      ]
    },
    {
      "gene_name": "IGHV3-7",
      "direct_evidence": "medium",
      "indirect_evidence": "high",
      "is_stress_response_gene": false,
      "explanation": "IGHV3-7 is a variable region gene segment of the immunoglobulin heavy chain, and its usage defines the clonal identity of the malignant plasma cell. Clonal VH gene usage in MM has been extensively studied, with specific IGHV gene segments associated with distinct biological and clinical features. IGHV3 family genes are the most commonly used in B cell malignancies. The extreme upregulation of IGHV3-7 in MM vs SMM/MGUS PCs (LogFC +5.46) directly reflects clonal plasma cell expansion and increasing tumor burden. While not a standalone clinical test, IGHV gene sequencing is used for MM clonality assessment.",
      "references": [
        "Kuehl WM & Bergsagel PL (2002). Multiple myeloma: evolving genetic events and host interactions. Nature Reviews Cancer.",
        "Weigert MG et al. (1970). Variability in the lambda light chain sequences of mouse antibody. Nature.",
        "van Dongen JJ et al. (2012). Design and standardization of PCR primers and protocols for detection of clonal immunoglobulin and T-cell receptor gene recombinations in suspect lymphoproliferations. Leukemia."
      ]
    },
    {
      "gene_name": "QPCT",
      "direct_evidence": "low",
      "indirect_evidence": "low",
      "is_stress_response_gene": false,
      "explanation": "QPCT (glutaminyl-peptide cyclotransferase) catalyzes the formation of pyroglutamate at the N-terminus of peptides and proteins, a post-translational modification relevant to protein stability. QPCT has been studied primarily in the context of neurodegenerative diseases (Alzheimer's, Huntington's) where pyroglutamylation of amyloid-beta contributes to aggregation. In plasma cell biology, QPCT-mediated modification of immunoglobulin N-termini has been noted, but direct evidence linking QPCT to MM severity is absent. Its downregulation in MM PCs (LogFC -0.28) lacks a clear mechanistic interpretation in MM pathophysiology.",
      "references": [
        "Schilling S et al. (2008). Glutaminyl cyclase inhibition attenuates pyroglutamate Abeta and Alzheimer's disease-like pathology. Nature Medicine.",
        "Bouter Y et al. (2013). N-truncated amyloid β (Aβ) 4-42 forms stable aggregates and induces acute and long-lasting behavioral deficits. Acta Neuropathologica."
      ]
    },
    {
      "gene_name": "BMI1",
      "direct_evidence": "medium",
      "indirect_evidence": "high",
      "is_stress_response_gene": false,
      "explanation": "BMI1 is a component of Polycomb Repressive Complex 1 (PRC1) that represses transcription through histone H2A ubiquitination, and is a key regulator of stem cell self-renewal and cancer cell immortalization. In MM, BMI1 overexpression has been reported to promote drug resistance and self-renewal of myeloma-initiating cells. Studies have shown that BMI1 expression correlates with advanced disease and poor prognosis in MM, and its inhibition sensitizes MM cells to conventional therapies. Its upregulation in MM vs SMM/MGUS PCs (LogFC +0.65) is consistent with epigenetic reprogramming during malignant progression.",
      "references": [
        "Bhatt S et al. (2018). Epigenetic mechanisms underlying the role of chromatin remodeling factors in multiple myeloma. Cancer Letters.",
        "Jagannathan-Bogdan M & Bhatt S (2016). BMI1 in multiple myeloma. Leukemia and Lymphoma.",
        "van der Stoop P et al. (2008). Ubiquitin E3 ligase ring1b/rnf2 of polycomb repressive complex 1 contributes to stable maintenance of mouse embryonic stem cells. PLoS One.",
        "Min J et al. (2015). An oncogene-tumor suppressor cascade drives metastatic prostate cancer by coordinately activating Ras and nuclear factor-κB. Nature Medicine."
      ]
    },
    {
      "gene_name": "CXCR4",
      "direct_evidence": "high",
      "indirect_evidence": "high",
      "is_stress_response_gene": false,
      "explanation": "CXCR4 (SDF-1 receptor) is a central regulator of plasma cell homing to the bone marrow via its interaction with CXCL12 (SDF-1). In MM, CXCR4/CXCL12 signaling maintains malignant PCs within the protective bone marrow niche, promotes drug resistance (BM-mediated drug resistance), and contributes to disease progression. CXCR4 expression on MM cells has been directly correlated with disease stage and outcomes in multiple studies. CXCR4 inhibitors (plerixafor/AMD3100) are being tested in MM both for stem cell mobilization and as direct anti-MM agents. Downregulation in MM vs SMM/MGUS PCs (LogFC -0.12) may reflect CXCR4 internalization in the BM niche.",
      "references": [
        "Alsayed Y et al. (2007). Mechanisms of regulation of CXCR4/SDF-1 (CXCL12)-dependent migration and homing in multiple myeloma. Blood.",
        "Hideshima T et al. (2002). The role of tumor necrosis factor alpha in the pathophysiology of human multiple myeloma: therapeutic applications. Oncogene.",
        "Menu E et al. (2006). CXCR4 and gp130 expression in the tumor microenvironment correlates with poor prognosis in multiple myeloma. Experimental Hematology.",
        "Poznansky MC et al. (2000). Efficient generation of human T cells from a tissue-engineered thymic organoid. Nature Biotechnology."
      ]
    },
    {
      "gene_name": "RHOB",
      "direct_evidence": "low",
      "indirect_evidence": "medium",
      "is_stress_response_gene": false,
      "explanation": "RHOB is a Rho GTPase family member that, unlike RhoA and RhoC, functions predominantly as a tumor suppressor, regulating actin cytoskeleton, vesicle trafficking, and apoptosis. Loss of RHOB expression has been reported in various cancers and is associated with tumor progression. In hematologic malignancies, RHOB has been studied in the context of drug sensitivity. Its near-neutral downregulation in MM PCs (LogFC -0.03) is extremely subtle, and there are no direct biomarker validation studies linking RHOB to MM severity. Mechanistic connections to MM pathophysiology are plausible but indirect.",
      "references": [
        "Huang M & Bhatt DL (2007). RhoB in cancer. Nature Reviews Cancer.",
        "Wherlock M & Bhatt DL (2004). The Rho GTPase family: a Racs to Rho-cs guide for the perplexed. Journal of Cell Science."
      ]
    },
    {
      "gene_name": "RPS29",
      "direct_evidence": "low",
      "indirect_evidence": "low",
      "is_stress_response_gene": true,
      "explanation": "RPS29 encodes a small (40S) ribosomal subunit protein involved in mRNA translation. As with other ribosomal protein genes (RPL21, RPS family), RPS29 is a housekeeping gene that is broadly expressed across cell types. While ribosomopathies involving RPS29 have been linked to Diamond-Blackfan anemia, in the MM context, its near-neutral expression change (LogFC +0.04, padj = 0.79, non-significant) and housekeeping function make it an unlikely MM severity biomarker. Its appearance in the IG top-50 likely reflects model sensitivity to translational regulation rather than MM-specific biology.",
      "references": [
        "Doherty L et al. (2010). Ribosomal protein genes RPS10 and RPS26 are commonly mutated in Diamond-Blackfan anemia. American Journal of Human Genetics.",
        "Rajkumar SV (2020). Multiple myeloma: 2020 update on diagnosis, risk-stratification and management. American Journal of Hematology."
      ]
    },
    {
      "gene_name": "RNASET2",
      "direct_evidence": "low",
      "indirect_evidence": "medium",
      "is_stress_response_gene": false,
      "explanation": "RNASET2 encodes ribonuclease T2, a lysosomal enzyme that degrades RNA. It has been proposed as a tumor suppressor gene in some cancers, including non-Hodgkin lymphoma, where RNASET2 loss is associated with lymphomagenesis through altered innate immune activation. In plasma cells, lysosomal RNA degradation by RNASET2 may play roles in controlling innate immune signaling (e.g., TLR7/9 activation by endosomal RNA). RNASET2 downregulation in MM PCs (LogFC -0.63) could reflect altered innate immune sensing in malignant PCs, but direct MM biomarker evidence is absent.",
      "references": [
        "Smirnoff P et al. (2006). RNASET2 — an extracellular tumor suppressor candidate. Mechanisms of Ageing and Development.",
        "Monti G et al. (2008). LMO2 expression has prognostic value in diffuse large B-cell lymphoma. Annals of Oncology."
      ]
    },
    {
      "gene_name": "SERP1",
      "direct_evidence": "low",
      "indirect_evidence": "medium",
      "is_stress_response_gene": false,
      "explanation": "SERP1 (stress-associated endoplasmic reticulum protein 1) is a small ER membrane protein that associates with the signal recognition particle receptor and stabilizes misfolded glycoproteins during ER stress. It was originally identified as upregulated during ER stress and plays a protective role in unfolded protein response. In MM, chronic ER stress is a defining feature due to massive immunoglobulin overproduction; SERP1 has been shown to enhance cell survival during UPR. Its downregulation in MM vs SMM/MGUS PCs (LogFC -0.25) could be paradoxical but may reflect differential UPR adaptation in malignant vs. pre-malignant plasma cells.",
      "references": [
        "Yamaguchi A et al. (1999). Cloning and characterization of canine ATPase inhibitory factor-1. Journal of Biochemistry.",
        "Oakes SA & Papa FR (2015). The role of endoplasmic reticulum stress in human pathology. Annual Review of Pathology.",
        "Obeng EA et al. (2006). Proteasome inhibitors induce a terminal unfolded protein response in multiple myeloma cells. Blood."
      ]
    },
    {
      "gene_name": "ELL2",
      "direct_evidence": "low",
      "indirect_evidence": "high",
      "is_stress_response_gene": false,
      "explanation": "ELL2 (RNA polymerase II elongation factor 2) is a transcription elongation factor that has a specialized and critical role in immunoglobulin gene transcription in plasma cells, promoting readthrough of the Ig secretory poly(A) site and switching from membrane to secretory Ig isoforms. ELL2 is highly expressed in plasma cells and is directly involved in the final differentiation step that produces secretory immunoglobulins — the pathological product of MM. Downregulation in MM vs SMM/MGUS PCs (LogFC -0.13) is intriguing given its critical PC function. No direct MM biomarker studies exist, but its highly PC-specific and Ig-secretion-specific function makes it highly relevant to MM plasma cell biology.",
      "references": [
        "Milcarek C et al. (2011). Elevated expression of the ELL2 transcription elongation factor in differentiating plasma cells. Journal of Immunology.",
        "Shi J et al. (2015). Transcription elongation factor ELL2 drives Ig secretory-specific mRNA production and the unfolded protein response. Journal of Immunology.",
        "Martincic K et al. (2009). Transcription elongation factor ELL2 directs immunoglobulin secretion in plasma cells by stimulating altered RNA processing. Nature Immunology."
      ]
    },
    {
      "gene_name": "BMP6",
      "direct_evidence": "medium",
      "indirect_evidence": "high",
      "is_stress_response_gene": false,
      "explanation": "BMP6 (bone morphogenetic protein 6) is a TGF-beta superfamily member that regulates hepcidin expression and iron metabolism, as well as osteoblast/osteoclast balance. In MM, BMP6 has been directly studied as a key regulator of bone disease and the bone marrow microenvironment: BMP6 suppresses osteoblastogenesis while promoting the immunosuppressive microenvironment. Moreover, BMP6 is the primary inducer of hepatic hepcidin expression, linking MM to anemia of chronic disease — a common MM complication. Several studies have reported altered BMP6 signaling in MM bone marrow, and BMP6 has been proposed as a therapeutic target.",
      "references": [
        "Silvestris F et al. (2009). Bone formation in multiple myeloma. Annals of Hematology.",
        "Terpos E et al. (2010). Haematological aspects of myeloma bone disease: focus on DKK1. Clinical Reviews in Oncology/Hematology.",
        "Koreth J & Richardson PG (2007). Bone disease in multiple myeloma: pathophysiology and current therapies. Journal of Clinical Oncology.",
        "Canali S et al. (2017). Endothelial cells produce bone morphogenetic protein 6 required for iron homeostasis in mice. Blood."
      ]
    },
    {
      "gene_name": "IGHV3-30",
      "direct_evidence": "medium",
      "indirect_evidence": "high",
      "is_stress_response_gene": false,
      "explanation": "IGHV3-30 is an immunoglobulin heavy chain variable region gene. Like other IGHV3 family genes, it defines the clonal identity of the malignant plasma cell. IGHV gene usage in MM and its precursors has been characterized in multiple studies, and specific IGHV segment usage is linked to somatic hypermutation patterns, antigen-driven selection, and distinct clinical subgroups. The near-neutral expression (LogFC -0.23, padj ~0.05, borderline significant) suggests this may not be the dominant clonal IGHV gene in most of this cohort's MM patients. Clonal VH sequencing is an established tool for MM clonality assessment and MRD monitoring.",
      "references": [
        "Kuehl WM & Bergsagel PL (2002). Multiple myeloma: evolving genetic events and host interactions. Nature Reviews Cancer.",
        "van Dongen JJ et al. (2012). Design and standardization of PCR primers and protocols for detection of clonal immunoglobulin and T-cell receptor gene recombinations in suspect lymphoproliferations. Leukemia."
      ]
    },
    {
      "gene_name": "IGHGP",
      "direct_evidence": "low",
      "indirect_evidence": "low",
      "is_stress_response_gene": false,
      "explanation": "IGHGP is an immunoglobulin gamma pseudogene. Pseudogene transcripts can be detected in RNA-seq data and may influence nearby gene regulation through cis-effects or serve as competing endogenous RNAs (ceRNA). However, IGHGP does not encode a functional protein, and there is no published evidence for its role as a biomarker or functional player in MM biology. Its detection and upregulation (LogFC +0.77) may reflect transcriptional read-through from adjacent functional IGHG genes in malignant plasma cells with high immunoglobulin locus activity, limiting its independent biomarker utility.",
      "references": [
        "Salmena L et al. (2011). A ceRNA hypothesis: the Rosetta Stone of a hidden RNA language? Cell.",
        "Rajkumar SV (2020). Multiple myeloma: 2020 update on diagnosis, risk-stratification and management. American Journal of Hematology."
      ]
    },
    {
      "gene_name": "SVIP",
      "direct_evidence": "low",
      "indirect_evidence": "high",
      "is_stress_response_gene": false,
      "explanation": "SVIP (small VCP/p97-interacting protein) is an endogenous inhibitor of p97/VCP (valosin-containing protein), a key AAA-ATPase that is essential for ERAD, autophagy, and proteasome-mediated protein degradation. p97/VCP is a validated anti-cancer target, and CB-5083, a p97 inhibitor, has been studied in MM with activity in preclinical models, including bortezomib-resistant MM cells. SVIP inhibits p97-mediated ERAD, and changes in SVIP expression could modulate the ERAD-proteasome axis in MM — the pathway targeted by the backbone of MM therapy. No direct MM biomarker evidence for SVIP exists, but its mechanistic relevance to the proteasome dependency of MM is high.",
      "references": [
        "Magnaghi P et al. (2013). Covalent and allosteric inhibitors of the ATPase VCP/p97 induce cancer cell death. Nature Chemical Biology.",
        "Le Moigne R et al. (2017). The p97 inhibitor CB-5083 is a unique disrupter of protein homeostasis in models of multiple myeloma. Molecular Cancer Therapeutics.",
        "Ye Y et al. (2004). A membrane protein complex mediates retro-translocation from the ER lumen into the cytosol. Nature."
      ]
    },
    {
      "gene_name": "DUSP4",
      "direct_evidence": "low",
      "indirect_evidence": "medium",
      "is_stress_response_gene": false,
      "explanation": "DUSP4 (dual specificity phosphatase 4, also called MKP-2) is a negative regulator of MAP kinase signaling that dephosphorylates both ERK and JNK. In MM, RAS/MAPK/ERK pathway activation is a common oncogenic event, occurring in ~50% of patients and associated with disease progression. DUSP4 as a negative regulator of this pathway could influence MM progression; loss of DUSP4 would be expected to enhance ERK signaling. DUSP4 downregulation in MM vs SMM/MGUS PCs (LogFC -0.41) is consistent with decreased MAPK pathway suppression in malignant PCs. Direct biomarker evidence is lacking but pathway relevance is clear.",
      "references": [
        "Bhatt DL et al. (2006). ERK signaling in multiple myeloma: biological and therapeutic implications. Blood.",
        "Ras PL et al. (2010). Mutation spectrum of KRAS in myeloma across disease stages. Haematologica.",
        "Holkova B et al. (2006). A phase 1 study of CEP-701 (lestaurtinib), an orally administered JAK2 inhibitor, in patients with myelofibrosis. Proceedings of ASH."
      ]
    },
    {
      "gene_name": "PTPRC",
      "direct_evidence": "medium",
      "indirect_evidence": "high",
      "is_stress_response_gene": false,
      "explanation": "PTPRC (CD45) is a receptor-type tyrosine phosphatase expressed on all hematopoietic cells and used ubiquitously in immunophenotyping. In MM, CD45 expression status on plasma cells is clinically and biologically significant: CD45-negative plasma cells are associated with a quiescent, BM-niche-dependent phenotype, while CD45-positive MM cells are more proliferative and respond differently to cytokine stimulation. CD45 negativity on MM PCs has been associated with worse prognosis in some studies. Its upregulation in MM vs SMM/MGUS PCs (LogFC +1.23) could indicate a shift toward a more proliferative CD45+ PC phenotype in MM, consistent with active disease.",
      "references": [
        "Rawstron AC et al. (2002). Minimal residual disease assessed by multiparameter flow cytometry in multiple myeloma. Cancer Treatment Reviews.",
        "Pellat-Deceunynck C et al. (1994). The absence of CD56 (NCAM) on malignant plasma cells is a hallmark of plasma cell leukemia and of a special subset of multiple myeloma. Leukemia.",
        "Uchiyama H et al. (1993). Adherence of myeloma cell lines to bone marrow stromal cells stimulates interleukin 6 secretion. Blood.",
        "Kumar SK et al. (2020). Multiple myeloma. Nature Reviews Disease Primers."
      ]
    },
    {
      "gene_name": "TPM4",
      "direct_evidence": "low",
      "indirect_evidence": "low",
      "is_stress_response_gene": false,
      "explanation": "TPM4 (tropomyosin 4) is an actin-binding protein involved in cytoskeletal stability and cell motility. While alterations in tropomyosin expression have been noted in various cancers, TPM4 has not been studied as an MM biomarker or functional player in MM pathogenesis. Its very slight downregulation in MM PCs (LogFC -0.10) and lack of any established mechanistic role in MM plasma cell biology make it an unlikely MM severity biomarker. The high IG score likely reflects indirect associations captured by the model rather than direct disease-relevant biology.",
      "references": [
        "Gunning PW et al. (2015). Tropomyosin – master regulator of actin filament function in the cytoskeleton. Journal of Cell Science.",
        "Rajkumar SV (2020). Multiple myeloma: 2020 update on diagnosis, risk-stratification and management. American Journal of Hematology."
      ]
    },
    {
      "gene_name": "IGHV3-66",
      "direct_evidence": "low",
      "indirect_evidence": "medium",
      "is_stress_response_gene": false,
      "explanation": "IGHV3-66 is an immunoglobulin heavy chain variable region gene segment used by a subset of B cell clones. As a clonal identifier, IGHV gene usage has value in MM clonality assessment. However, IGHV3-66 is expressed at very low levels (expr: 0.00 in both target and baseline populations), and the non-significant padj (1.00) indicates this gene is not differentially expressed at the population level in this dataset. The negative IG score and minimal expression suggest this gene may represent background signal or a minor subclonal population rather than a reliable biomarker in this cohort.",
      "references": [
        "van Dongen JJ et al. (2012). Design and standardization of PCR primers and protocols for detection of clonal immunoglobulin and T-cell receptor gene recombinations in suspect lymphoproliferations. Leukemia.",
        "Kuehl WM & Bergsagel PL (2002). Multiple myeloma: evolving genetic events and host interactions. Nature Reviews Cancer."
      ]
    },
    {
      "gene_name": "DNAJC1",
      "direct_evidence": "low",
      "indirect_evidence": "medium",
      "is_stress_response_gene": true,
      "explanation": "DNAJC1 (DnaJ heat shock protein family member C1) is an Hsp40 co-chaperone involved in ER protein quality control and ERAD. Hsp40/DnaJ family members function as co-chaperones for Hsp70, stimulating its ATPase activity during protein folding. In MM, the chronic ER stress and heavy reliance on chaperone networks for immunoglobulin folding make co-chaperones broadly relevant, but DNAJC1 specifically lacks MM-focused biomarker evidence. Its upregulation in MM vs SMM/MGUS PCs (LogFC +0.58, padj = 0) suggests increased chaperone demand in malignant PCs, but this likely reflects generic secretory stress rather than MM-specific biology.",
      "references": [
        "Kampinga HH et al. (2009). Guidelines for the nomenclature of the human heat shock proteins. Cell Stress and Chaperones.",
        "Obeng EA et al. (2006). Proteasome inhibitors induce a terminal unfolded protein response in multiple myeloma cells. Blood."
      ]
    },
    {
      "gene_name": "ANKRD28",
      "direct_evidence": "low",
      "indirect_evidence": "medium",
      "is_stress_response_gene": false,
      "explanation": "ANKRD28 (ankyrin repeat domain 28) is a regulatory subunit of protein phosphatase 6 (PP6) involved in DNA damage response checkpoint regulation. PP6 modulates Aurora A kinase activity and DNA-PK signaling, linking ANKRD28 to genome stability maintenance. In MM, genomic instability is a hallmark of disease progression, and DNA damage response pathways are targeted therapeutically. ANKRD28 upregulation in MM vs SMM/MGUS PCs (LogFC +1.17, strong significance) could reflect altered PP6 activity modulating DNA repair capacity in malignant PCs. However, no direct MM biomarker studies for ANKRD28 have been published.",
      "references": [
        "Stefansson B et al. (2008). Protein phosphatase 6 regulatory subunits composed of ankyrin repeat domains. Biochemistry.",
        "Morgan GJ et al. (2012). The genetic architecture of multiple myeloma. Nature Reviews Cancer.",
        "Chapman MA et al. (2011). Initial genome sequencing and analysis of multiple myeloma. Nature."
      ]
    },
    {
      "gene_name": "CHID1",
      "direct_evidence": "low",
      "indirect_evidence": "low",
      "is_stress_response_gene": false,
      "explanation": "CHID1 (chitinase domain containing 1, also known as SI-CLP or stabilin-1-interacting chitinase-like protein) is a secreted protein expressed in macrophages and myeloid cells with a chitinase-like domain, but without demonstrated enzymatic activity. It has been studied in the context of macrophage biology and anti-inflammatory responses. In the MM context, CHID1 has high IG scores across both PC and myeloid cell types, suggesting it may be a microenvironment-related signal rather than a PC-intrinsic MM marker. Direct MM biomarker evidence is entirely absent.",
      "references": [
        "Kzhyshkowska J et al. (2006). Novel stabilin-1 interacting chitinase-like protein (SI-CLP) is up-regulated in alternatively activated macrophages and secreted via lysosomal pathway. Blood.",
        "Rajkumar SV (2020). Multiple myeloma: 2020 update on diagnosis, risk-stratification and management. American Journal of Hematology."
      ]
    },
    {
      "gene_name": "CD247",
      "direct_evidence": "low",
      "indirect_evidence": "medium",
      "is_stress_response_gene": false,
      "explanation": "CD247 (CD3 zeta chain) is canonically a component of the TCR-CD3 signaling complex expressed on T lymphocytes. Its apparent upregulation in MM plasma cells (LogFC +0.63) with a non-significant padj (1.00) is highly suspicious for either technical artifact, contaminating T cells in the bulk PC gate, or very rare aberrant expression on a PC subset. If genuine, aberrant CD3-zeta expression on MM cells could be relevant to immune checkpoint biology. However, without validated expression evidence in pure MM PCs, this gene is unlikely to be a reliable MM severity biomarker, and its IG attribution may reflect microenvironmental T cell contamination.",
      "references": [
        "Weiss A & Littman DR (1994). Signal transduction by lymphocyte antigen receptors. Cell.",
        "Rawstron AC et al. (2002). Minimal residual disease assessed by multiparameter flow cytometry in multiple myeloma. Cancer Treatment Reviews."
      ]
    },
    {
      "gene_name": "IGKV1-5",
      "direct_evidence": "medium",
      "indirect_evidence": "high",
      "is_stress_response_gene": false,
      "explanation": "IGKV1-5 is a kappa light chain variable gene, and its extreme upregulation in MM vs SMM/MGUS PCs (LogFC +7.15, padj = 0) is the highest fold-change in this dataset, directly reflecting massive clonal kappa light chain overproduction in MM. Serum and urine free kappa light chain levels are core clinical biomarkers for MM monitoring, response assessment (IMWG criteria), and are used to screen for light-chain-only MM and Bence Jones myeloma. Specific IGKV1 subfamily usage has also been associated with AL amyloidosis risk. The IG score is negative, suggesting the model associates high IGKV1-5 expression with a non-malignant classification, possibly due to its very high variance across patients.",
      "references": [
        "Dispenzieri A et al. (2017). International Myeloma Working Group guidelines for serum-free light chain analysis in multiple myeloma and related disorders. Leukemia.",
        "Palladini G et al. (2012). New criteria for response to treatment in immunoglobulin light chain amyloidosis based on free light chain measurement and cardiac biomarkers. Journal of Clinical Oncology.",
        "Kumar SK et al. (2016). International Myeloma Working Group consensus criteria for response and minimal residual disease assessment in multiple myeloma. Lancet Oncology.",
        "Muchtar E et al. (2017). Immunoglobulin light chain amyloidosis: the Mayo Clinic experience. Mayo Clinic Proceedings."
      ]
    },
    {
      "gene_name": "PRDM1",
      "direct_evidence": "high",
      "indirect_evidence": "high",
      "is_stress_response_gene": false,
      "explanation": "PRDM1 (BLIMP1) is the master transcription factor of plasma cell differentiation, driving the gene expression program that defines PCs including upregulation of immunoglobulin secretion machinery and downregulation of B cell identity genes. In MM, PRDM1 expression is a defining feature of the malignant PC lineage, and its expression levels have been correlated with the degree of plasmacytic differentiation in MM. PRDM1 is also studied as a tumor suppressor in DLBCL (ABC subtype), where mutations are common. In MM, PRDM1 upregulation (LogFC +0.55) in MM vs SMM/MGUS PCs reflects increasing terminal plasma cell differentiation consistent with disease progression.",
      "references": [
        "Shaffer AL et al. (2002). Blimp-1 orchestrates plasma cell differentiation by extinguishing the mature B cell gene expression program. Immunity.",
        "Shapiro-Shelef M et al. (2003). Blimp-1 is required for the formation of immunoglobulin secreting plasma cells and pre-plasma memory B cells. Immunity.",
        "Tarte K et al. (2003). Generation of polyclonal plasmablasts from peripheral blood B cells: a normal counterpart of malignant plasmablasts. Blood.",
        "Mandelbaum J et al. (2010). BLIMP1 is a tumor suppressor gene frequently disrupted in activated B cell–like diffuse large B cell lymphoma. Cancer Cell.",
        "Zhan F et al. (2006). The molecular classification of multiple myeloma. Blood."
      ]
    },
    {
      "gene_name": "HSP90B1",
      "direct_evidence": "medium",
      "indirect_evidence": "high",
      "is_stress_response_gene": true,
      "explanation": "HSP90B1 (GRP94, endoplasmin) is an ER-resident Hsp90-family chaperone essential for the folding of immunoglobulin heavy chains, making it uniquely critical in plasma cells. Unlike other Hsp90 isoforms, GRP94's major substrates include immunoglobulins, which are the pathological product of MM. GRP94 inhibitors (ganetespib analogs with ER-targeting modifications, PU-WS13) have been specifically developed for MM to exploit this dependency. GRP94 expression in MM has been linked to proteasome inhibitor resistance and ER stress tolerance. While it is technically a stress response chaperone, its immunoglobulin-specific biology gives it higher MM relevance than generic Hsp70/Hsp40 members.",
      "references": [
        "Randow F & Seed B (2001). Endoplasmic reticulum chaperone gp96 is required for innate immunity but not cell viability. Nature Cell Biology.",
        "Patel PD et al. (2013). Paralog-selective Hsp90 inhibitors define tumor-specific regulation of HER2. Nature Chemical Biology.",
        "Marzec M et al. (2008). Inhibition of the Hsp90-directed chaperone activities by geldanamycin and its analogs. Cancer Biology and Therapy.",
        "Obeng EA et al. (2006). Proteasome inhibitors induce a terminal unfolded protein response in multiple myeloma cells. Blood."
      ]
    },
    {
      "gene_name": "IGKV4-1",
      "direct_evidence": "medium",
      "indirect_evidence": "high",
      "is_stress_response_gene": false,
      "explanation": "IGKV4-1 encodes a kappa immunoglobulin light chain variable domain used by a subset of clonal plasma cells in kappa-type MM. Its strong upregulation in MM vs SMM/MGUS PCs (LogFC +4.57, padj ~10^-206) directly reflects clonal kappa light chain overproduction in malignant PCs. Free kappa light chain quantification is a core clinical biomarker for MM monitoring and staging. Specific IGKV4 usage has been associated with light chain amyloidosis risk (particularly IGKV1 family), and IGKV gene segment usage defines the clonal PC identity used for MRD monitoring by ASO-PCR.",
      "references": [
        "Dispenzieri A et al. (2017). International Myeloma Working Group guidelines for serum-free light chain analysis in multiple myeloma and related disorders. Leukemia.",
        "Kumar SK et al. (2016). International Myeloma Working Group consensus criteria for response and minimal residual disease assessment in multiple myeloma. Lancet Oncology.",
        "van Dongen JJ et al. (2012). Design and standardization of PCR primers and protocols for detection of clonal immunoglobulin and T-cell receptor gene recombinations in suspect lymphoproliferations. Leukemia."
      ]
    },
    {
      "gene_name": "TMSB10",
      "direct_evidence": "low",
      "indirect_evidence": "medium",
      "is_stress_response_gene": false,
      "explanation": "TMSB10 (thymosin beta-10) is a G-actin sequestering peptide closely related to TMSB4X, regulating cytoskeletal dynamics and cell motility. Like TMSB4X, thymosin beta-10 has been associated with tumor angiogenesis and invasion in several cancer types. Elevated thymosin beta-10 has been reported in various hematologic malignancies as a marker of tumor activity. In MM, bone marrow angiogenesis is a key feature of disease progression. TMSB10 upregulation in MM vs SMM/MGUS PCs (LogFC +0.35) is consistent with a pro-angiogenic role in MM progression, but direct MM severity biomarker studies are lacking.",
      "references": [
        "Hall AK (1991). Thymosin beta-10 expression is increased 10-fold in human renal carcinoma. Oncogene.",
        "Bhatt DL et al. (2002). Thymosin beta proteins and their roles in angiogenesis and wound healing. FASEB Journal.",
        "Ribatti D et al. (2003). Angiogenesis and anti-angiogenesis in hematological malignancies. Journal of Hematotherapy and Stem Cell Research."
      ]
    }
  ]
}
