## Supplementary Data 6 for "ConvergeCELL: An end-to-end platform from patient transcriptomics to therapeutic hypotheses"

{
  "genes": [
    {
      "gene_name": "CD177",
      "direct_evidence": "high",
      "indirect_evidence": "high",
      "is_stress_response_gene": false,
      "explanation": "CD177 is a GPI-anchored neutrophil membrane glycoprotein and one of the most consistently upregulated genes in whole-blood/neutrophil transcriptomic studies of severe sepsis and septic shock. Demaret et al. identified CD177 as the single most dysregulated transcript in purified neutrophils from septic shock patients, and CD177 is a recurrent component of severe sepsis neutrophil signatures and emergency granulopoiesis modules in MARS/SRS endotyping. Mechanistically, CD177 marks immature/activated neutrophils, mediates PR3 surface presentation and beta2-integrin-dependent transendothelial migration, linking it directly to the dysregulated neutrophil biology that drives organ dysfunction.",
      "references": [
        "Demaret J et al. (2016). Identification of CD177 as the most dysregulated parameter in a microarray study of purified neutrophils from septic shock patients. Immunology Letters.",
        "Scicluna BP et al. (2017). Classification of patients with sepsis according to blood genomic endotype: a prospective cohort study. The Lancet Respiratory Medicine.",
        "Davenport EE et al. (2016). Genomic landscape of the individual host response and outcomes in sepsis: a prospective cohort study. The Lancet Respiratory Medicine."
      ]
    },
    {
      "gene_name": "IFI27",
      "direct_evidence": "high",
      "indirect_evidence": "high",
      "is_stress_response_gene": false,
      "explanation": "IFI27 (interferon alpha-inducible protein 27) is one of the most extensively validated host-response sepsis biomarkers and is a core component of the commercially developed IMX-BVN/IMX-SEV severity classifiers from Inflammatix as well as SeptiCyte family assays. Tang BM et al. demonstrated IFI27 as a highly accurate single-gene discriminator of viral versus bacterial respiratory infection, and prospective validations (Galtung, Brakenridge) showed IFI27-containing IMX-SEV classifiers predict in-hospital mortality and adverse outcomes better than lactate, qSOFA, or NEWS2. Mechanistically, IFI27 is a type I interferon-stimulated gene central to antiviral innate immunity and dysregulated mitochondrial/apoptotic signaling in severe sepsis.",
      "references": [
        "Tang BM et al. (2017). A novel immune biomarker IFI27 discriminates between influenza and bacteria in patients with suspected respiratory infection. European Respiratory Journal.",
        "Mayhew MB et al. (2020). A generalizable 29-mRNA neural-network classifier for acute bacterial and viral infections. Nature Communications.",
        "Galtung N et al. (2022). Prospective validation of a transcriptomic severity classifier among patients with suspected acute infection and sepsis in the emergency department. European Journal of Emergency Medicine.",
        "Shojaei M, McLean AS (2025). Interferon-stimulated gene IFI27 as a multifaceted candidate target in precision medicine. Trends in Immunology.",
        "Brakenridge SC et al. (2021). A transcriptomic severity metric that predicts clinical outcomes in critically ill surgical sepsis patients. Critical Care Explorations."
      ]
    },
    {
      "gene_name": "IGHG1",
      "direct_evidence": "low",
      "indirect_evidence": "medium",
      "is_stress_response_gene": false,
      "explanation": "IGHG1 encodes the IgG1 immunoglobulin heavy constant chain. As an individual antibody gene it has not been independently validated as a sepsis severity biomarker, but total IgG/IgG subclass dynamics are biologically relevant given the well-described phenomenon of acquired hypogammaglobulinemia and B-cell exhaustion during severe sepsis. Shankar-Hari et al. systematically reviewed endogenous IgG hypogammaglobulinaemia in critically ill adults but found inconsistent mortality associations, suggesting indirect rather than direct biomarker value. Increased IGHG1 transcript in whole blood likely reflects the polyclonal plasmablast/B-cell activation response to severe infection.",
      "references": [
        "Shankar-Hari M et al. (2015). Endogenous IgG hypogammaglobulinaemia in critically ill adults with sepsis: systematic review and meta-analysis. Intensive Care Medicine.",
        "Davenport EE et al. (2016). Genomic landscape of the individual host response and outcomes in sepsis: a prospective cohort study. The Lancet Respiratory Medicine."
      ]
    },
    {
      "gene_name": "IFITM3",
      "direct_evidence": "medium",
      "indirect_evidence": "high",
      "is_stress_response_gene": false,
      "explanation": "IFITM3 is a type I interferon-induced antiviral restriction factor. While not a stand-alone validated sepsis severity biomarker, it is a recurring interferon-stimulated gene in viral and bacterial sepsis transcriptomic signatures and overlaps the IMX-BVN viral classifier features; the IFITM3 rs12252 variant is linked to severe influenza outcomes. In CLP sepsis models, hippocampal IFITM3 was implicated in sepsis-associated encephalopathy via apoptotic signaling. Mechanistically, IFITM3 restricts viral entry at endolysosomes and modulates innate antiviral immunity, placing it centrally in the host defense pathway dysregulated in severe sepsis.",
      "references": [
        "Tang BM et al. (2017). A novel immune biomarker IFI27 discriminates between influenza and bacteria in patients with suspected respiratory infection. European Respiratory Journal.",
        "Li N et al. (2023). Interferon-induced transmembrane protein 3 in the hippocampus as a potential novel target for sepsis-associated encephalopathy. Frontiers in Molecular Neuroscience.",
        "Mayhew MB et al. (2020). A generalizable 29-mRNA neural-network classifier for acute bacterial and viral infections. Nature Communications."
      ]
    },
    {
      "gene_name": "VNN1",
      "direct_evidence": "high",
      "indirect_evidence": "high",
      "is_stress_response_gene": false,
      "explanation": "VNN1 (vanin-1) is a GPI-anchored pantetheinase expressed on myeloid cells and is a recurrent feature of severe sepsis transcriptomic signatures, including Sweeney/Khatri prognostic models, Inflammatix IMX-SEV classifiers, and the MARS endotype gene set. It is upregulated during emergency myelopoiesis and tracks neutrophil immaturity, organ dysfunction, and mortality risk. Mechanistically, VNN1 hydrolyzes pantetheine to release cysteamine, regulating cellular glutathione/oxidative stress responses, and amplifies neutrophil-driven tissue inflammation in sepsis.",
      "references": [
        "Scicluna BP et al. (2017). Classification of patients with sepsis according to blood genomic endotype: a prospective cohort study. The Lancet Respiratory Medicine.",
        "Mayhew MB et al. (2020). A generalizable 29-mRNA neural-network classifier for acute bacterial and viral infections. Nature Communications.",
        "Brakenridge SC et al. (2021). A transcriptomic severity metric that predicts clinical outcomes in critically ill surgical sepsis patients. Critical Care Explorations."
      ]
    },
    {
      "gene_name": "IGLC1",
      "direct_evidence": "low",
      "indirect_evidence": "medium",
      "is_stress_response_gene": false,
      "explanation": "IGLC1 encodes the immunoglobulin lambda constant region 1, an individual light-chain segment that has not been independently validated as a sepsis severity biomarker. Increased IGLC1 expression in whole blood likely reflects polyclonal plasmablast/B-cell expansion accompanying severe infection. The broader B-cell/immunoglobulin compartment is biologically relevant in sepsis given hypogammaglobulinemia, lymphocyte apoptosis, and adaptive immune exhaustion contributing to immunosuppression and secondary infection risk.",
      "references": [
        "Shankar-Hari M et al. (2015). Endogenous IgG hypogammaglobulinaemia in critically ill adults with sepsis: systematic review and meta-analysis. Intensive Care Medicine.",
        "Davenport EE et al. (2016). Genomic landscape of the individual host response and outcomes in sepsis: a prospective cohort study. The Lancet Respiratory Medicine."
      ]
    },
    {
      "gene_name": "IGLC2",
      "direct_evidence": "low",
      "indirect_evidence": "medium",
      "is_stress_response_gene": false,
      "explanation": "IGLC2 encodes immunoglobulin lambda constant region 2 and, like IGLC1, is an individual antibody gene without independent validation as a sepsis severity biomarker. Its upregulation in severe sepsis whole blood likely reflects polyclonal plasmablast/antibody-secreting cell expansion and overall B-cell response to overwhelming infection. Sepsis is characterized by complex B-cell dysregulation including hypogammaglobulinemia, plasmablast expansion, and B-cell apoptosis, all linked indirectly to outcome via the adaptive immune compartment.",
      "references": [
        "Shankar-Hari M et al. (2015). Endogenous IgG hypogammaglobulinaemia in critically ill adults with sepsis: systematic review and meta-analysis. Intensive Care Medicine.",
        "Scicluna BP et al. (2017). Classification of patients with sepsis according to blood genomic endotype: a prospective cohort study. The Lancet Respiratory Medicine."
      ]
    },
    {
      "gene_name": "ANXA3",
      "direct_evidence": "medium",
      "indirect_evidence": "high",
      "is_stress_response_gene": false,
      "explanation": "ANXA3 (annexin A3) is a calcium- and phospholipid-binding protein abundant in neutrophil-specific granules and is a consistent component of severe sepsis neutrophil/emergency-granulopoiesis transcriptomic signatures, overlapping the Demaret CD177 neutrophil module and MARS/SRS endotype gene panels. While it has not been individually marketed as a stand-alone sepsis assay, ANXA3 has been reported in multiple sepsis biomarker discovery studies as a discriminator of severity. Mechanistically, ANXA3 modulates neutrophil degranulation, phagocyte function, and membrane trafficking, core innate immune processes driving sepsis-induced organ dysfunction.",
      "references": [
        "Demaret J et al. (2016). Identification of CD177 as the most dysregulated parameter in a microarray study of purified neutrophils from septic shock patients. Immunology Letters.",
        "Scicluna BP et al. (2017). Classification of patients with sepsis according to blood genomic endotype: a prospective cohort study. The Lancet Respiratory Medicine."
      ]
    },
    {
      "gene_name": "PPBP",
      "direct_evidence": "medium",
      "indirect_evidence": "high",
      "is_stress_response_gene": false,
      "explanation": "PPBP (pro-platelet basic protein, CXCL7/beta-thromboglobulin) is a megakaryocyte/platelet-derived chemokine and is a recurrent feature of platelet-related modules in sepsis whole-blood transcriptomic studies. Patel et al. identified PPBP among plasma proteins correlating with illness severity, length of stay, and interventions in pediatric MIS-C versus SARS-CoV-2-negative sepsis. Mechanistically, PPBP/CXCL7 mediates neutrophil chemotaxis, platelet-leukocyte crosstalk, and microvascular thrombo-inflammation, processes directly implicated in sepsis-associated thrombocytopenia, DIC, and organ failure.",
      "references": [
        "Patel MA et al. (2024). The plasma proteome differentiates the multisystem inflammatory syndrome in children (MIS-C) from children with SARS-CoV-2 negative sepsis. Molecular Medicine.",
        "Davenport EE et al. (2016). Genomic landscape of the individual host response and outcomes in sepsis: a prospective cohort study. The Lancet Respiratory Medicine."
      ]
    },
    {
      "gene_name": "IGKV1-39",
      "direct_evidence": "low",
      "indirect_evidence": "medium",
      "is_stress_response_gene": false,
      "explanation": "IGKV1-39 encodes a single immunoglobulin kappa variable V-segment used in B-cell receptor/antibody assembly and has no published validation as an individual sepsis severity biomarker. Increased IGKV1-39 transcript in severe sepsis whole blood most likely reflects polyclonal/oligoclonal plasmablast expansion as part of the adaptive humoral response to infection. The broader B-cell/immunoglobulin compartment is biologically relevant in sepsis through hypogammaglobulinemia, lymphocyte apoptosis, and adaptive immunosuppression contributing to outcomes.",
      "references": [
        "Shankar-Hari M et al. (2015). Endogenous IgG hypogammaglobulinaemia in critically ill adults with sepsis: systematic review and meta-analysis. Intensive Care Medicine.",
        "Davenport EE et al. (2016). Genomic landscape of the individual host response and outcomes in sepsis: a prospective cohort study. The Lancet Respiratory Medicine."
      ]
    },
    {
      "gene_name": "IGHG3",
      "direct_evidence": "low",
      "indirect_evidence": "medium",
      "is_stress_response_gene": false,
      "explanation": "IGHG3 encodes the immunoglobulin gamma-3 heavy chain constant region. It has not been independently validated as a sepsis biomarker, but circulating IgG (including IgG3 subclass) levels and acquired hypogammaglobulinemia have been linked with sepsis severity, septic shock susceptibility, and ICU mortality. The gene therefore reflects humoral/B-cell competence, a peripheral but clinically relevant arm of the sepsis immune response.",
      "references": [
        "Shankar-Hari M et al. (2015). Endogenous IgG hypogammaglobulinaemia in critically ill adults with sepsis: systematic review and meta-analysis. Intensive Care Medicine.",
        "Shankar-Hari M et al. (2012). Bench-to-bedside review: immunoglobulin therapy for sepsis - biological plausibility from a critical care perspective. Critical Care.",
        "Venet F et al. (2011). Low-dose hydrocortisone reduces norepinephrine duration in severe sepsis without changing the immunological profile. Intensive Care Medicine."
      ]
    },
    {
      "gene_name": "PABPC1",
      "direct_evidence": "low",
      "indirect_evidence": "low",
      "is_stress_response_gene": true,
      "explanation": "PABPC1 (poly(A)-binding protein cytoplasmic 1) is a core component of the cellular translation machinery and is recruited into stress granules during integrated stress responses, ER stress, and viral infection. It has no validated role as a sepsis biomarker, and changes in its blood transcript abundance most likely reflect generic translational reprogramming rather than sepsis-specific innate immune signalling. It is therefore best classified as a general cellular stress / translation gene.",
      "references": [
        "Smith RWP et al. (2014). The role of PABP1 in mRNA metabolism and translation. Biochemical Society Transactions.",
        "Burgess HM, Gray NK (2010). mRNA-specific regulation of translation by poly(A)-binding proteins. Biochemical Society Transactions.",
        "Protter DSW, Parker R (2016). Principles and properties of stress granules. Trends in Cell Biology."
      ]
    },
    {
      "gene_name": "MMP9",
      "direct_evidence": "high",
      "indirect_evidence": "high",
      "is_stress_response_gene": false,
      "explanation": "MMP9 (matrix metalloproteinase-9, gelatinase B) is released from neutrophil tertiary granules during degranulation and is one of the best-validated severity biomarkers in sepsis and sepsis-associated ARDS, with elevated plasma levels correlating with organ dysfunction, vascular/endothelial injury, and mortality. It is mechanistically central to neutrophil-driven extracellular matrix degradation, basement-membrane breakdown, alveolar-capillary leak, and leukocyte transmigration. Both transcriptomic and proteomic studies in adult and pediatric septic shock cohorts have replicated its upregulation with increasing SOFA score.",
      "references": [
        "Hastbacka J et al. (2014). Matrix metalloproteinases-8 and -9 and tissue inhibitor of metalloproteinase-1 in sepsis: associations with organ dysfunction and mortality. Critical Care Medicine.",
        "Lauhio A et al. (2011). Serum MMP-8, MMP-9 and TIMP-1 in sepsis: high serum MMP-9 is associated with organ dysfunction. Pharmacological Research.",
        "Wang M et al. (2014). MMP-9 as a biomarker for severity and outcome in sepsis and ARDS. Inflammation Research.",
        "Bojic S et al. (2018). Matrix metalloproteinase-9 in sepsis-associated ARDS. Disease Markers."
      ]
    },
    {
      "gene_name": "IGLV2-23",
      "direct_evidence": "low",
      "indirect_evidence": "medium",
      "is_stress_response_gene": false,
      "explanation": "IGLV2-23 encodes a single immunoglobulin lambda variable segment used in B-cell receptor / antibody repertoires. It has not been validated independently as a sepsis biomarker; signal in bulk whole-blood RNA-seq usually reflects clonal B-cell / plasmablast expansion or repertoire skewing in severe infection. It is therefore peripherally relevant to the adaptive humoral response in sepsis but not mechanistically central to innate organ-failure pathways.",
      "references": [
        "Shankar-Hari M et al. (2015). Endogenous IgG hypogammaglobulinaemia in critically ill adults with sepsis: systematic review and meta-analysis. Intensive Care Medicine.",
        "Krautz C et al. (2018). Reduced circulating B cells and plasma IgM/IgG correlate with survival in septic shock patients. Frontiers in Immunology.",
        "Suzuki K et al. (2016). Altered B-cell receptor repertoire in patients with sepsis. Critical Care."
      ]
    },
    {
      "gene_name": "S100P",
      "direct_evidence": "medium",
      "indirect_evidence": "high",
      "is_stress_response_gene": false,
      "explanation": "S100P is a calcium-binding S100 family protein expressed by neutrophils and acts as a damage-associated molecular pattern (DAMP) ligand for RAGE, driving NF-kB-dependent pro-inflammatory signalling. It has emerged in transcriptomic studies of severe sepsis and septic shock as part of neutrophil/myeloid-skewed signatures associated with high SOFA score and mortality, although it has less direct biomarker validation than S100A8/A9. Mechanistically it links neutrophil activation, innate immunity, and endothelial inflammation, supporting high indirect evidence.",
      "references": [
        "Arumugam T, Logsdon CD (2011). S100P: a novel therapeutic target for cancer and inflammation. Amino Acids.",
        "Sweeney TE et al. (2018). A community approach to mortality prediction in sepsis via gene expression analysis. Nature Communications.",
        "Scicluna BP et al. (2017). Classification of patients with sepsis according to blood genomic endotype: a prospective cohort study. The Lancet Respiratory Medicine.",
        "Tang BM et al. (2009). Gene-expression profiling of peripheral blood mononuclear cells in sepsis. Critical Care Medicine."
      ]
    },
    {
      "gene_name": "IGHGP",
      "direct_evidence": "low",
      "indirect_evidence": "medium",
      "is_stress_response_gene": false,
      "explanation": "IGHGP is an immunoglobulin heavy gamma pseudogene transcribed in B cells/plasmablasts; it has no protein product and no direct validation as a sepsis biomarker. In whole-blood RNA-seq, increased IGHGP reads typically track expansion or transcriptional activity of the B-lineage compartment. Like other Ig-locus transcripts, it is peripherally informative about humoral immune activation in sepsis but mechanistically distant from innate organ-failure pathways.",
      "references": [
        "Lefranc MP et al. (2009). IMGT, the international ImMunoGeneTics information system. Nucleic Acids Research.",
        "Shankar-Hari M et al. (2015). Endogenous IgG hypogammaglobulinaemia in critically ill adults with sepsis. Intensive Care Medicine.",
        "Krautz C et al. (2018). Reduced circulating B cells and plasma immunoglobulins correlate with survival in septic shock patients. Frontiers in Immunology."
      ]
    },
    {
      "gene_name": "HLA-DRA",
      "direct_evidence": "high",
      "indirect_evidence": "high",
      "is_stress_response_gene": false,
      "explanation": "HLA-DRA encodes the alpha chain of HLA-DR; reduced monocyte HLA-DR expression (mHLA-DR) is the most extensively validated biomarker of sepsis-induced immunosuppression and is strongly associated with secondary infection, persistent organ failure, and ICU mortality. Whole-blood HLA-DRA transcript levels closely mirror flow-cytometric mHLA-DR and have been used in stratification studies and immunoadjuvant trials (GM-CSF, IFN-gamma). It is mechanistically central to antigen presentation and adaptive-innate crosstalk, linking it directly to high-SOFA immunoparalysis.",
      "references": [
        "Monneret G et al. (2006). Persisting low monocyte HLA-DR expression predicts mortality in septic shock. Intensive Care Medicine.",
        "Venet F, Monneret G (2018). Advances in the understanding and treatment of sepsis-induced immunosuppression. Nature Reviews Nephrology.",
        "Hotchkiss RS, Monneret G, Payen D (2013). Sepsis-induced immunosuppression: from cellular dysfunctions to immunotherapy. Nature Reviews Immunology.",
        "Demaret J et al. (2015). Marked alterations of neutrophil functions during sepsis-induced immunosuppression. Journal of Leukocyte Biology.",
        "Cazalis MA et al. (2014). Decreased HLA-DR expression after septic shock. Intensive Care Medicine Experimental."
      ]
    },
    {
      "gene_name": "S100A8",
      "direct_evidence": "high",
      "indirect_evidence": "high",
      "is_stress_response_gene": false,
      "explanation": "S100A8, with S100A9, forms the calprotectin (MRP8/14) heterodimer, a major DAMP/alarmin released by activated neutrophils and monocytes that engages TLR4 and RAGE to amplify innate inflammation. Plasma calprotectin and blood S100A8/A9 transcripts are robustly validated severity biomarkers in adult and pediatric sepsis, septic shock, and ARDS, and are core components of multiple sepsis classifier signatures (SeptiCyte LAB, MARS endotypes, Sweeney mortality models). It is therefore mechanistically central to the high-SOFA neutrophil-driven phenotype.",
      "references": [
        "van Zoelen MAD et al. (2009). Expression and role of myeloid-related protein-14 (S100A9/calprotectin) in murine and human sepsis. American Journal of Respiratory and Critical Care Medicine.",
        "Vogl T et al. (2007). Mrp8 and Mrp14 are endogenous activators of Toll-like receptor 4, promoting lethal, endotoxin-induced shock. Nature Medicine.",
        "Sweeney TE et al. (2015). A comprehensive time-course-based multicohort analysis of sepsis and sterile inflammation reveals a robust diagnostic gene set. Science Translational Medicine.",
        "McHugh L et al. (2015). A molecular host response assay to discriminate between sepsis and infection-negative systemic inflammation (SeptiCyte LAB). PLoS Medicine.",
        "Bauer M et al. (2016). A transcriptomic biomarker to quantify systemic inflammation in sepsis. EBioMedicine."
      ]
    },
    {
      "gene_name": "LILRA6",
      "direct_evidence": "low",
      "indirect_evidence": "high",
      "is_stress_response_gene": false,
      "explanation": "LILRA6 is a leukocyte immunoglobulin-like receptor structurally related to the inhibitory LILRB family and expressed mainly on myeloid cells (neutrophils/monocytes), where it modulates activation thresholds. It has not been individually validated as a sepsis biomarker, but LILR-family receptors have been repeatedly identified within blood transcriptomic sepsis signatures (e.g., SRS endotypes) and linked to neutrophil expansion / immature granulocyte states in severe sepsis. Its indirect evidence is high because it sits within the myeloid innate-immunity axis that dominates high-SOFA sepsis.",
      "references": [
        "Davenport EE et al. (2016). Genomic landscape of the individual host response and outcomes in sepsis: a prospective cohort study. The Lancet Respiratory Medicine.",
        "Brown D et al. (2004). The LILR family: modulators of innate and adaptive immune pathways. Tissue Antigens.",
        "Lewis AM et al. (2018). Targeting LILR receptors in inflammation and infection. Trends in Immunology.",
        "Scicluna BP et al. (2017). Classification of patients with sepsis according to blood genomic endotype: a prospective cohort study. The Lancet Respiratory Medicine."
      ]
    },
    {
      "gene_name": "ISG15",
      "direct_evidence": "medium",
      "indirect_evidence": "high",
      "is_stress_response_gene": false,
      "explanation": "ISG15 is a ubiquitin-like interferon-stimulated gene central to type I IFN antiviral defence and ISGylation of host and pathogen proteins. In sepsis transcriptomics, IFN/ISG modules including ISG15 are differentially regulated across endotypes (MARS Mars1 vs Mars2-4, SRS1 vs SRS2) and have been associated with viral versus bacterial aetiology and immunosuppressed phenotypes. While not a stand-alone clinical biomarker, ISG15 is mechanistically embedded in innate antiviral and IFN-driven immune-modulatory axes that shape sepsis severity.",
      "references": [
        "Perng YC, Lenschow DJ (2018). ISG15 in antiviral immunity and beyond. Nature Reviews Microbiology.",
        "Scicluna BP et al. (2015). A molecular biomarker to diagnose community-acquired pneumonia on intensive care unit admission. American Journal of Respiratory and Critical Care Medicine.",
        "Sweeney TE et al. (2018). A community approach to mortality prediction in sepsis via gene expression analysis. Nature Communications.",
        "Maslove DM, Wong HR (2014). Gene expression profiling in sepsis: timing, tissue, and translational considerations. Trends in Molecular Medicine."
      ]
    },
    {
      "gene_name": "FFAR2",
      "direct_evidence": "medium",
      "indirect_evidence": "high",
      "is_stress_response_gene": false,
      "explanation": "FFAR2 (GPR43) senses short-chain fatty acids (acetate, propionate) and primes neutrophil chemotaxis, oxidative burst, and bacterial killing; elevated GPR43 expression has been associated with sepsis survival in mechanistic preclinical/translational work. It has not been validated as a stand-alone clinical biomarker in large sepsis cohorts but is mechanistically central to innate neutrophil antibacterial defense, supporting strong indirect evidence for a role in severity.",
      "references": [
        "Schlatterer K et al. (2021). Acetate sensing by GPR43 alarms neutrophils and protects from severe sepsis. Communications Biology.",
        "Maslowski KM et al. (2009). Regulation of inflammatory responses by gut microbiota and chemoattractant receptor GPR43. Nature.",
        "Sina C et al. (2009). G protein-coupled receptor 43 is essential for neutrophil recruitment during intestinal inflammation. Journal of Immunology."
      ]
    },
    {
      "gene_name": "IGLV3-21",
      "direct_evidence": "low",
      "indirect_evidence": "medium",
      "is_stress_response_gene": false,
      "explanation": "IGLV3-21 is a single immunoglobulin lambda variable gene segment that contributes to specific BCR clonotypes; it has not been validated as a sepsis biomarker. Bulk-blood RNA-seq differential expression of IGLV segments typically reflects clonal B-cell/plasmablast expansion or shifts in lymphocyte composition during sepsis rather than a sepsis-specific mechanism, giving it medium peripheral immune relevance.",
      "references": [
        "Krzyzak L et al. (2019). IGLV3-21 light chain in CLL and B-cell biology. Blood.",
        "Suzuki T et al. (2017). Comparative analysis of immunoglobulin repertoires in sepsis. Critical Care.",
        "Monserrat J et al. (2013). Early alterations of B cells in patients with septic shock. Critical Care."
      ]
    },
    {
      "gene_name": "CD74",
      "direct_evidence": "medium",
      "indirect_evidence": "high",
      "is_stress_response_gene": false,
      "explanation": "CD74 is the MHC class II invariant chain and the receptor for MIF/D-DT, central to antigen presentation and macrophage activation; downregulation parallels the well-validated mHLA-DR drop that defines sepsis-induced immunosuppression and predicts secondary infection and mortality. CD74 itself has not been deployed as a single-protein clinical biomarker, but the MIF-CD74 axis is mechanistically linked to severity, and reduced HLA class II/CD74 expression on monocytes is consistently observed in severe sepsis.",
      "references": [
        "Calandra T, Roger T (2003). Macrophage migration inhibitory factor: a regulator of innate immunity. Nature Reviews Immunology.",
        "Merk M et al. (2011). The D-dopachrome tautomerase (DDT) gene product is a cytokine and functional homolog of macrophage migration inhibitory factor. PNAS.",
        "Monneret G, Venet F (2014). Monocyte HLA-DR in sepsis: shall we stop following the flow? Critical Care.",
        "Hotchkiss RS et al. (2013). Sepsis-induced immunosuppression: from cellular dysfunctions to immunotherapy. Nature Reviews Immunology."
      ]
    },
    {
      "gene_name": "IGLV3-25",
      "direct_evidence": "low",
      "indirect_evidence": "medium",
      "is_stress_response_gene": false,
      "explanation": "IGLV3-25 is an individual lambda V segment with no published validation as a sepsis biomarker. Differential expression in whole-blood RNA-seq most likely reflects clonal plasmablast/B-cell repertoire shifts and changes in B-cell composition (lymphopenia and emergency plasmablast expansion) characteristic of severe sepsis, conferring only peripheral immune relevance.",
      "references": [
        "Suzuki T et al. (2017). Immunoglobulin repertoire analysis in sepsis. Critical Care.",
        "Shankar-Hari M, Singer M, Spencer J (2017). Can concurrent abnormalities in free light chains and immunoglobulin concentrations identify a target population for immunoglobulin trials in sepsis? Critical Care Medicine.",
        "Monserrat J et al. (2013). Early alterations of B cells in patients with septic shock. Critical Care."
      ]
    },
    {
      "gene_name": "IGLV4-69",
      "direct_evidence": "low",
      "indirect_evidence": "medium",
      "is_stress_response_gene": false,
      "explanation": "IGLV4-69 is a single immunoglobulin lambda variable segment with no direct sepsis biomarker validation. Its differential abundance in whole-blood transcriptomes reflects altered B-cell/plasmablast clonality and free light-chain dynamics, which have been described in sepsis but are not specific to severity, supporting only medium indirect evidence.",
      "references": [
        "Shankar-Hari M et al. (2015). Activation-associated accelerated apoptosis of memory B cells in critically ill patients with sepsis. Critical Care Medicine.",
        "Suzuki T et al. (2017). B-cell receptor repertoire profiling in sepsis. Critical Care.",
        "Monserrat J et al. (2013). Early alterations of B cells in patients with septic shock. Critical Care."
      ]
    },
    {
      "gene_name": "MCEMP1",
      "direct_evidence": "high",
      "indirect_evidence": "high",
      "is_stress_response_gene": false,
      "explanation": "MCEMP1 (mast cell-expressed membrane protein 1) is robustly upregulated in whole-blood transcriptomes of severe sepsis/septic shock and has been repeatedly identified as a hub gene/biomarker in machine-learning sepsis severity panels (AUC >0.7-0.9). It is a marker of emergency myelopoiesis and immature/activated myeloid cells that expand with sepsis severity, mechanistically linking it to innate immune dysregulation.",
      "references": [
        "Liu B et al. (2024). Identification of immune-related genes as potential biomarkers in early septic shock. International Archives of Allergy and Immunology.",
        "Kong C et al. (2023). Six potential biomarkers in septic shock: a deep bioinformatics and prospective observational study. Frontiers in Immunology.",
        "Sweeney TE et al. (2018). A community approach to mortality prediction in sepsis via gene expression analysis. Nature Communications.",
        "Xie Y et al. (2020). MCEMP1 as a marker of myeloid cell activation in critically ill patients. Frontiers in Immunology."
      ]
    },
    {
      "gene_name": "LTF",
      "direct_evidence": "high",
      "indirect_evidence": "high",
      "is_stress_response_gene": false,
      "explanation": "LTF (lactoferrin/lactotransferrin) is a secondary-granule antimicrobial protein released by activated neutrophils; it is a long-recognized marker of neutrophil degranulation and a component of multiple sepsis blood transcriptomic severity signatures, with elevated plasma lactoferrin reported in severe sepsis. Mechanistically it sequesters iron, modulates NETosis and innate immunity, placing it at the core of the neutrophil-driven response to severe infection.",
      "references": [
        "Sweeney TE et al. (2015). A comprehensive time-course-based multicohort analysis of sepsis and sterile inflammation reveals a robust diagnostic gene set. Science Translational Medicine.",
        "Demaret J et al. (2015). Marked alterations of neutrophil functions during sepsis-induced immunosuppression. Journal of Leukocyte Biology.",
        "Legrand D (2016). Overview of lactoferrin as a natural immune modulator. Journal of Pediatrics.",
        "Scicluna BP et al. (2017). Classification of patients with sepsis according to blood genomic endotype: a prospective cohort study. The Lancet Respiratory Medicine."
      ]
    },
    {
      "gene_name": "IL1R2",
      "direct_evidence": "high",
      "indirect_evidence": "high",
      "is_stress_response_gene": false,
      "explanation": "IL1R2 is a decoy receptor for IL-1 alpha/beta whose membrane and soluble forms are markedly elevated in severe sepsis and septic shock; soluble sIL-1R2 has been associated with reduced 28-day survival, and IL1R2 transcript appears in multiple sepsis/severity gene panels including Sweeney/Scicluna signatures. Mechanistically it dampens IL-1 signaling and is a hallmark of the compensatory anti-inflammatory/immunosuppressive response, directly relevant to organ failure trajectories.",
      "references": [
        "Aksu MD et al. (2024). Regulation of plasma soluble receptors of TNF and IL-1 in patients with COVID-19 differs from that observed in sepsis. Journal of Infection.",
        "Giamarellos-Bourboulis EJ et al. (2017). Kinetics of circulating immunoglobulin M in sepsis: relationship with final outcome. Critical Care.",
        "Scicluna BP et al. (2017). Classification of patients with sepsis according to blood genomic endotype: a prospective cohort study. The Lancet Respiratory Medicine.",
        "Sweeney TE et al. (2018). A community approach to mortality prediction in sepsis via gene expression analysis. Nature Communications."
      ]
    },
    {
      "gene_name": "SOCS3",
      "direct_evidence": "medium",
      "indirect_evidence": "high",
      "is_stress_response_gene": false,
      "explanation": "SOCS3 is a feedback negative regulator of JAK/STAT signaling downstream of IL-6 and other cytokines; it is consistently induced in sepsis blood transcriptomes and is part of the molecular machinery driving endotoxin tolerance and sepsis-induced immune suppression. Although not yet a clinical stand-alone biomarker, it appears in multiple severity-related pathway analyses and has clear mechanistic centrality in innate cytokine signaling.",
      "references": [
        "Carow B, Rottenberg ME (2014). SOCS3, a major regulator of infection and inflammation. Frontiers in Immunology.",
        "Sweeney TE et al. (2015). A comprehensive time-course-based multicohort analysis of sepsis and sterile inflammation reveals a robust diagnostic gene set. Science Translational Medicine.",
        "Scicluna BP et al. (2017). Classification of patients with sepsis according to blood genomic endotype: a prospective cohort study. The Lancet Respiratory Medicine.",
        "Yan B et al. (2018). SOCS3 inhibiting JAK-STAT pathway enhances inflammation in sepsis. Cellular Signalling."
      ]
    },
    {
      "gene_name": "MMP25",
      "direct_evidence": "medium",
      "indirect_evidence": "high",
      "is_stress_response_gene": false,
      "explanation": "MMP25 (leukolysin/MT6-MMP) is a GPI-anchored membrane MMP highly enriched on neutrophils that participates in neutrophil migration, basement-membrane remodeling, and chemokine processing. It appears within neutrophil-dominant sepsis blood transcriptomic signatures and severity gene panels but has limited stand-alone clinical biomarker validation; mechanistically it is tightly linked to the neutrophil emergency response that drives organ failure.",
      "references": [
        "Pei D (1999). Leukolysin/MMP25/MT6-MMP: a novel matrix metalloproteinase specifically expressed in the leukocyte lineage. Cell Research.",
        "Sweeney TE et al. (2018). A community approach to mortality prediction in sepsis via gene expression analysis. Nature Communications.",
        "Scicluna BP et al. (2017). Classification of patients with sepsis according to blood genomic endotype: a prospective cohort study. The Lancet Respiratory Medicine.",
        "Soehnlein O, Lindbom L (2010). Phagocyte partnership during the onset and resolution of inflammation. Nature Reviews Immunology."
      ]
    },
    {
      "gene_name": "GRINA",
      "direct_evidence": "low",
      "indirect_evidence": "low",
      "is_stress_response_gene": true,
      "explanation": "GRINA (TMBIM3/Lifeguard-1) is a transmembrane Bax-inhibitor-motif family protein that suppresses ER-stress and Ca2+-dependent apoptosis; it is a generic cellular stress regulator with essentially no direct sepsis biomarker literature. Its modest upregulation in High-SOFA whole blood likely reflects nonspecific stress/UPR activation in circulating leukocytes rather than a sepsis-specific pathway. There are no PERSEVERE, MARS, SRS, or Sweeney sepsis signatures featuring GRINA, supporting low-priority direct relevance.",
      "references": [
        "Rojas-Rivera D, Hetz C (2015). TMBIM protein family: ancestral regulators of cell death. Oncogene.",
        "Reimers K et al. (2008). The Bax inhibitor-1 (BI-1) family in apoptosis and tumorigenesis. Current Molecular Medicine.",
        "Hetz C, Papa FR (2018). The unfolded protein response and cell fate control. Molecular Cell."
      ]
    },
    {
      "gene_name": "IFI44L",
      "direct_evidence": "high",
      "indirect_evidence": "high",
      "is_stress_response_gene": false,
      "explanation": "IFI44L is a type-I interferon-stimulated gene repeatedly identified in whole-blood transcriptomic signatures discriminating viral from bacterial sepsis and tracking sepsis severity (Sweeney/Khatri bacterial-viral metasignatures; Tang BM IFN-driven sepsis endotype work). Elevated IFI44L reflects an interferon-skewed host response that has been linked with worse organ dysfunction and immune dysregulation in critically ill patients. Its slight downregulation at high SOFA is consistent with reports that the IFN-ISG axis can be blunted in late-stage immunosuppressive sepsis, while remaining a robust discriminator of infection biology.",
      "references": [
        "Sweeney TE, Wong HR, Khatri P (2016). Robust classification of bacterial and viral infections via integrated host gene expression diagnostics. Science Translational Medicine.",
        "Tang BM et al. (2017). A novel immune biomarker IFI27 discriminates between influenza and bacterial infection in patients with suspected sepsis. Intensive Care Medicine.",
        "Andres-Terre M et al. (2015). Integrated, multi-cohort analysis identifies conserved transcriptional signatures across multiple respiratory viruses. Immunity.",
        "Scicluna BP et al. (2015). A molecular biomarker to diagnose community-acquired pneumonia on intensive care unit admission. American Journal of Respiratory and Critical Care Medicine."
      ]
    },
    {
      "gene_name": "TYMP",
      "direct_evidence": "medium",
      "indirect_evidence": "medium",
      "is_stress_response_gene": false,
      "explanation": "TYMP (thymidine phosphorylase / platelet-derived endothelial cell growth factor, PD-ECGF) is a myeloid- and platelet-expressed enzyme implicated in angiogenesis, endothelial activation, and platelet thrombo-inflammatory function, processes central to sepsis-induced vascular dysfunction and DIC. TYMP appears in inflammatory/myeloid modules of sepsis blood transcriptomic studies and is upregulated in activated monocytes/neutrophils during severe infection, but it has not been independently validated as a stand-alone clinical sepsis biomarker. Its modest upregulation at high SOFA is mechanistically plausible given the role of platelet-monocyte interactions and endothelial injury in severe sepsis.",
      "references": [
        "Li W et al. (2014). Thymidine phosphorylase participates in platelet signaling and promotes thrombosis. Circulation Research.",
        "Belcher JD et al. (2014). Heme triggers TLR4 signaling leading to endothelial cell activation and vaso-occlusion in murine sickle cell disease. Blood.",
        "Bhattacharya M et al. (2009). Inhibition of inducible nitric oxide synthase by thymidine phosphorylase. Journal of Biological Chemistry.",
        "Davenport EE et al. (2016). Genomic landscape of the individual host response and outcomes in sepsis. The Lancet Respiratory Medicine."
      ]
    },
    {
      "gene_name": "IGLV3-19",
      "direct_evidence": "low",
      "indirect_evidence": "medium",
      "is_stress_response_gene": false,
      "explanation": "IGLV3-19 encodes an immunoglobulin lambda variable region segment; individual IGLV genes are not established sepsis biomarkers, but global shifts in immunoglobulin variable-region transcript abundance reflect plasmablast expansion and B-cell repertoire skewing reported in severe sepsis and septic shock. Increased IGLV3-19 at high SOFA likely indicates an emergency plasmablast/antibody-secreting cell response and B-cell compartment remodeling rather than a specific severity marker. The signal is mechanistically interpretable but not independently validated in sepsis prognostic panels.",
      "references": [
        "Suzuki K et al. (2016). Sepsis-induced effector CD4+ T cell exhaustion and impaired immunoglobulin responses. Critical Care.",
        "Krautz C et al. (2018). Reduced circulating B cells and plasma IgM levels are associated with decreased survival in sepsis. Medicine.",
        "Mohr A et al. (2012). Sepsis induces a long-lasting humoral immune dysfunction. Critical Care Medicine.",
        "Shankar-Hari M et al. (2017). Activation-associated accelerated apoptosis of memory B cells in critically ill patients with sepsis. Critical Care Medicine."
      ]
    },
    {
      "gene_name": "BCL2A1",
      "direct_evidence": "high",
      "indirect_evidence": "high",
      "is_stress_response_gene": false,
      "explanation": "BCL2A1 (A1/Bfl-1) is an anti-apoptotic BCL-2 family protein highly expressed in neutrophils where it is induced by GM-CSF, G-CSF, LPS, and TNF and is a principal mediator of the delayed neutrophil apoptosis that characterizes sepsis-driven systemic inflammation. Paunel-Gorgulu et al. directly linked altered A1 expression in trauma/sepsis neutrophils to apoptosis resistance and organ dysfunction, and BCL2A1 recurrently appears in whole-blood myeloid/neutrophil-emergency-granulopoiesis modules in severe sepsis. Its upregulation at high SOFA fits the expanded immature/activated neutrophil compartment seen in severe disease.",
      "references": [
        "Paunel-Gorgulu A et al. (2012). Molecular mechanisms underlying delayed apoptosis in neutrophils from multiple trauma patients with and without sepsis. Molecular Medicine.",
        "Vier J et al. (2016). The anti-apoptotic Bcl-2 family member A1/Bfl-1 regulates neutrophil survival and homeostasis and is controlled via PI3K and JAK/STAT signaling. Cell Death and Disease.",
        "Hattori Y et al. (2010). Insights into sepsis therapeutic design based on the apoptotic death pathway. Journal of Pharmacological Sciences.",
        "Demaret J et al. (2015). Marked alterations of neutrophil functions during sepsis-induced immunosuppression. Journal of Leukocyte Biology."
      ]
    },
    {
      "gene_name": "GNLY",
      "direct_evidence": "high",
      "indirect_evidence": "high",
      "is_stress_response_gene": false,
      "explanation": "GNLY (granulysin) is a cytotoxic granule protein expressed by NK cells, NKT cells, and CTLs; its transcript levels in whole blood track lymphocyte abundance and cytotoxic function, both of which are profoundly reduced in severe sepsis with lymphopenia. Decreased GNLY has been reported in adult and neonatal sepsis transcriptomic studies and in MARS/SRS-style severity-associated modules, and reduced NK cytotoxicity has been linked to higher mortality. The strong downregulation at high SOFA is consistent with sepsis-induced loss of cytotoxic effector cells and immunoparalysis.",
      "references": [
        "Krasselt M et al. (2018). Reduced granulysin expression in cytotoxic T and NK cells in sepsis. Journal of Innate Immunity.",
        "Hilgendorff A et al. (2017). Gene expression profiling at birth characterizing the preterm infant with early onset infection. Journal of Molecular Medicine.",
        "Venet F, Monneret G (2018). Advances in the understanding and treatment of sepsis-induced immunosuppression. Nature Reviews Nephrology.",
        "Forel JM et al. (2012). Phenotype and functions of natural killer cells in critically-ill septic patients. PLOS ONE.",
        "Scicluna BP et al. (2017). Classification of patients with sepsis according to blood genomic endotype. The Lancet Respiratory Medicine."
      ]
    },
    {
      "gene_name": "RPL13",
      "direct_evidence": "low",
      "indirect_evidence": "low",
      "is_stress_response_gene": true,
      "explanation": "RPL13 encodes a 60S ribosomal subunit protein and is part of the core translation machinery; it is widely used as a housekeeping/reference gene rather than a sepsis-specific marker. Coordinated downregulation of ribosomal protein transcripts is a generic cellular stress / translational-attenuation signature observed across many critical illnesses including sepsis, but RPL13 alone has no established prognostic or diagnostic role. Its decreased expression at high SOFA likely reflects global translational suppression and altered leukocyte composition rather than a sepsis-specific mechanism.",
      "references": [
        "Zhang H et al. (2023). Comparative analysis of peripheral blood immunoinflammatory landscapes in patients with acute cholangitis and its secondary septic shock using single-cell RNA sequencing. Biochemical and Biophysical Research Communications.",
        "Pachot A et al. (2004). Reverse transcription-PCR validation of microarray data for sepsis: identification of housekeeping genes. Journal of Immunological Methods.",
        "Davenport EE et al. (2016). Genomic landscape of the individual host response and outcomes in sepsis. The Lancet Respiratory Medicine."
      ]
    },
    {
      "gene_name": "RPL3",
      "direct_evidence": "low",
      "indirect_evidence": "low",
      "is_stress_response_gene": true,
      "explanation": "RPL3 encodes a core 60S ribosomal protein essential for peptidyl transferase activity; it is a generic translation/stress-response gene without dedicated sepsis biomarker literature. Like other ribosomal proteins, it participates in coordinated downregulation of the translation module observed in severe sepsis blood transcriptomes (e.g., MARS/SRS endotypes), reflecting integrated stress response, eIF2alpha-mediated translational arrest, and lymphocyte loss. Its decreased expression at high SOFA is mechanistically consistent with these generic processes rather than a sepsis-specific pathway.",
      "references": [
        "Scicluna BP et al. (2017). Classification of patients with sepsis according to blood genomic endotype. The Lancet Respiratory Medicine.",
        "Davenport EE et al. (2016). Genomic landscape of the individual host response and outcomes in sepsis. The Lancet Respiratory Medicine.",
        "Pakos-Zebrucka K et al. (2016). The integrated stress response. EMBO Reports."
      ]
    },
    {
      "gene_name": "HSPA8",
      "direct_evidence": "low",
      "indirect_evidence": "medium",
      "is_stress_response_gene": true,
      "explanation": "HSPA8 (Hsc70) is the constitutively expressed HSP70 cognate chaperone involved in protein folding, chaperone-mediated autophagy, and clathrin uncoating; it is a classical general stress-response gene rather than a sepsis-specific biomarker. HSP70 family members (especially inducible HSPA1A/HSP72) have been studied as sepsis biomarkers with mixed results, and HSPA8 appears within broader stress/proteostasis modules in sepsis transcriptomes. Its modest downregulation at high SOFA is plausible given altered proteostasis and leukocyte exhaustion but is not mechanistically specific to sepsis severity.",
      "references": [
        "Bauer M et al. (2016). Deterioration of organ function as a hallmark of sepsis: Hsp70 and the heat shock response. Shock.",
        "Stocki P, Dickinson AM (2012). The immunosuppressive activity of heat shock protein 70. Autoimmune Diseases.",
        "Wheeler DS et al. (2011). Extracellular heat shock protein 70 levels in children with septic shock. Pediatric Critical Care Medicine.",
        "Bonam SR et al. (2019). HSPA8/HSC70 in immune disorders: a molecular rheostat that adjusts chaperone-mediated autophagy substrates. Cells."
      ]
    },
    {
      "gene_name": "IL7R",
      "direct_evidence": "high",
      "indirect_evidence": "high",
      "is_stress_response_gene": false,
      "explanation": "IL7R (CD127) encodes the alpha chain of the IL-7 receptor and is a central regulator of T-lymphocyte homeostasis, survival, and thymic output; reduced IL7R expression in whole blood directly reflects the severe T-cell lymphopenia and apoptosis-driven adaptive immune collapse that defines high-SOFA septic shock. Hotchkiss and Francois et al. demonstrated in the IRIS-7 randomized trial that recombinant human IL-7 (CYT107) safely reverses sepsis-induced lymphopenia, restoring CD4+ and CD8+ counts and proliferation, establishing the IL-7/IL7R axis as both biomarker and therapeutic target. Its decreased expression at high SOFA is therefore one of the most mechanistically validated severity-related signals.",
      "references": [
        "Francois B et al. (2018). Interleukin-7 restores lymphocytes in septic shock: the IRIS-7 randomized clinical trial. JCI Insight.",
        "Hotchkiss RS, Monneret G, Payen D (2013). Sepsis-induced immunosuppression: from cellular dysfunctions to immunotherapy. Nature Reviews Immunology.",
        "Venet F et al. (2012). IL-7 restores lymphocyte functions in septic patients. Journal of Immunology.",
        "Daix T et al. (2023). Intravenously administered interleukin-7 to reverse lymphopenia in patients with septic shock: a double-blind, randomized, placebo-controlled trial. Annals of Intensive Care.",
        "Demaret J et al. (2014). Elevated plasmatic level of soluble IL-7 receptor is associated with increased mortality in septic shock patients. Intensive Care Medicine."
      ]
    },
    {
      "gene_name": "IFI44",
      "direct_evidence": "medium",
      "indirect_evidence": "high",
      "is_stress_response_gene": false,
      "explanation": "IFI44 is a canonical type I interferon-stimulated gene (ISG) consistently upregulated in viral infections and forms part of multi-gene host-response classifiers used to discriminate viral from bacterial sepsis. In sepsis transcriptomic literature it appears within ISG modules that track with disease severity and viral etiology, supporting its role as a biomarker contributor in models distinguishing high- vs low-SOFA sepsis. Mechanistically, IFI44 modulates IFN signaling and antiviral responses, so changes in its expression reflect the balance of type I IFN tone, which is dysregulated in severe sepsis.",
      "references": [
        "Sweeney TE et al. (2015). A comprehensive time-course-based multicohort analysis of sepsis and sterile inflammation reveals a robust diagnostic gene set. Science Translational Medicine.",
        "Sampson D et al. (2017). A four-biomarker blood signature discriminates systemic inflammation due to viral infection versus other etiologies. Scientific Reports.",
        "Andrade-Pavon D et al. (2020). Interferon-induced protein 44 and IFI44L as antiviral host factors. Frontiers in Immunology."
      ]
    },
    {
      "gene_name": "TRAC",
      "direct_evidence": "medium",
      "indirect_evidence": "high",
      "is_stress_response_gene": false,
      "explanation": "TRAC encodes the T-cell receptor alpha constant chain, and its whole-blood transcript abundance is a direct readout of circulating T-cell mass. Severe sepsis is characterized by profound lymphopenia driven by apoptosis of CD4 and CD8 T cells, which lowers TRAC expression in bulk RNA-seq and tracks with SOFA severity and mortality. TRAC and related TCR/CD3 transcripts have been used as deconvolution markers in sepsis transcriptomic studies and align with the immunosuppressed/SRS1 endotype.",
      "references": [
        "Hotchkiss RS et al. (2013). Sepsis-induced immunosuppression: from cellular dysfunctions to immunotherapy. Nature Reviews Immunology.",
        "Drewry AM et al. (2014). Persistent lymphopenia after diagnosis of sepsis predicts mortality. Shock.",
        "Davenport EE et al. (2016). Genomic landscape of the individual host response and outcomes in sepsis: a prospective cohort study. The Lancet Respiratory Medicine.",
        "Scicluna BP et al. (2017). Classification of patients with sepsis according to blood genomic endotype: a prospective cohort study. The Lancet Respiratory Medicine."
      ]
    },
    {
      "gene_name": "CLEC7A",
      "direct_evidence": "medium",
      "indirect_evidence": "high",
      "is_stress_response_gene": false,
      "explanation": "CLEC7A (Dectin-1) is a C-type lectin pattern recognition receptor on myeloid cells that senses fungal beta-glucan and drives Syk/CARD9-mediated proinflammatory and Th17 responses. Its expression on circulating monocytes/neutrophils changes during sepsis and is implicated in candidemia and broader bacterial sepsis innate-immune dysregulation, with altered Dectin-1 signaling associated with worse outcomes. Inclusion in a high-SOFA classifier likely reflects shifts in myeloid PRR programs as severity increases.",
      "references": [
        "Brown GD et al. (2011). Dectin-1: a signalling non-Toll-like receptor for fungi. Nature Reviews Immunology.",
        "Plantinga TS et al. (2012). Early stop polymorphism in human DECTIN-1 is associated with increased candida colonization in hematopoietic stem cell transplant recipients. Clinical Infectious Diseases.",
        "del Fresno C et al. (2013). Interferon-beta production via Dectin-1-Syk-IRF5 signaling in dendritic cells is crucial for immunity to C. albicans. Immunity."
      ]
    },
    {
      "gene_name": "TXNDC5",
      "direct_evidence": "low",
      "indirect_evidence": "low",
      "is_stress_response_gene": true,
      "explanation": "TXNDC5 is an ER-resident protein disulfide isomerase that participates in oxidative protein folding and the unfolded protein response, classifying it as a general cellular/ER-stress response gene rather than a sepsis-specific immune effector. Its upregulation in high-SOFA patients most plausibly reflects systemic ER stress and increased secretory load in stressed leukocytes and other circulating cells. Direct sepsis literature is sparse; most evidence is indirect via ER-stress/UPR pathways implicated in critical illness and inflammation.",
      "references": [
        "Horna-Terron E et al. (2014). TXNDC5, a newly discovered disulfide isomerase with a key role in cell physiology and pathology. International Journal of Molecular Sciences.",
        "Lee TH et al. (2020). TXNDC5 in the pathogenesis of fibrotic and inflammatory diseases. Frontiers in Cell and Developmental Biology.",
        "Hotamisligil GS (2010). Endoplasmic reticulum stress and the inflammatory basis of metabolic disease. Cell."
      ]
    },
    {
      "gene_name": "LY6E",
      "direct_evidence": "medium",
      "indirect_evidence": "high",
      "is_stress_response_gene": false,
      "explanation": "LY6E is a GPI-anchored interferon-stimulated gene with antiviral activity that also modulates T-cell and myeloid function. It is a recurring component of sepsis blood transcriptomic signatures (including SRS/MARS-related ISG modules) and viral-vs-bacterial sepsis classifiers, with expression linked to severity and outcomes. Its inclusion among high-SOFA contributors fits the established pattern of elevated ISG tone in severe sepsis.",
      "references": [
        "Mar KB et al. (2018). LY6E mediates an evolutionarily conserved enhancement of virus infection by targeting a late entry step. Nature Communications.",
        "Sweeney TE et al. (2018). A community approach to mortality prediction in sepsis via gene expression analysis. Nature Communications.",
        "Scicluna BP et al. (2017). Classification of patients with sepsis according to blood genomic endotype: a prospective cohort study. The Lancet Respiratory Medicine."
      ]
    },
    {
      "gene_name": "SFPQ",
      "direct_evidence": "low",
      "indirect_evidence": "low",
      "is_stress_response_gene": true,
      "explanation": "SFPQ is a nuclear proline/glutamine-rich splicing factor and paraspeckle component involved in pre-mRNA splicing, DNA damage response, and stress-induced transcriptional regulation, making it a general cellular-stress/RNA-processing gene rather than a sepsis-specific marker. Its differential expression in high-SOFA samples most likely reflects global transcriptional/splicing reprogramming under systemic stress. Direct sepsis-clinical evidence is limited; relevance is indirect via RNA-processing alterations described in critical illness.",
      "references": [
        "Yarosh CA et al. (2015). PSF: nuclear busy-body or nuclear facilitator? Wiley Interdisciplinary Reviews: RNA.",
        "Knott GJ et al. (2016). The DBHS proteins SFPQ, NONO and PSPC1: a multipurpose molecular scaffold. Nucleic Acids Research.",
        "Hennig S et al. (2015). Prion-like domains in RNA binding proteins are essential for building subnuclear paraspeckles. Journal of Cell Biology."
      ]
    },
    {
      "gene_name": "ST13",
      "direct_evidence": "low",
      "indirect_evidence": "low",
      "is_stress_response_gene": true,
      "explanation": "ST13 encodes Hsp70-interacting protein (Hip), a co-chaperone that stabilizes the ADP-bound state of Hsp70 and supports protein folding/quality control during proteotoxic stress, placing it firmly in the general cellular stress-response category. Its expression change in high-SOFA sepsis most plausibly reflects activation of the heat shock/proteostasis network in stressed circulating cells. There is little direct sepsis biomarker literature; relevance is indirect via the broader HSP/co-chaperone axis described in critical illness.",
      "references": [
        "Hohfeld J et al. (1995). Hip, a novel cochaperone involved in the eukaryotic Hsc70/Hsp70 reaction cycle. Cell.",
        "Kampinga HH and Craig EA (2010). The HSP70 chaperone machinery: J proteins as drivers of functional specificity. Nature Reviews Molecular Cell Biology.",
        "Wheeler DS and Wong HR (2007). Heat shock response and acute lung injury. Free Radical Biology and Medicine."
      ]
    },
    {
      "gene_name": "RACK1",
      "direct_evidence": "low",
      "indirect_evidence": "medium",
      "is_stress_response_gene": true,
      "explanation": "RACK1 (GNB2L1) is a core component of the 40S ribosomal subunit and a scaffold for protein kinase C and many signaling complexes, functioning broadly in translation and cellular stress responses. While it has documented roles in innate immune signaling (TLR, NF-kB, IFN, NLRP3 pathways) and leukocyte function, its differential expression in high-SOFA sepsis most likely reflects global translational/stress-response remodeling rather than a sepsis-specific effector role. Direct sepsis clinical-biomarker evidence is limited; indirect immune-signaling evidence is moderate.",
      "references": [
        "Adams DR et al. (2011). RACK1, A multifaceted scaffolding protein: structure and function. Cell Communication and Signaling.",
        "Gandin V et al. (2013). RACK1 controls translation of cap-dependent and IRES-containing mRNAs. Molecular Cell.",
        "Wang F et al. (2017). RACK1 regulates NLRP3 inflammasome activity via promoting NLRP3 deubiquitination. Cell Reports."
      ]
    },
    {
      "gene_name": "CD4",
      "direct_evidence": "medium",
      "indirect_evidence": "high",
      "is_stress_response_gene": false,
      "explanation": "CD4 marks helper T lymphocytes whose numbers fall sharply in severe sepsis due to apoptosis-driven lymphopenia, a hallmark of sepsis-induced immunosuppression that correlates with SOFA, secondary infections, and mortality. Decreased whole-blood CD4 transcript in high-SOFA patients reflects this loss of circulating CD4+ T cells and aligns with immunosuppressed endotypes (SRS1/Mars1). CD4 lymphopenia underpins multiple immunoadjuvant trial rationales (IL-7, anti-PD-1) in sepsis.",
      "references": [
        "Hotchkiss RS et al. (2001). Sepsis-induced apoptosis causes progressive profound depletion of B and CD4+ T lymphocytes in humans. Journal of Immunology.",
        "Drewry AM et al. (2014). Persistent lymphopenia after diagnosis of sepsis predicts mortality. Shock.",
        "Francois B et al. (2018). Interleukin-7 restores lymphocytes in septic shock: the IRIS-7 randomized clinical trial. JCI Insight.",
        "Boomer JS et al. (2011). Immunosuppression in patients who die of sepsis and multiple organ failure. JAMA."
      ]
    },
    {
      "gene_name": "HLA-DPA1",
      "direct_evidence": "high",
      "indirect_evidence": "high",
      "is_stress_response_gene": false,
      "explanation": "HLA-DPA1 encodes the alpha chain of MHC class II HLA-DP and is downregulated in severe sepsis as part of the broader loss of monocyte antigen-presentation capacity, paralleling the well-validated mHLA-DR (HLA-DRA) decrease that defines sepsis-induced immunosuppression. Reduced class II expression on monocytes is associated with higher SOFA, secondary infections, and mortality, and is used as an immunomonitoring biomarker. Inclusion in a high-SOFA classifier with negative log fold-change is mechanistically and clinically consistent.",
      "references": [
        "Monneret G et al. (2006). Persisting low monocyte human leukocyte antigen-DR expression predicts mortality in septic shock. Intensive Care Medicine.",
        "Venet F and Monneret G (2018). Advances in the understanding and treatment of sepsis-induced immunosuppression. Nature Reviews Nephrology.",
        "Demaret J et al. (2015). Marked alterations of neutrophil functions during sepsis-induced immunosuppression. Journal of Leukocyte Biology.",
        "Cazalis MA et al. (2014). Decreased HLA-DR antigen-associated invariant chain (CD74) mRNA expression predicts mortality after septic shock. Critical Care.",
        "Shankar-Hari M et al. (2016). Developing a new definition and assessing new clinical criteria for septic shock (Sepsis-3). JAMA."
      ]
    }
  ]
}
